# Using structurally fungible biosensors to evolve improved alkaloid biosyntheses

**DOI:** 10.1101/2021.06.07.447399

**Authors:** Simon d’Oelsnitz, Wantae Kim, Nathaniel T. Burkholder, Kamyab Javanmardi, Ross Thyer, Yan Zhang, Hal Alper, Andrew D. Ellington

## Abstract

A key bottleneck in the microbial production of therapeutic plant metabolites is identifying enzymes that can greatly improve yield. The facile identification of genetically encoded biosensors can overcome this limitation and become part of a general method for engineering scaled production. We have developed a unique combined screening and selection approach that quickly refines the affinities and specificities of generalist transcription factors, and using RamR as a starting point we evolve highly specific (>100-fold preference) and sensitive (EC_50_ <30 μM) biosensors for the alkaloids tetrahydropapaverine, papaverine, glaucine, rotundine, and noscapine. High resolution structures reveal multiple evolutionary avenues for the fungible effector binding site, and the creation of new pockets for different chemical moieties. These sensors further enabled the evolution of a streamlined pathway for tetrahydropapaverine, an immediate precursor to four modern pharmaceuticals, collapsing multiple methylation steps into a single evolved enzyme. Our methods for evolving biosensors now enable the rapid engineering of pathways for therapeutic alkaloids.

## INTRODUCTION

Microbes have been extensively engineered for commercial-scale production of therapeutic plant metabolites, yielding many benefits over traditional plant cultivation methods, such as reduced water and land use, faster and more reliable production cycles, and higher purity of target metabolites. Microbial fermentation is currently used for the production of artemisinic acid, the immediate precursor to the antimalarial drug artemisinin, and in development for commercial production of cannabinoids, opiates, and tropane alkaloids^**1–5**^. However, scaling production typically requires several years and hundreds of person-years to complete^**6**^, and is largely bottlenecked by a reliance on low-throughput analytical methods for assessing strain and pathway performance^**7**^. We believe that prokaryotic transcriptional regulators can be readily repurposed as biosensors to directly report on compound production and pathway performance in living cells^**8,9**^, but because methods for generating specific biosensors are lacking there are virtually no extant biosensors for most plant metabolites. Directed evolution is potentially a starting point for the generation of new biosensor specificities, but to date has proven quite limited, yielding improvements in responsiveness only to known effectors or close analogs thereof^**10–13**^.

To overcome this limitation, we looked to exploit a key insight from natural selection, that a protein’s substrate promiscuity correlates with its evolvability^**14**^. Thus, by starting with biosensors that are broadly represented in phylogeny, and whose substrate specificities have already been shown to be fungible in terms of natural ligands, it should be possible to create biosensors for virtually any compound. In particular, prokaryotic multidrug resistance regulators, typically studied as mediators of broad-spectrum antibiotic resistance, have large substrate binding pockets and are known to recognize a raft of structurally-diverse lipophilic molecules via non-specific interactions^**15**^. Early studies suggest that they may also be highly evolvable; notably, just a single point mutation enabled one of these regulators, TtgR, to adopt substantial affinity for the non-cognate ligand resveratrol^**16**^.

Using a novel directed evolution circuit architecture that relies on both screening and selection, we can seamlessly filter sensor libraries of over 10^5^ members into just a few high performing variants in under one week. As proof, we start with a single multidrug resistance regulator, RamR from *Salmonella typhimurium*, and evolve it to sensitively and specifically recognize five diverse therapeutic alkaloids. The high resolution structure of these sensors reveal how the malleable effector binding site can learn to specifically interact with entirely new ligands in wildly different ways. Ultimately, to demonstrate the utility of these sensors as a tool for metabolic engineering, we apply one sensor to engineer a multifunctional plant alkaloid methyltransferase capable of biosynthesizing tetrahydropapaverine, an immediate precursor to four modern pharmaceuticals.

## RESULTS

### Identifying a BIA-responsive multidrug resistance regulator

We have focused on generating sensors for benzylisoquinoline alkaloids (BIAs) since they (1) are rich in therapeutic activity, (2) have largely resolved biosynthetic pathways, and (3) are the subject of ongoing academic and commercial efforts^**3,4**^. We reasoned that the lipophilic nature of alkaloids might lead multidrug resistance regulators to display a basal affinity for these compounds. Therefore, we initially targeted five structurally diverse BIAs tetrahydropapaverine (THP), papaverine (PAP), rotundine (ROTU), glaucine (GLAU), and noscapine (NOS). These compounds are all therapeutically relevant, commercially available, and belong to the structurally distinct benzylisoquinoline (THP and PAP), protoberberine, aporphine, and phthalideisoquinoline BIA families, respectively (**Fig. 1a** and **Supplementary Fig. 1**). Furthermore, the complete microbial biosynthesis of noscapine and rotundine have recently been reported^**17,18**^.

**Figure 1.**
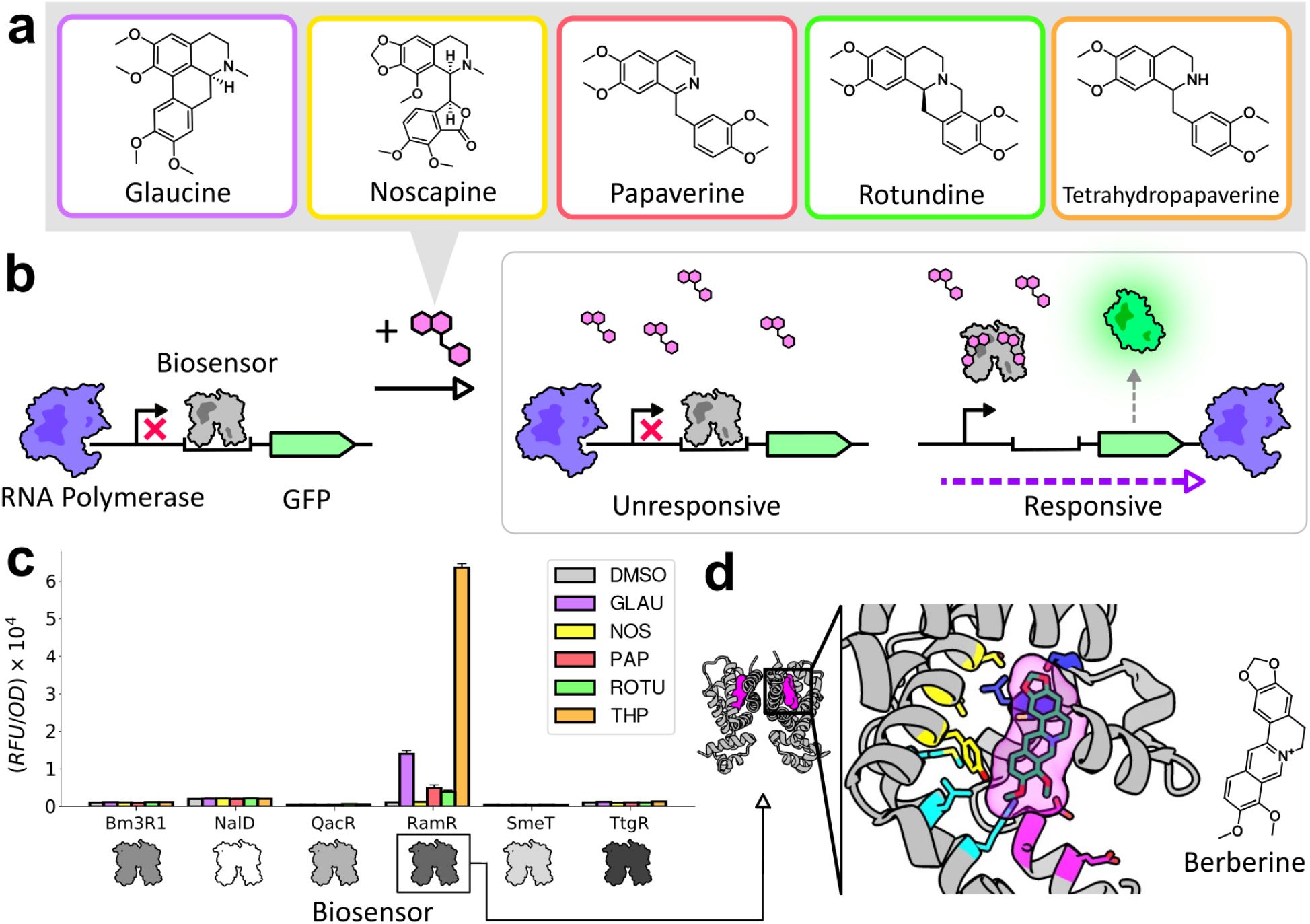
Screening identifies a biosensor responsive to benzylisoquinoline alkaloids (BIAs) (**a**) Chemical structures of the five BIAs used in the screen. (**b**) Schematic of the genetic circuit used for screening the responsiveness of candidate sensors to target BIAs. (**c**) Fluorescence response of six biosensors to all five BIAs. Ligand concentrations used for induction are indicated as follows. Glaucine: 1 mM, noscapine: 100 μM, papaverine: 500 μM, rotundine: 250 μM, tetrahydropapaverine: 1 mM. Fluorescence values are the averages of three biological replicates (**d**) The global structure (left) and ligand binding pocket (right) of RamR in complex with berberine (PDB: 3VW2). Colored residues were targeted for mutagenesis.

To identify a template biosensor with some degree of BIA affinity, we assayed the responsiveness of six well-characterized multidrug resistance regulators, QacR, TtgR, RamR, SmeT, NalD, and Bm3R1, to the target BIAs. Regulators were constitutively expressed on one plasmid (pReg) that was co-transformed with another plasmid bearing the regulator’s cognate promoter upstream of sfGFP (pGFP). Promoters for QacR and TtgR were obtained from the literature^**16,19**^ while promoters for the remainder were designed by either placing the sensor’s operator downstream a medium strength promoter (Bm3R1) or by modifying the −35 or −10 regions of the sensor’s native promoter towards the *E. coli* consensus (NalD, SmeT, RamR)^**20,21**^ (**Supplementary Fig. 2**). This design strategy was surprisingly successful, as each regulator could readily repress transcription, as measured via fluorescence (**Supplementary Fig. 2**). This also allowed further screening with otherwise unknown effectors for the semi-specific transcription factors, and RamR from *S. typhimurium* was in fact found to be moderately responsive to many target BIAs (**Fig. 1c**). In addition, the structure of RamR had already been solved in complex with berberine (PDB: 3VW2), an alkaloid related to our target ligands^**22**^. These conjoined informatics and experimental efforts thus quickly led to a quite rational approach to library design for directed evolution to improve affinity: five libraries encompassing five separate helices facing the ligand binding pocket were created in which three residues in each library were site-saturated (**Fig. 1d**). Each library contained ~32,000 unique genotypes and three to four libraries were pooled prior to selection, meaning that ~100,000 - 130,000 unique genotypes were assessed per round of evolution. In addition, error-prone libraries of the entire coding sequence were generated that had an average of two mutations per gene.

### Circuit design for biosensor evolution

While biosensors are typically evolved by screening sensor libraries via fluorescence activated cell sorting (FACS) in the absence and presence of the target ligand^**10–13**^, this approach generally yields sensors with a high background signal, likely since the most commonly identified mutations and phenotypes shift the allosteric equilibrium towards the ‘on’ state, generally increasing activity both with and without effector. Enrichment for strongly repressing sensor variants can be difficult via sorting due to inadequate filtering resolution and false positives that arise from dead or inactive cells. Selections, however, enable higher resolution filtering by enriching strongly repressing sensor variants in live, active cells through exponential growth. In turn, simpler fluorescence-based screens are already well-suited to distinguish highly responsive sensors. Given these considerations, we designed a new directed evolution circuit architecture, specifically tailored for sensor evolution, that leverages the advantages of both selections and screens: Seamless Enrichment of Ligand Inducible Sensors (SELIS).

To selectively remove sensors with a reduced ability to repress transcription in the absence of the target ligand and variants that were responsive to non-target ligands, we implemented an inverter circuit involving the Lambda cI repressor that would lead to expression of the zeocin resistance protein encoded by the *Sh ble* gene only in the absence of ligand (**Fig. 2b**). *Sh ble* was chosen for its non-catalytic mechanism of action, enabling a more linear application of selection stringency than would be the case for other antibiotic resistance elements^**23**^. Trial selections indeed showed enrichment for functionally repressing RamR variants in a zeocin-dependent manner (**Supplementary Fig. 3**).

**Figure 2.**
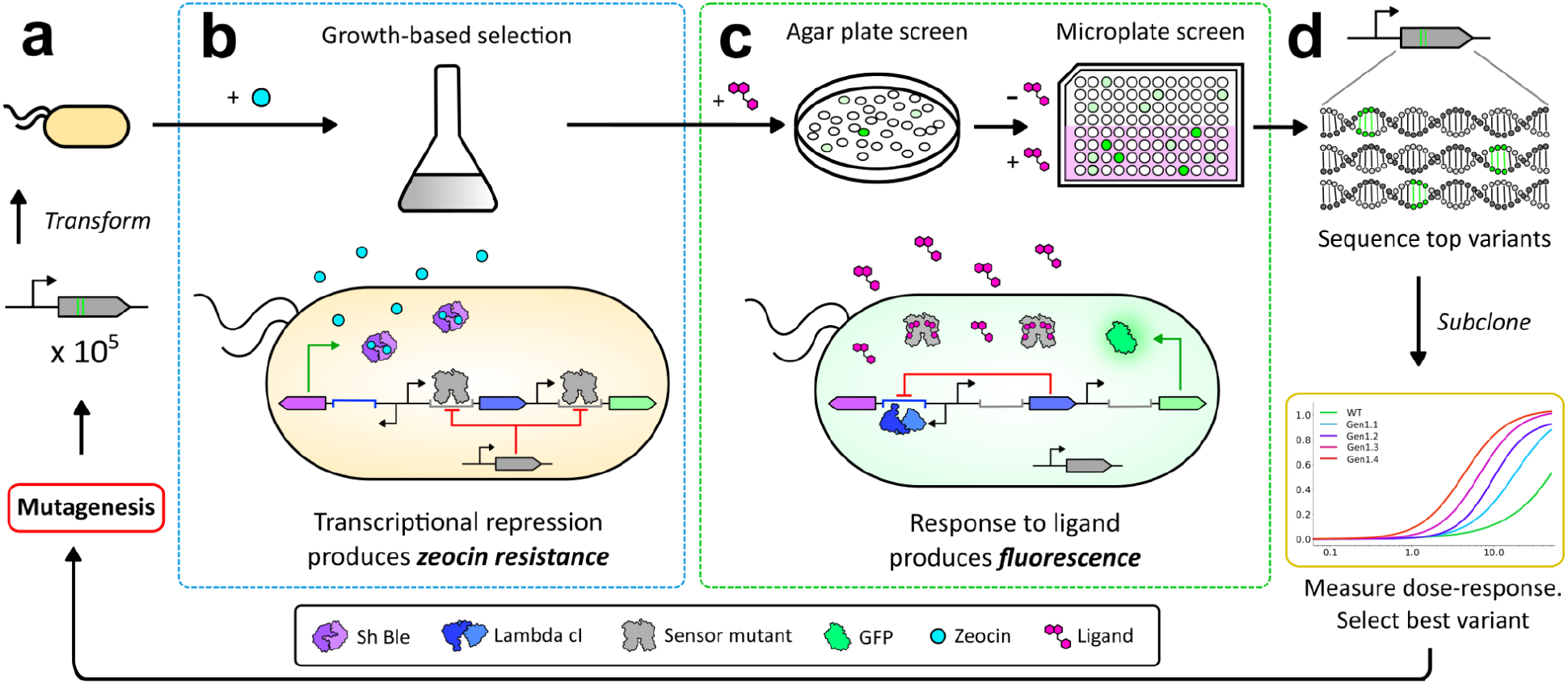
The SELIS (Seamless Enrichment of Ligand-Inducible Sensors) approach for biosensor evolution. (**a**) Libraries are generated and transformed into *E. coli* cells (**b**) Cells containing the sensor library are cultured in the presence of zeocin. Transcriptional repression by sensor variants prevents the expression of Lambda cI, which enables the expression of Sh Ble and confers zeocin resistance. Cells containing sensor variants that are unable to repress are eliminated from the population. Adding non-target ligands at this stage enables counter-selection for specificity. (**c**) Binding of the sensor variant to the target ligand relieves repression of GFP expression, producing fluorescence. Cultures are plated on an LB agar plate containing the target ligand and highly fluorescent colonies are cultured overnight. Subsequently, cultures from each picked colony are split and grown either with or without the target ligand. (**d**) Variants that display high signal/noise ratios are sequenced, subcloned, and re-phenotyped with a wider range of ligand concentrations. The top performing variant is then used for the next cycle of evolution.

Following this selection, we screened for variants that were more responsive to the target ligand by linking sensor output to the expression of GFP (**Fig. 2c**). Liquid cultures grown in the presence of zeocin (subjected to the negative selection) were plated directly onto solid media containing the target ligand. Highly fluorescent clones were isolated and re-phenotyped in liquid medium in both the presence and absence of the target ligand to better determine the signal/noise ratio for each chosen sensor variant. The stringency of the positive screen could be readily tuned by altering the amount of the target ligand applied on the solid media.

Ultimately, variants with low background and a high signal/noise ratio were sequenced and unique variants more fully characterized using a wider range of ligand concentrations (**Fig. 2d**). The highest performing biosensor variant was then used as a template for the next round of selection and screening. A library containing ~10^5^ variants can be deconvoluted to yield phenotype and genotype data for high performing clones in just one week, without the need for specialized equipment. The SELIS methodology should be broadly applicable to evolve virtually any prokaryotic ligand-inducible transcription factor.

### Evolving RamR specificity towards benzylisoquinoline alkaloids

While multidrug resistance regulators are known to recognize structurally diverse ligands, the ability to hone their inherently broad effector specificity has been rarely explored^**16**^. In order to demonstrate the utility and speed of SELIS, wild-type RamR was used as a starting point and four rounds of evolution were performed for the five BIAs (tetrahydropapaverine (THP), papaverine (PAP), rotundine (ROTU), glaucine (GLAU), and noscapine (NOS)) for a total of 20 RamR sensor generations. As library positions became fixed, new site-saturation libraries were included at non-fixed positions to reintroduce diversity (**Supplementary Fig. 4**). Following the first round of directed evolution, the strength of the promoter expressing the RamR variant and the concentration of the target BIA were reduced to increase the selection stringency for repression and ligand responsiveness, respectively (**Supplementary Table 1**). A key feature of SELIS is that non-target ligands can be added at this stage to counterselect against non-specific sensors. After the second round of evolution, 100 μM of non-target BIAs were added during the growth-based selection to eliminate less specific sensor variants.

Over the course of four generations of evolution, discrete evolutionary lineages became highly sensitive to their cognate BIA (**Supplementary Figs. 5-9**). Despite having a barely detectable response to most target BIAs initially, four of the five final RamR variants had an EC_50_ value under 7 μM, highlighting the plasticity of this biosensor scaffold (**Fig. 3a–e**). Notably, the detectable concentration range for the final noscapine biosensor is well within the level that is reported to be produced *de novo* in yeast^**17**^. In addition, the background signal was also reduced to less than 40% of wild-type RamR for four of the five final biosensors (**Fig. 3f–j**). Low background signals generally correlated with an increased signal-to-noise ratio and better limit of detection for the ligand.

**Figure 3.**
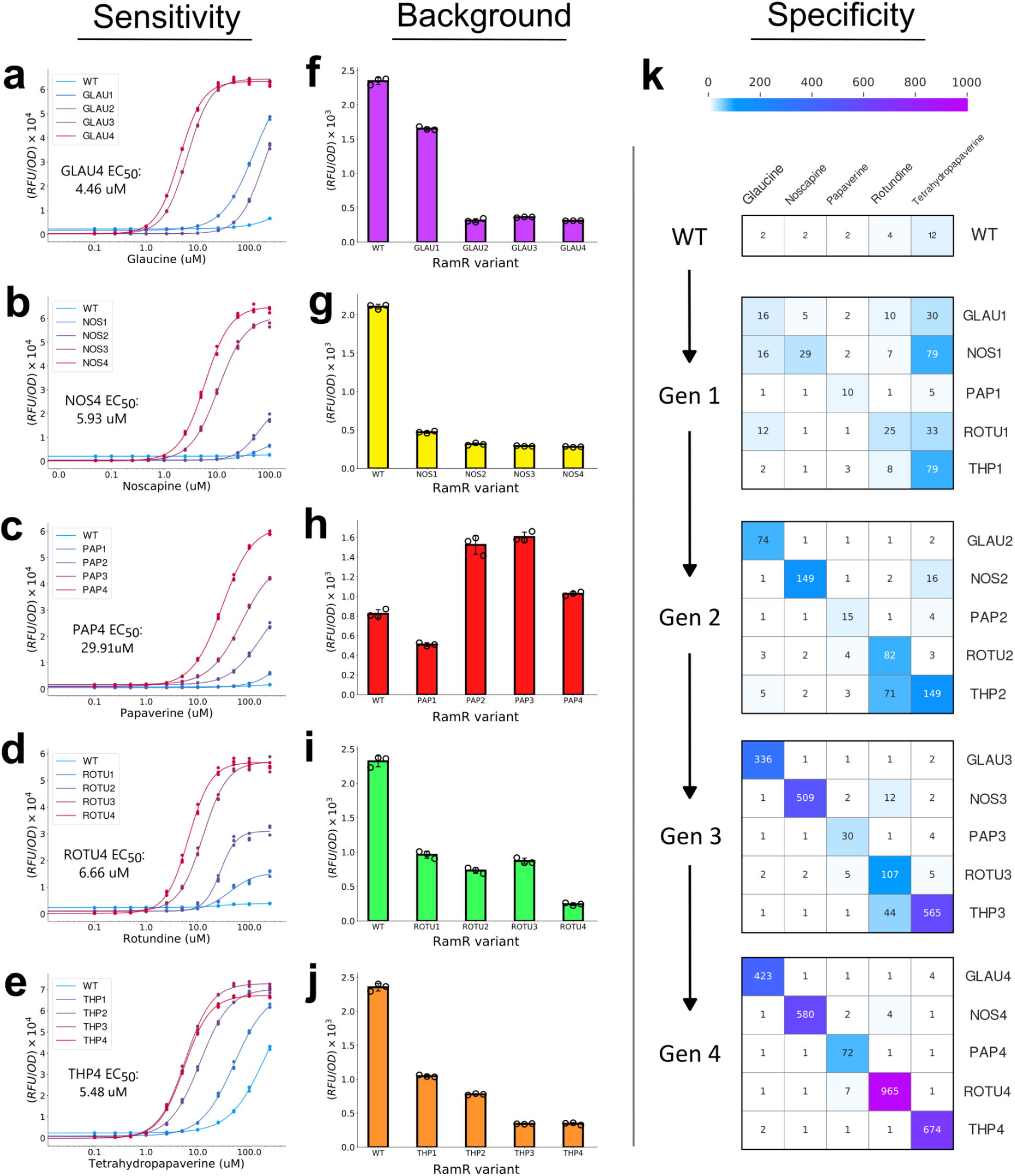
Evolution of highly specific BIA sensors from a generalist template (**a-e**) Transfer functions for all four generations of RamR variants with five different BIAs. The maximum ligand concentration was chosen based on the compound’s solubility limit in 1% DMSO (100 μM for Noscapine and 250 μM for all other BIAs). Fluorescent measurements for each condition were an average of three biological replicates. (**f-j**) Background fluorescence measurements for all RamR generations. The same promoter was used to express variants from each evolutionary trajectory (*see methods*) (**k**) Orthogonality matrix of all evolved sensors. Fold-response is shown for all BIAs for the native RamR protein, the first, second, third, and fourth generations from top to bottom, respectively. 100 μM of the indicated BIA was applied in all conditions. Measurements for each condition were an average of three biological replicates.

What was most surprising was that despite starting from the same generalist template all five final biosensor variants proved to be extremely specific for their matching BIA, displaying >100-fold preference for their cognate BIA over all other non-cognate BIAs at solubility-limiting concentrations (100 μM) of each compound (**Fig. 3k** and **SI Fig. 10**). High specificity is crucial for sensors used in strain engineering to avoid false positives arising from cross-reactivity with non-cognate ligands, particularly biosynthetic precursors.

### Structures reveal shared and unique adaptations to diverse alkaloids

Since both the ligand sensitivity and specificity of RamR could be dramatically transformed throughout evolution, with sensors accumulating from nine to thirteen mutations, we sought to better understand the molecular adaptations by solving the crystal structures of the best evolved sensors. We solved the structures of four of the five evolved sensors in complex with their cognate BIA: PAP4 with papaverine (1.6 Å), ROTU4 with rotundine (1.8 Å), GLAU4 with glaucine (2.0 Å), and NOS4 with noscapine (2.2 Å) (**Supplementary Table 2**). The overall folding and dimerization of the evolved variants is highly identical to that of wild-type RamR (**Fig. 4a**). A strong positive electron density was consistently detected at the binding site for each molecule in the asymmetric unit, which perfectly fit with the BIA chemical structures. (**Fig. 4b** and **SI Fig. 11**).

**Figure 4.**
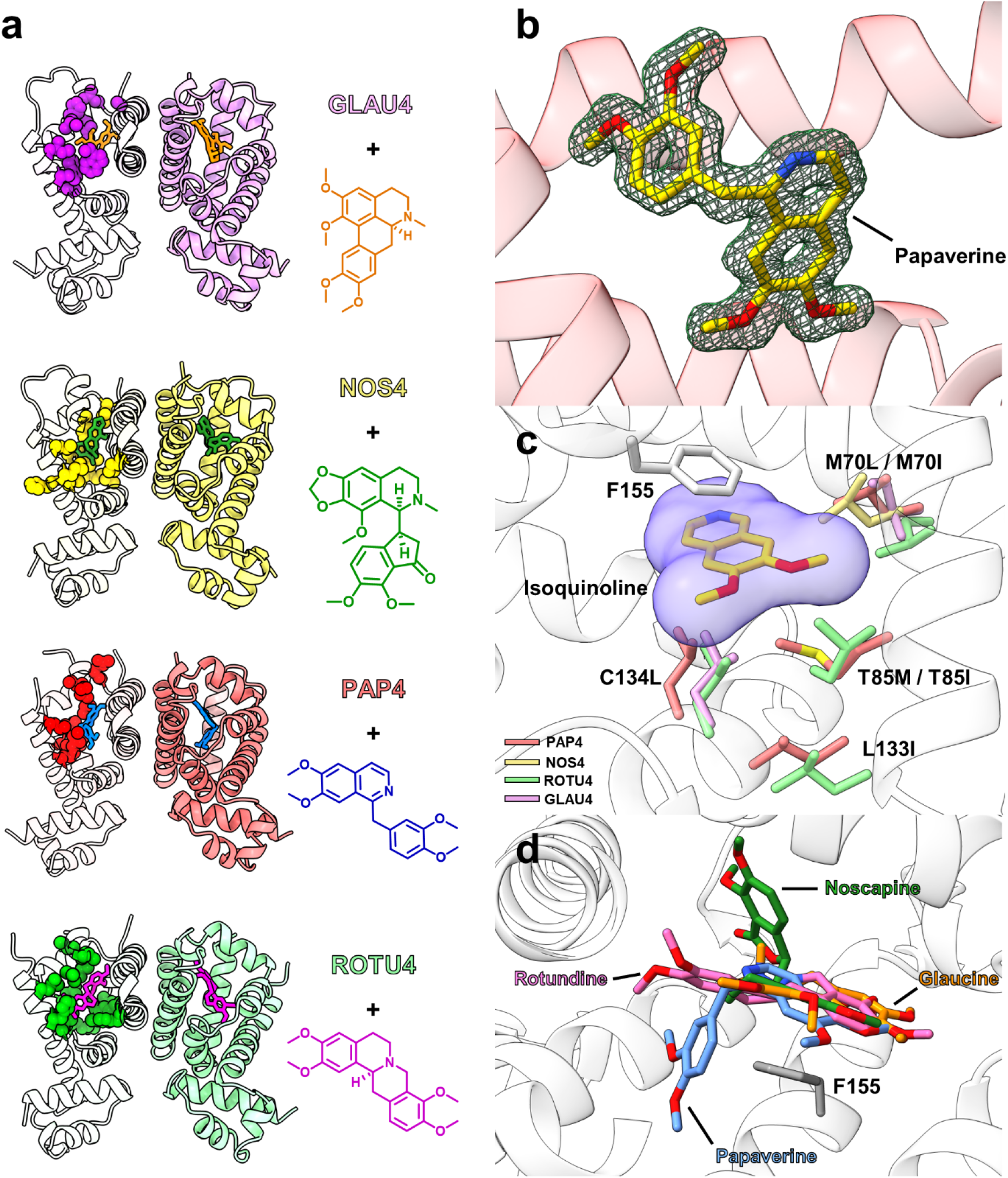
Crystal structures of evolved biosensors bound to cognate benzylisoquinoline alkaloids. (**a**) Overall structures of the four evolved RamR variants in ribbon diagram. The specific ligand for each variant is shown in stick with the binding site for one of the monomer shown in space-filling model to highlight the binding pockets.(**b**) Omit Fo-Fc map (contoured at 3.0σ) shown as a green mesh superimposed on the stick model of papaverine molecule (carbon atoms in yellow, oxygen atoms in red, and nitrogen atom in blue). (**c**) Superimposed structures of the complexes with the side chains of residues 70, 85, 133, and 134 in stick with color scheme as PAP4 (salmon), NOS4 (yellow), ROTU4 (green), and GLAU4 (violet). The isoquinoline ring part of all four ligands (yellow stick) are shown as space-filling with isoquinoline shown in stick (color scheme identical as b). The side chain of F155 π-π stacking with the isoquinoline ring is shown in stick and colored gray. (**d**) Superimposed BIA structures from all BIA-specific RamR variants shown relative to each other and the F155 residue of RamR (ligand color scheme identical as a).

BIAs contain heterocycle isoquinoline and benzyl group moieties, with a wide variety of intervening connectivities distinguishing individual compounds. The configuration of each ligand complexed with RamR variants reveals that the isoquinoline component is always ‘fixed’ beneath Phe155 with π - π stacking interactions, with further recognition by a hydrophobic pocket created by mutations in residues 70, 85, 133, and 134 (**Fig. 4c**). For example, C134 is consistently changed to leucine to form a hydrophobic interaction with the isoquinoline ring. Similarly, M70 frequently mutates into a shorter hydrophobic residue (leucine or isoleucine) to reinforce hydrophobic interactions with the BIA ligand. Mutations are not restricted to direct contact with the ligand: the L133I substitution supports other changes at position 85 (PAP4: T85M / L133I; ROTU4: T85I / L133I), making room for bulkier mutations at T85 with higher hydrophobicity. Identification of this common binding pattern and key residues involved in BIA recognition may facilitate structure-guided engineering of further sensors for morphinans and other therapeutic alkaloids.

Most surprisingly, despite the structural similarities among BIA ligands, each BIA biosensor employs unique mechanisms to accommodate heteroatoms and the benzyl ring moiety. This is easy to see given that the binding sites for the benzyl moieties of papaverine and noscapine extend in entirely different directions (**Fig. 4d**) and occupy different, newly evolved pockets. Examining the unique benzyl pockets in greater detail, noscapine extends into a side pocket for its specificity. The H135Y substitution assists the accommodation of dimethoxybenzyl moiety by forming pseudo π - π interaction and participating in the hydrogen bond network associated with the ester group of noscapine, while the mutation of E120 and D124 into highly flexible arginine residues creates an electrophilic network with H135Y and D152 that form favorable hydrophilic interactions (**Fig. 5a**). Unique accommodations are observed for the benzyl moieties of other ligands, as well. A Y92G mutation opens a new binding cavity in PAP4 allowing occupancy by the dimethoxybenzyl group of papaverine (**Fig. 5b**), while K63Y and L156Y mutations in ROTU4, along with Y92, form a triple-tyrosine ‘hydrophobic cage’ that traps the dimethoxybenzyl group of rotundine (**Fig. 5c**), and the L66W and Y92W substitutions in GLAU4 create a large tryptophan sandwich motif that pin the hydrophobic glaucine fused rings (**Fig. 5d**). Similarly, the nitrogen atom of papaverine is coordinated by the K63R substitution in PAP4 (strongly anchored by the adjacent A123D substitution (**Fig. 5b**)), while in ROTU4 the K63Y and L156Y substitutions coordinate two ordered water molecules to interact with the nitrogen atom of rotundine (**Fig. 5c**), and the native D152 residue of GLAU4 interacts with glaucine’s nitrogen atom (**Fig. 5d**). Indeed, though all alkaloids exhibit similar orientation towards the nitrogen atom (**Supplementary Fig. 12**), each RamR variant employs a unique adaptation to stabilize it (**Fig. 5a–d**). These structural data highlight the inherent plasticity of the RamR protein to rapidly evolve new ligand specificity and invent completely new modes of binding, suggesting that it may be a “privileged template” for biosensor engineering.

**Figure 5.**
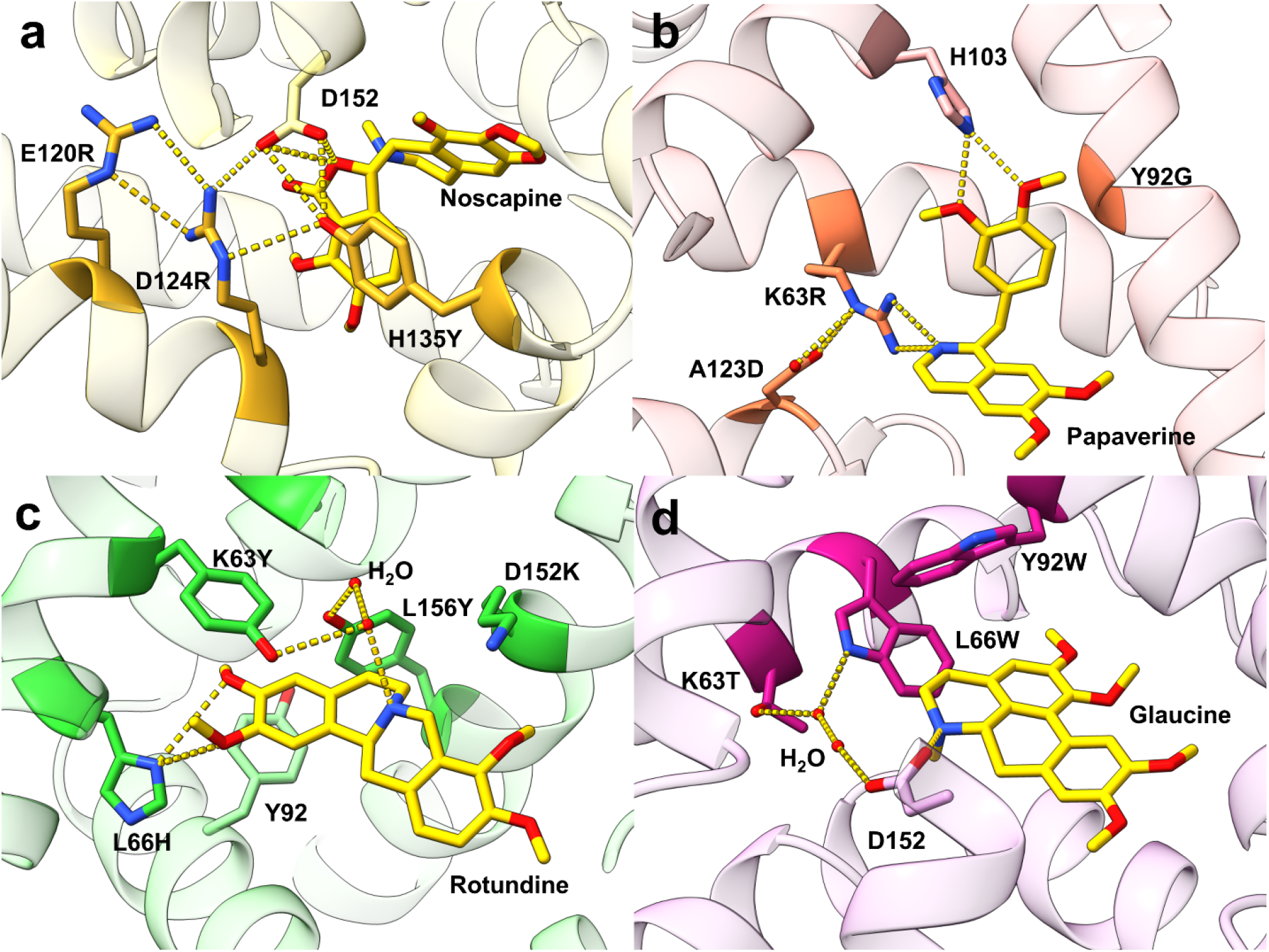
Unique molecular adaptations confer alkaloid specificity. (**a-d**) Structure of evolved sensors in complex with their cognate BIAs (shown in stick with carbon atoms colored yellow, oxygen atoms in red and nitrogen in blue). Residues involved in specific interactions with the cognate ligand are displayed in stick and labeled with color scheme as carbon atoms in yellow, oxygen atoms in red, and nitrogen atoms in blue.

### Evolved biosensors enable a new tetrahydropapaverine biosynthetic pathway

Numerous BIAs have been recognized for their therapeutic value as modern pharmaceuticals. Specifically tetrahydropapaverine (THP), which is naturally produced in plants, is used as an immediate precursor to four FDA-approved drugs: atracurium, cisatracurium, mivacurium, and papaverine^**24**^. In plants, THP is thought to be produced from the common intermediate norcoclaurine via an oxidase and four separate O-methyltransferases^**25**^. Since some BIA methyltransferases have promiscuous activities^**26**^ we reasoned that it might be possible to use our biosensor to evolve a methyltransferase that could uniquely methylate all four positions, thereby reducing a four enzyme pathway into a single multifunctional enzyme.

To create an abbreviated THP biosynthetic circuit suitable for screening and eventually directed evolution, we sought to identify an optimal ‘one-step’ enzyme by screening several promiscuous methyltransferases. To aid this effort, as well as subsequent evolution, we developed and tuned a THP reporter plasmid (pThpR) (**Supplementary Fig. 13**) that relies on the most sensitive THP biosensor (THP4.2), with an EC_50_ under 2uM (**Supplementary Fig. 9**). Since THP4.2 was somewhat responsive to a semi-methylated intermediate (norreticuline) but not the unmethylated substrate, this circuit should report on general methylation activity, likely favoring more completely methylated derivatives (**Supplementary Fig. 13**). To screen methyltransferase variants, *E. coli* cells co-transformed with pThpR and a plasmid expressing the methyltransferase were cultured in the presence of the substrate norlaudanosoline (NOR) for 24 hours and fluorescence was subsequently measured. Upon screening, one methyltransferase, GfOMT1 from *Glaucium flavum*, produced the highest fluorescent signal and was chosen as the starting point for evolution (**Supplementary Fig. 14**).

Error-prone libraries of GfOMT1 with an average of two mutations per gene were co-transformed with pThpR and highly fluorescent colonies were screened on solid media supplemented with NOR (**Fig. 6a)**. The resulting enzyme variants were sub-cloned and re-phenotyped (**Supplementary Fig. 15**), and the best performing variant was then used as the template for the next round of directed evolution. Over five rounds of screening an OMT variant with seven substitutions (GEN5) produced a 6- and 47-fold increase in fluorescent signal using pThpR when cultured with 100 μM or 10 μM of NOR, respectively, when compared to wild-type GfOMT1 (**Fig. 6b–c**).

**Figure 6.**
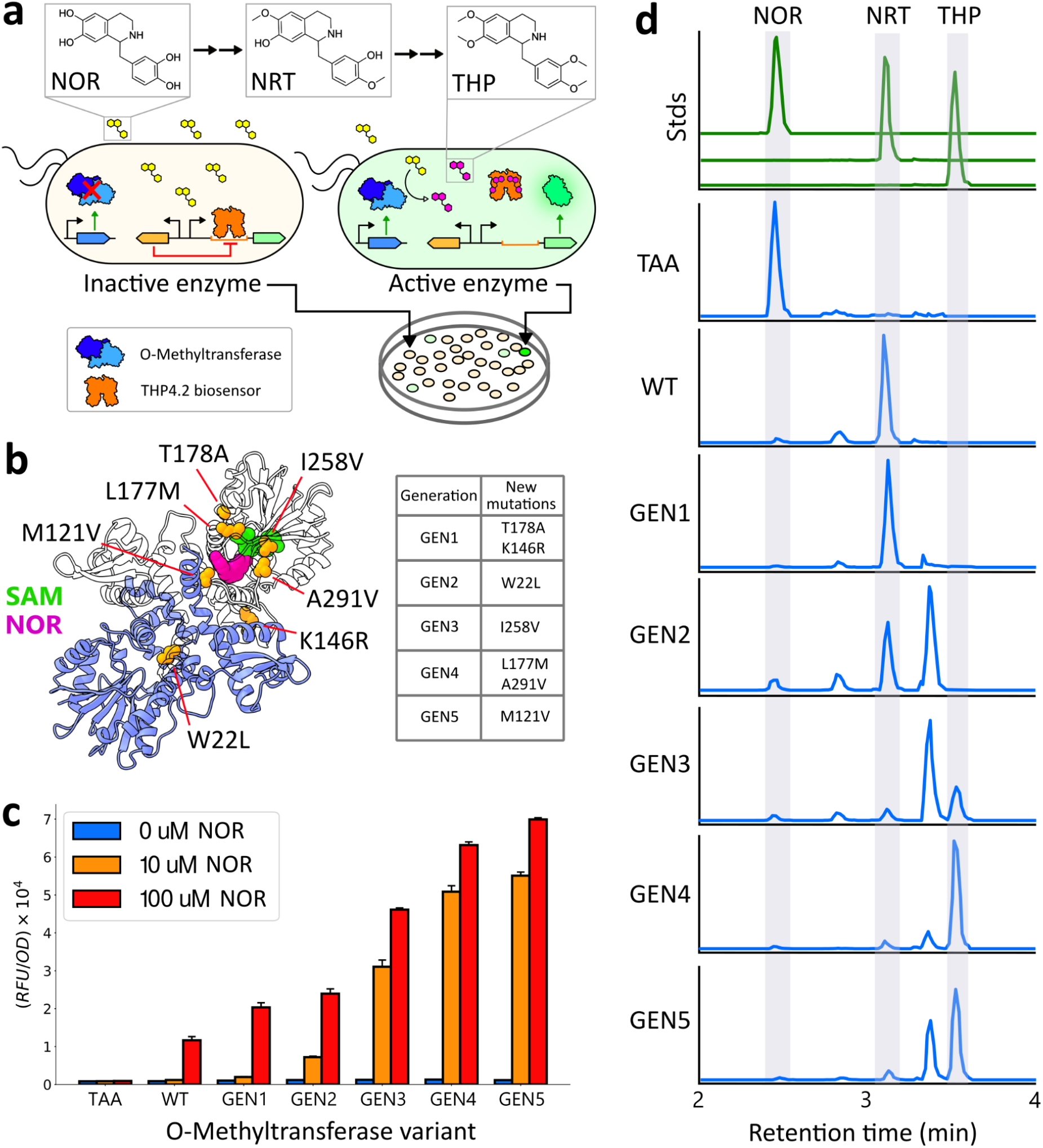
Evolved biosensor enables a new tetrahydropapaverine biosynthetic pathway **(a**) Genetic circuit for O-methyltransferase (OMT) evolution. A plasmid expressing the OMT variant (blue) is co-transformed with a plasmid expressing GFP regulated by a tetrahydropapaverine-responsive biosensor (THP4.2 in orange). OMT libraries are plated with norlaudanosoline (NOR) and highly fluorescent colonies are picked, characterized, and the top variant is used for the following round of evolution. (**b**) Homology structure of the template OMT (GfOMT1) with mutations in evolved variants shown in orange and labelled. The substrate NOR and cofactor S-adenosyl-methionine are shown in pink and green, respectively. (**c**) Fluorescence response of the THP4.2 reporter plasmid cultured with either 0 μM, 10 μM, or 100 μM of NOR and an empty plasmid (TAA), GfOMT1 (WT), or evolved OMT variants. (**d**) Representative ion extracted chromatograms of strains expressing engineered OMT variants, or controls, grown in the presence of 10 μM NOR. All LC-MS chromatograms were selected for the theoretical *m/z* values and retention times of the respective compounds of interest (**Supplementary Table 3**). A background peak for THP was subtracted; full chromatograms can be found in **Supplementary Fig. 18**.

Throughout OMT evolution, later generations successively methylate more positions on NOR **(Fig. 6d)**. Notably, the W22L mutation that occurred in GEN2 is thought to expand the substrate binding pocket and enable production of the trimethylated intermediate whereas the I258V mutation from GEN3 is adjacent to the expected S-adenosyl-methionine binding site and enables complete methylation to THP (**Supplementary Fig. 16**). The GEN3, GEN4, and GEN5 variants proved fully capable of making THP, with a conversion efficiency of 6%, 43%, and 26%, respectively **(Supplementary Fig. 17**). Interestingly, the GEN4 variant produced THP more selectively and efficiently than GEN5, despite both producing a similar fluorescent response with the pThpR reporter (**Fig. 6c)**.

## DISCUSSION

Using a directed evolution circuit architecture, Seamless Enrichment of Ligand-Inducible Sensors (SELIS), we demonstrate that the TetR family member RamR can be readily adapted to specifically and sensitively recognize therapeutic alkaloids for which no other extant biosensors exist. Even though the wild-type sensor only poorly recognized a small subset of the target alkaloids, we were able to evolve biosensors to a wide variety of structures, most of which had EC_50_ values under 7 μM and >100-fold specificities for their cognate ligands, making them practical for screening even low-flux pathways during strain development.

While previous work has largely relied on identifying specialist biosensors that recognize ligands closely related to a target^**10–13**^, we leverage insights from natural selection: start with a generalist, and evolve to a specialist^**14**^. This approach not only expands the chemical search space, but also accelerates protein evolution by circumventing the exploration of generalist intermediates on the way to new specializations^**27,28**^, an approach that is similar to the directed evolution of new specificities for highly promiscuous enzymes^**29,30**^. Pairing this generalist-first approach with our novel selection-screening circuit architecture and focused mutagenesis libraries afforded massive improvements in biosensor performance with otherwise structurally diverse alkaloids (for example, the response to 100 μM noscapine jumped from 2-fold to 509-fold after just three rounds of evolution (**Fig. 3k**)). Structural data for evolved RamR variants reveal that the RamR effector binding site is extraordinarily malleable. Directed evolution of the generalist led to the same effector binding site being able to accommodate both fixed interconnections between the rings (glaucine and rotundine) and flexible interconnections (noscapine, papaverine, and THP). Common binding motifs for the isoquinoline ring abutted divergent benzyl binding pockets, all the while still allowing for extremely fine discriminations (papaverine and THP differ by only two hydrogen atoms).

One of the newly evolved biosensors (THP4.2) was introduced into a genetic circuit for the further evolution of a biosynthetic pathway for the therapeutic alkaloid tetrahydropapaverine. By identifying and broadening the substrate specificity of GfOMT1, we streamlined a more complex set of methylations for the production of THP to a single enzyme, with yield increasing from undetectable to 1.5 mg/L, providing an excellent starting point for additional scaling, especially since upstream biosynthetic machinery to enable *de novo* THP biosynthesis is already well established^**31**^. In particular, increasing the supply of the early intermediate L-DOPA and cofactor S-adenosylmethionine should greatly improve yield^**32,33**^.

These efforts should have immediate practical benefits, as THP is a precursor to several modern pharmaceuticals, including atracurium and cisatracurium used as first line drugs in the management of COVID-19 patients, which have been at risk of experienced shortages due to increased demand^**34**^. Another drug synthesized using THP, papaverine, is used for surgery on blood vessels. A shortage of this drug during 2016 in the US forced surgeons to trial other medications unapproved for these surgeries^**35**^. The combined sequence and structural data can also now inform intelligent library design for identifying additional biosensors for ligands bearing the isoquinoline moiety, or even related groups, such as the quinoline and indole moieties abundant in natural and synthetic pharmaceuticals^**36**^, potentially providing engineered sensors that could accelerate ongoing efforts to improve morphinan producing strains^**3**^.

While previous efforts have largely focused on individually improving either sensitivity, specificity, or background individually^**10–13**^, we show that by combining screening and selection all of these properties can be improved concurrently. Desired biosensor performance outcomes can be finely and rationally tuned during the implementation of SELIS; for example, the stringency for repression can be altered by modifying the strength of the promoter expressing the sensor, wherein a weaker promoter would select for variants that repress more strongly. Moreover, biosensors with varying ligand sensitivities that arise during intermediate rounds of selection, could be successively employed through multiple stages of strain development as product titers improve

As biocatalyst engineering projects become increasingly ambitious, via reconstituting long pathways in microbial hosts^**37**^ or evolving enzyme cascades for pharmaceutical synthesis^**38**^, there will likely be an increased reliance on high-throughput screening capabilities. We envisage that our approach to biosensor development will provide a chemical measurement platform integral to these future endeavors. Beyond their utility in high-throughput screening, biosensors have been used in dynamic regulatory schemes to improve production strain fitness and extend productivity lifetime^**39,40**^. Engineered sensors can also be paired with recently described genetic circuitry to reduce the limit of detection or improve the signal/noise ratio^**41,42**^. Furthermore, since TetR can be employed as a repressor in a wide range of medically and industrially relevant hosts, such as yeasts^**43**^, plants^**44**^, and mammalian cells^**45**^, via a simple ‘road-blocking’ mechanism, the prospect now exists for evolving useful regulatory elements in bacteria and then directly adapting them to eukaryotic systems.

## METHODS

### Strains, plasmids, and media

*E. coli* DH10B (New England BioLabs, Ipswich, MA, USA) was used for all routine cloning and directed evolution. All biosensor systems were characterized in *E. coli* DH10B. *E. coli* BL21 DE3 (New England BioLabs, Ipswich, MA, USA) was used for protein expression. LB-Miller (LB) media (BD, Franklin Lakes, NJ, USA) was used for routine cloning, fluorescence assays, directed evolution, and orthogonality assays unless specifically noted. Terrific broth (TB) (Thermo Fisher Scientific, CAT#: 22711022) was used for protein purification. LB + 1.5% agar (BD, Franklin Lakes, NJ, USA) plates were used for routine cloning and directed evolution. The plasmids described in this work were constructed using Gibson assembly and standard molecular biology techniques. Synthetic genes, obtained as gBlocks, and primers were purchased from IDT. Relevant plasmid sequences are provided in **Supplementary Table 4** and those for final alkaloid sensors are available through Addgene. The pSelis plasmid can be requested from the corresponding authors.

### Benzylisoquinoline alkaloids

Cells were induced with the following chemicals: norlaudanosoline (NOR) (HDH Pharma Inc. CAT#: 29030); tetrahydropapaverine (THP) (Tokyo Chemical Company, product#: N0918); papaverine (PAP) (MP Biomedicals LLC. CAT#: 190261); glaucine (GLAU) (Carbosynth Ltd. product#: FG137572); rotundine (ROTU) (Alfa Aesar, product#: J63328); noscapine (NOS) (Aldrich, SKU: 363960-5G); norreticuline (NRT) (Selena Chem Ltd. product#: CSC000735172).

### Chemical transformation

For routine transformations, strains were made competent for chemical transformation. 5 mL of an overnight culture of DH10B cells were subcultured into 500 mL of LB media and grows at 37°C, 250 r.p.m. for 3 h. Cultures were centrifuged (3,500 g, 4 °C, 10 min), and pellets were washed in 70 mL of chemical competence buffer (10% glycerol, 100mM CaCl2) and centrifuged again (3,500 g, 4°C, 10 min). The resulting pellets were resuspended in 20 mL of chemical competence buffer. After 30 minutes on ice, cells were divided into 250 μL aliquots and flash frozen in liquid nitrogen. Competent cells were stored at −80 °C until use.

### Promoter design and biosensor response assay

Promoters for TtgR and QacR were derived from the literature^**16,19**^. For the RamR promoter, a region 60 base pairs upstream the known operator sequence as well as the operator itself was extracted from the *Salmonella typhimurium* genome (WP_000113609.1). NalD and SmeT are homologs of TtgR, therefore modifications from the Pttgr promoter were made to match the sequence of the NalD operator^**20**^ and SmeT operator^**21**^. For the Pbm3r1, the known Bm3R1 operator^**19**^ was placed immediately after the −10 region of a synthetic medium strength promoter. All promoter sequences are listed in **SI Fig. 2**. The pReg and pGFP equivalents for each regulator were co-transformed into DH10B cells and plated on an LB agar plate with appropriate antibiotics. Three separate colonies were picked for each transformation and were grown overnight. The following day, 20 μL of each culture was then used to inoculate six separate wells within a 2 mL 96-deep-well plate (Corning, Product #: P-DW-20-C-S) sealed with an AeraSeal film (Excel Scientific, Victorville, CA, USA) containing 900 μL of LB media, one for each test ligand and a solvent control. After two hours of growth at 37°C cultures were induced with 100uL of LB media containing either 10 μL of DMSO or 100 μL of LB media containing one of the five target BIAs dissolved in 10 μL of DMSO. Cultures were grown for an additional 4 hours at 37°C, 250 r.p.m and subsequently centrifuged (3,500 g, 4°C, 10 min). Supernatant was removed and cell pellets were resuspended in 1 mL of PBS (137 mM NaCl, 2.7 mM KCl, 10 mM Na2HPO4, 1.8 mM KH2PO4. pH 7.4). 100 μL of the cell resuspension for each condition was transferred to a 96 well microtiter plate (Corning, Product #: 3904), from which the fluorescence (Ex: 485 nm, Em: 509 nm) and absorbance (600 nm) was measured using the Tecan Infinite M1000 plate reader.

### RamR library design and construction

Five semi-rational libraries were designed, each targeting three inward-facing residues on one of five helices of the RamR ligand binding pocket (**Fig. 1d** and **SI Fig. 4**). Libraries were generated using overlap PCR with redundant NNS codons using Accuprime *Pfx* (Thermo Fisher, CAT#: 12344024) and cloned into pReg. *E. coli* DH10B bearing pSelis was transformed with the resulting library. Transformation efficiency always exceeded 10^6^ for each round of selection, indicating several fold coverage of the library. Transformed cells were grown in LB media overnight at 37°C in carbenicillin and chloramphenicol.

### Directed evolution of RamR biosensors

Twenty μL of cell culture bearing the sensor library was seeded into 5 mL of fresh LB containing appropriate antibiotics, 100 μg/mL zeocin (Thermo Fisher. CAT#: R25001), and 100 μM of non-target BIAs (for rounds three and four) and were grown at 37°C for seven hours. Following incubation, 0.5 μL of culture was diluted into 1 mL of LB media, from which 100 μL was further diluted into 900 μL of LB media. 300 μL of this mixture was then plated across three LB agar plates containing carbenicillin, chloramphenicol and the target BIA dissolved in DMSO. Plates were incubated overnight at 37°C. The following day the brightest colonies were picked and grown overnight in 1 mL of LB media containing appropriate antibiotics within a 96-deep-well plate sealed with an AeraSeal film at 37°C. A glycerol stock of cells containing pSelis and pReg bearing the parental RamR variant was also inoculated in 5mL of LB for overnight growth.

The following day, 20 μL of each culture was used to inoculate two separate wells within a new 96-deep-well plate containing 900 μL of LB media. Additionally, eight separate wells containing 1 mL of LB media were inoculated with 20 μL of the overnight culture expressing the parental RamR variant. A typical arrangement would have 44 unique clones on the top half of the plate, duplicates of those clones on the bottom half of the plate, and the right-most column occupied by cells harboring the parental RamR variant. After 2 hours of growth at 37°C the top half of the 96-well plate was induced with 100 μL of LB media containing 10uL of DMSO whereas the bottom half of the plate was induced with 100 μL of LB media containing the target BIA dissolved in 10 μL of DMSO. The concentration of BIA used for induction is typically the same concentration used in the LB agar plate for screening during that particular round of evolution. Cultures were grown for an additional 4 hours at 37°C, 250 r.p.m and subsequently centrifuged (3,500 g, 4°C, 10 min). Supernatant was removed and cell pellets were resuspended in 1mL of PBS. 100 μL of the cell resuspension for each condition was transferred to a 96 well microtiter plate, from which the fluorescence (Ex: 485 nm, Em: 509 nm) and absorbance (600 nm) was measured using the Tecan Infinite M1000. Clones with the highest signal-to-noise ratio were then sequenced and subcloned into a fresh pReg vector.

For sensor variant validation, the subcloned pReg vectors expressing the sensor variants were transformed into DH10B cells bearing pGFP. These cultures were then assayed, as described “Response function measurements” using eight different concentrations of the target BIA. Sensor variants that displayed a combination of a low background, a reduced EC_50_ for the target BIA, and a high signal/noise ratio were used as templates for the next round of evolution.

### Dose response measurements

Glycerol stocks (20% glycerol) of strains containing the plasmids of interest were inoculated into 1 mL of LB media and grown overnight at 37 °C. 20 μL of overnight culture was seeded into 900 μL of LB media containing ampicillin and chloramphenicol within a 2 mL 96-deep-well plate sealed with an AeraSeal film. Following growth at 37°C, 250 r.p.m. for two hours, cultures were induced with 100 μL of a LB media solution containing appropriate antibiotics and the inducer molecule dissolved in 10 μL of DMSO. Cultures were grown for an additional four hours at 37 °C, 250 r.p.m and subsequently centrifuged (3,500 g, 4 °C, 10 min). Supernatant was removed and cell pellets were resuspended in 1 mL of PBS. 100 μL of the cell resuspension for each condition was transferred to a 96-well microtiter plate, from which the fluorescence (Ex: 485 nm, Em: 509 nm) and absorbance (600 nm) was measured using the Tecan Infinite M1000 plate reader.

### Orthogonality assays

For each evolutionary lineage (for example, WT, THP1, THP2, THP3, THP4) all regulators were expressed on the pReg plasmid using the same promoter, which is P114-RBS(riboJ), P114-RBS(riboJ), P103-RBS(elvJ), P114-RBS(riboJ), and P103-RBS(riboJ) for the GLAU, NOS, PAP, ROTU, and THP lineages, respectively. These plasmids were co-transformed with pGFP and the following day three individual colonies were picked into LB and grown overnight. Fluorescence assays were performed as in the “Dose response measurements” section above, but either 100 μM of each BIA in 1% DMSO or 1% DMSO itself was used for induction.

### Protein purification

Coding sequences for RamR variants were cloned into an ampicillin resistant pUC plasmid with a T7 RNA polymerase promoter driving the gene of interest with an N-terminal His6-3C tag. Plasmids were transformed into electrocompetent BL21 DE3 cells and single transformants were grown to saturation in LB supplemented with 1,000 μg/mL carbenicillin. Cultures were diluted 1/250 in terrific broth supplemented with antibiotics in baffled flasks and incubated at 37 °C with agitation (250 r.p.m.) until reaching mid-log phase. Protein expression was induced by addition of IPTG to achieve a final concentration of 0.5 mM. For PAP4 only, papaverine was also added during IPTG induction to reach a final concentration of 100 μM. Cells were cultured for 18 hours at 18 °C. Cells were harvested by centrifugation at 8,000 g for 10 min and the cell pellets were resuspended in 25 mL of wash buffer (50 mM K2HPO4, 300 mM NaCl, and 10% glycerol at pH 8.0) with protease inhibitor cocktail (cOmplete, mini EDTA free, Roche) and lysozyme (0.5 mg/mL). Cells were incubated for 20 min at 4 °C with gentle agitation and lysed by sonication (Model 500, Fisher Scientific). Lysate was repeatedly clarified by centrifugation (35,000xg for 30 min), and protein was recovered by immobilized metal ion affinity chromatography (IMAC) using Ni-NTA resin and gravity flow columns. Eluate was concentrated and dialyzed, with 3C protease added to the dialysis cassette, into the appropriate buffer followed by purification to apparent homogeneity by size exclusion fast protein liquid chromatography (FPLC). All RamR variants were dialyzed into 20 mM Tris (pH 8.0), 200 mM NaCl and 3 mM DTT.

### X-ray crystallography

To form co-crystals of RamR variants in complex with individual ligands, 1mM substrate was added to 10 mg/ml of purified protein and incubated overnight at 4°C except for PAP4 protein, which already formed complex with papaverine during the protein expression step. Rod-shaped co-crystals grew by using sitting-drop vapor diffusion method at room temperature for PAP4, ROTU, GLAU4, and NOS4 in conditions containing 0.1M MES (pH 6.0 – 7.5), 14 – 23% PEG 3350, 0.2 M Ammonium Sulfate, and 0.1 M Sodium Chloride. Individual crystals were flash-frozen directly in liquid nitrogen after brief incubation with a reservoir solution supplemented with 25% (v/v) glycerol. X-ray diffraction data were collected at BL 5.0.1 beamline in ALS (Berkeley, CA). X-ray diffraction was processed to 1.6Å, 1.8Å, 2.0Å, and 2.2Å resolution for PAP4 with papaverine, ROTU4 with rotundine, GLAU4 with glaucine, and NOS4 with noscapine using HKL2000. In Phenix software, phases were obtained by molecular replacement using a previously solved RamR wildtype structure as the initial search model (PDB code 3VVX). The molecular replacement solutions for each structure were iteratively built using Coot and Phenix refine package. The quality of the final refined structures was evaluated by MolProbity. The final statistics for data collection and structure determination are shown in **SI Table 2.**

### Biosensor-linked O-Methyltransferase activity assay

All OMTs were expressed with the P114-RBS(riboJ) promoter/RBS on the pReg plasmid backbone (no regulator present). Cells were co-transformed with both the OMT plasmid and the THP reporter plasmid and plated on an LB agar plate containing appropriate antibiotics. Three individual colonies from each transformation were picked into LB and grown overnight. Resulting cultures were diluted 50-fold into 1mL of LB media containing norlaudanosoline and 1 mg/mL of ascorbic acid in a 96-deep-well plate and were grown at 30 °C for 18 hours. Subsequently, the fluorescence of cultures was measured in the same manner as previously described in “Dose response measurement” above.

### O-Methyltransferase evolution

The OMT coding region of the GfOMT1 expression vector was targeted for random mutagenesis introducing an average of two mutations relative to the template. Libraries were assembled using Gibson assembly and were subsequently transformed into chemically competent cells containing the THP reporter plasmid. In parallel, the template OMT was transformed into the same cells for downstream comparative analysis. Cultures were then plated on an LB agar plate containing norlaudanosoline. Highly fluorescent colonies were picked and grown overnight in appropriate antibiotics. The following day, cultures were grown with norlaudanosoline and 1 mg/mL of ascorbic acid at 30 °C for 18 hours. Fluorescence of the resulting cultures were measured, as described above and variants that were significantly more fluorescent than the template OMT were sequenced. Unique top-performing OMT variants were then subcloned into the pReg vector and transformed into cells bearing the THP reporter plasmid. Three individual colonies from each transformation were subcultured into fresh LB media and the fluorescence of each culture in the presence of norlaudanosoline (refer to **SI Fig. 15**) was subsequently measured, as described above. The top performing OMT variant was then used as the template for the following round of evolution.

### Homology model

The homology model of the GEN5 GfOMT1 variant was constructed using SWISS-MODEL (https://swissmodel.expasy.org/). 5ICE was used as a template for structure generation.

### Liquid-chromatography/Mass spectrometry

Cells containing the plasmid expressing each OMT variant with the P114-RBS(RiboJ) promoter were transformed and plated onto an LB agar plate containing appropriate antibiotics. The following day, three colonies from each plate were cultured overnight in LB and subsequently diluted 50-fold into 1mL of LB containing 10 μM of NOR and 1mg/mL ascorbic acid. These cultures were grown for 18 hours at 30 °C, centrifuged at 16,000xg for 1 minute, and the resulting supernatant was filtered using a 0.2 μM filter. The samples were then diluted 1:100 fold into filter sterilized water prior to LC/MS analysis. Samples were analyzed using an Agilent 6546 Q-TOF LC/MS with a dual Agilent Jet Stream electrospray ionization (ESI) source in positive mode. Chromatographic separations were obtained under gradient conditions by injecting 5 μL onto an Agilent RRHD Eclipse Plus C18 column (50 × 2.1 mm, 1.8 micron particle size) with an Agilent Zorbax Eclipse Plus C18 narrow bore guard column (12.5 × 2.1 mm, 5 micron particle size) on an Agilent 1260 Infinity II liquid chromatography system. The mobile phase consisted of eluent A (water + 0.1% formic acid) and eluent B (acetonitrile). The gradient was as follows: 5% B to 95% B from 0 to 6 min (0.3 mL/min), held at 95% B from 6 to 7 min (0.3 mL/min), 95% B to 5% B from 7 to 7.1 min (0.3 mL/min), and held at 5% B from 7.1 to 9 min (0.45 mL/min). The sample tray and column compartment were set to 7 °C and 30 °C, respectively. The fragmentor was set to 180 V. Q-TOF data was processed using the Agilent MassHunter Qualitative Analysis software.

To create chromatograms shown in **SI Fig. 18**, signal count from the EIC within a window +/- 0.05 minutes relative to the retention time for the corresponding alkaloid, see **SI Table 3**, was extracted for each scan (m/z ratios of the following: 288.1230, 302.1387, 316.1543, 330.1700, and 344.1856) and added together. Since a significant background peak was detected around the expected retention time for THP, the sigal count in this window of time was removed for all samples that had a count at or below this background level. The resulting chromatograms were then used to create **Fig. 5d.**

### Statistical analysis and reproducibility

All data in the manuscript are displayed as mean ± s.e.m. unless specifically indicated. Bar graphs, fluorescence/growth curves, dose response functions, and orthogonality matrices were all plotted in Python 3.6.9 using matplotlib and seaborn. Dose response curves and EC_50_ values were estimated by fitting to the hill equation y = d + (a-d)*x^b^ / (c^b^ + x^b^) (where y = output signal, b = hill coefficient, x = ligand concentration, d = background signal, a = the maximum signal, and c = the EC_50_), with the scipy.optimize.curve_fit library in Python.

## Acknowledgements

Funding from DARPA Soils (HR00111920019), Welch (F-1654), and AFSOR - (FA9550-14-1-0089) is acknowledged. This work is partially supported by grants from the National Institutes of Health (R01GM104896 and R01GM125882 to Y.Z.)

## Accession numbers

Coordinates for the complex structures have been deposited in the Protein Data Bank with: PAP4 in complex with papaverine as 7N53, ROTU4 with rotundine as 7N4W, NOS4 with noscapine as 7N4Z, and GLAU4 with glaucine as 7N54.

## Author contributions

S.D. designed the experiments and performed biosensor evolution and characterization. S.D. and R.T. performed protein purification. Enzyme evolution was done by S.D. and K.J. and X-ray crystallography was conducted by W.K. and N.T.B.. We would like to thank Kristin J Blake, in the Chemistry Department at the University of Texas at Austin, for performing LC/MS analysis. The manuscript was written by S.D. with support from A.D.E, R.T., Y.Z., and H.A. S.D., A.D.E., and H.A. supervised all aspects of the study.

## Corresponding authors

Correspondence to: Simon d’Oelsnitz, and Andrew D Ellington

## Supplementary information

Supplementary figures and tables.

