## Supplemental Information for "Using structurally fungible biosensors to evolve improved alkaloid biosyntheses"

**Supplementary Information:**
Supporting Figures 1-18
Supporting Tables 1-4

### Supplementary Figures

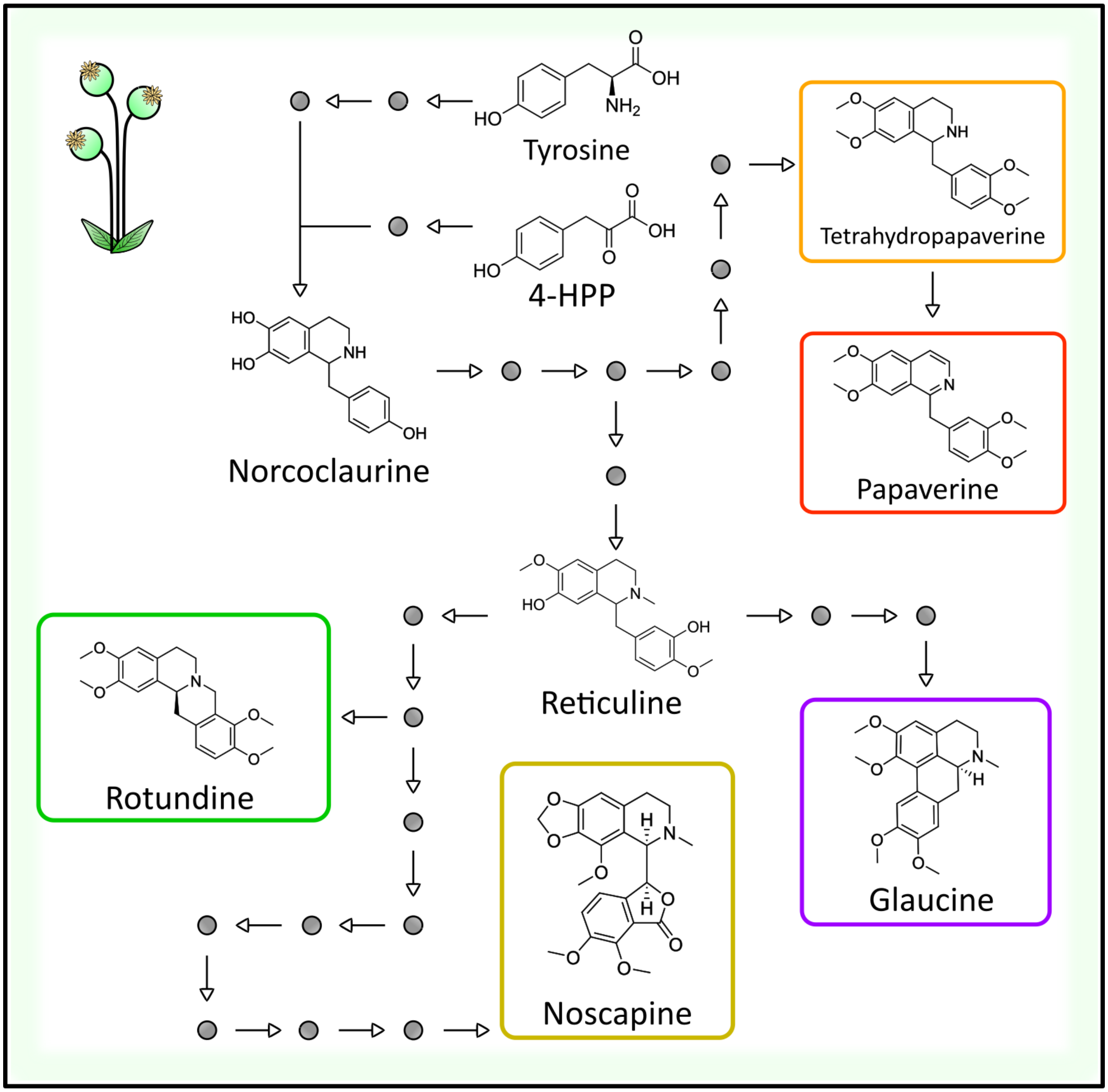


**Supplementary Figure 1**. Benzylisoquinoline pathway map.
Arrows represent enzymatic steps and grey circles represent metabolites. Target alkaloids in this work are highlighted with a colored border.

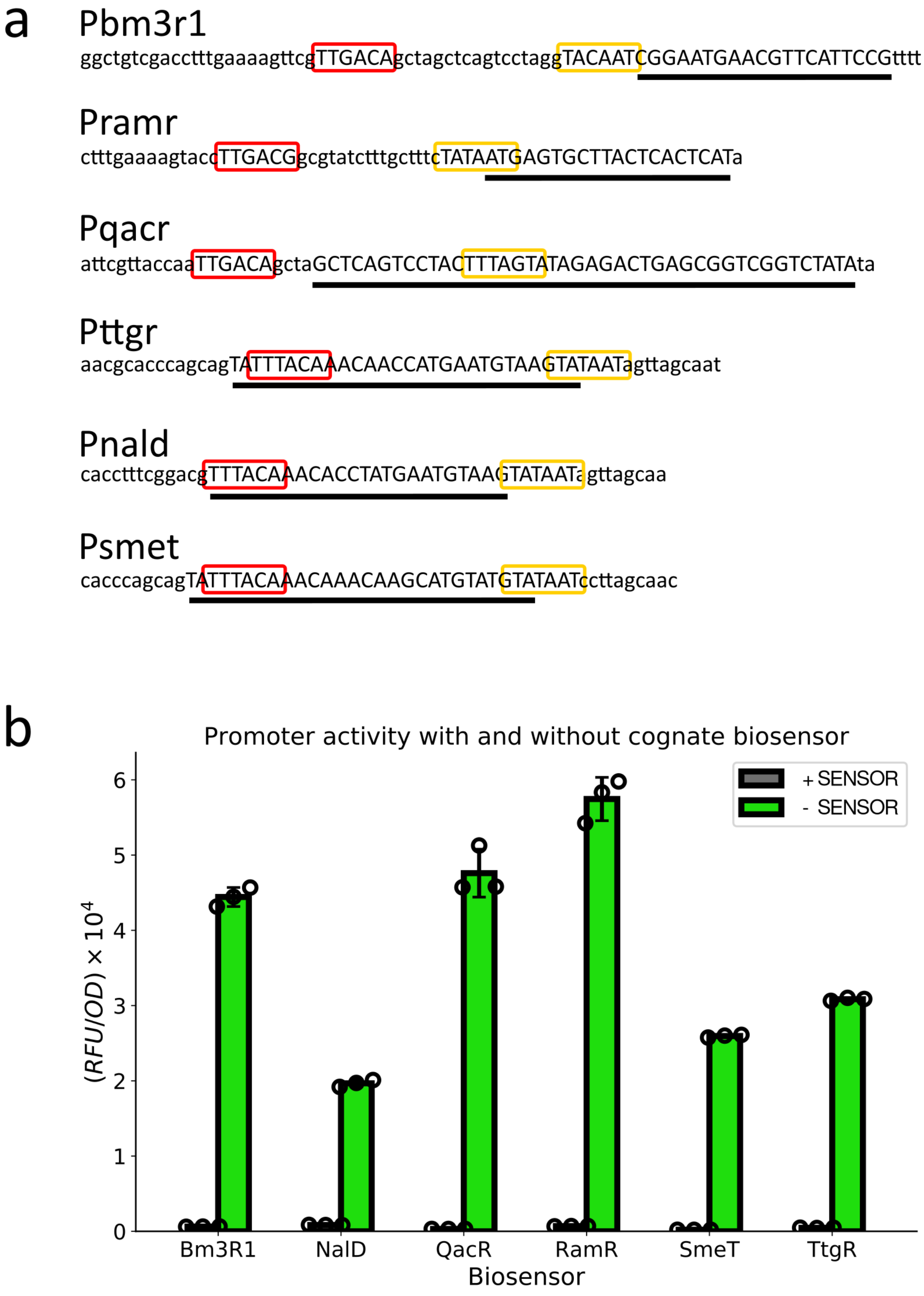


**Supplementary Figure 2**. Multidrug resistance regulator design and validation.
(**a**) Promoter design for each regulator. -35 and -10 promoter regions are highlighted with a red and yellow box, respectively. Operator sequences are underlined. All promoters are followed by the RiboJ riboregulator, a medium strength RBS, the sfGFP gene, and a strong terminator. (**b**) Validation of promoter activity and regulator repression in *E. coli*. Cells were co-transformed with the regulator's promoter and either an empty vector (- Sensor) or a vector expressing the cognate regulator (+ Sensor) and promoter activity was monitored via fluorescence.

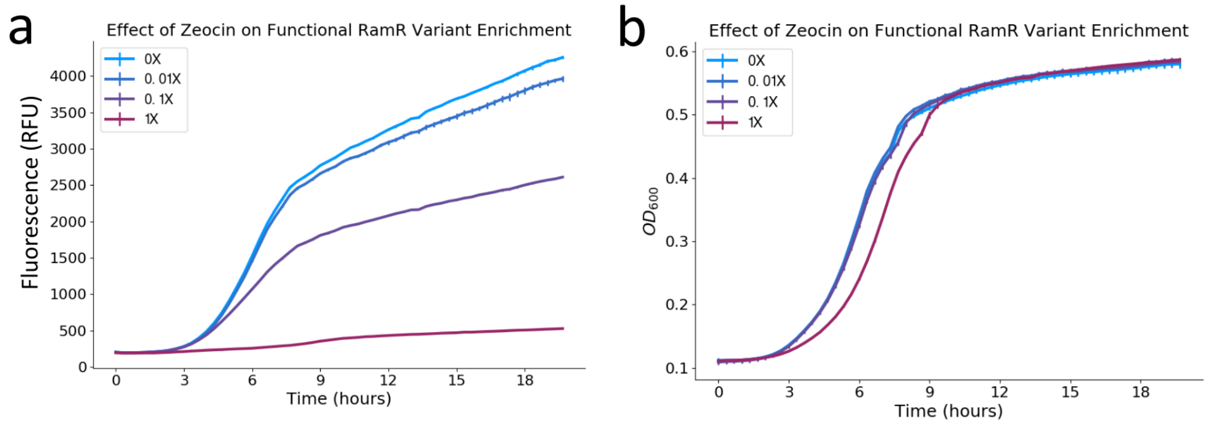


**Supplementary Figure 3**. Trial negative selections with the pZeoG plasmid.
Cells co-transformed with both pReg expressing a library of RamR variants and the pZeoG plasmid were grown for 20 hours in the presence of variable amounts of zeocin and fluorescence (**a**) and cell density (**b**) were monitored. The "1X" concentration represents 100 ug/mL of zeocin. Assays were performed in biological triplicate.

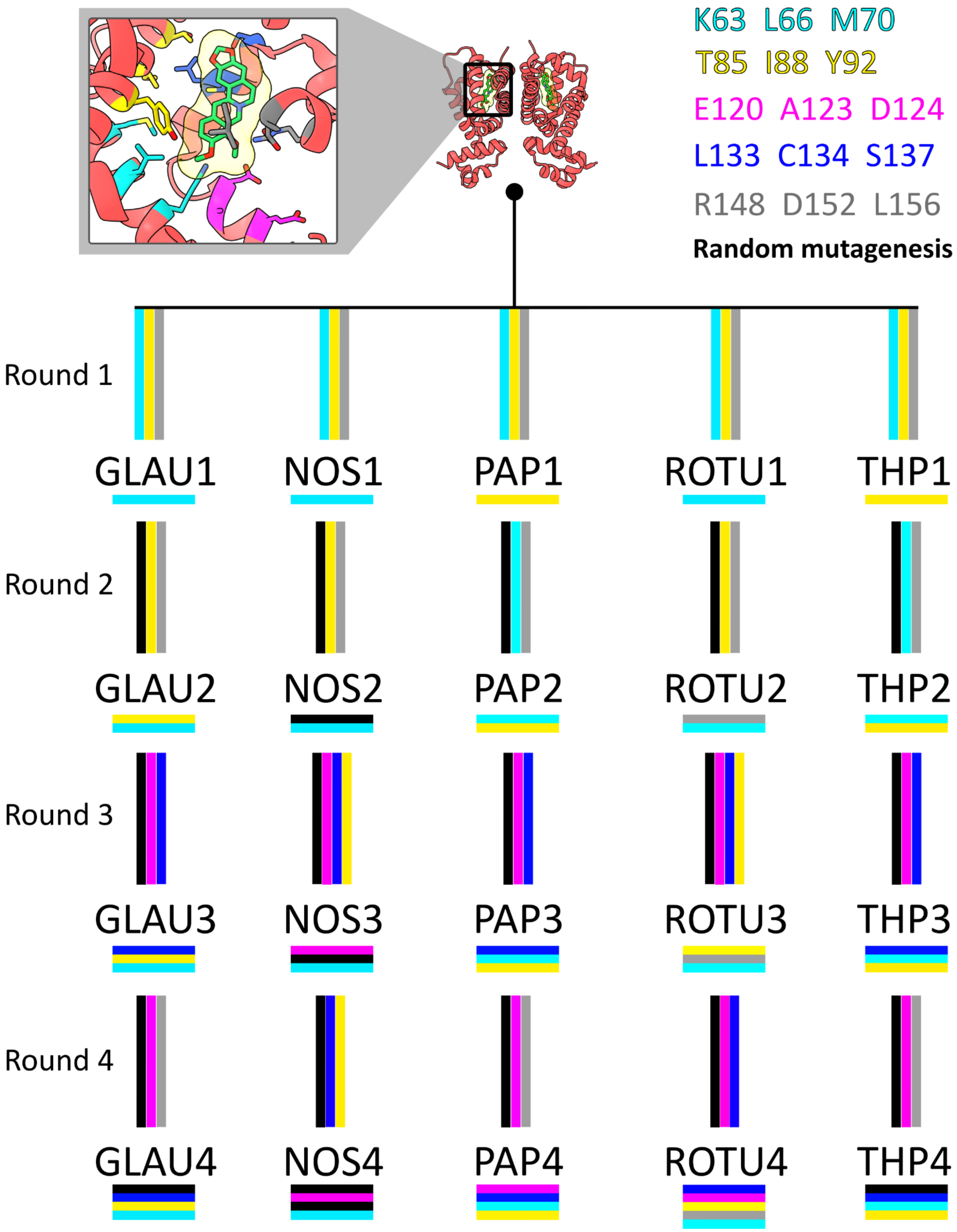


**Supplementary Figure 4**. Visual representation of libraries used throughout evolution.
(top left) A magnified structure of RamR bound to berberine (PDB: 3VW2) displays residues targeted for mutagenesis. These residues were chosen based on their proximity to berberine. (top middle) global structure of RamR. (top right) The mapping of library color code to the corresponding residues targeted for combinatorial site saturation mutagenesis. (bottom) Libraries used and fixed during evolution. Colored vertical lines represent libraries used to introduce diversity prior to selection. Colored horizontal lines represent library positions fixed.

a

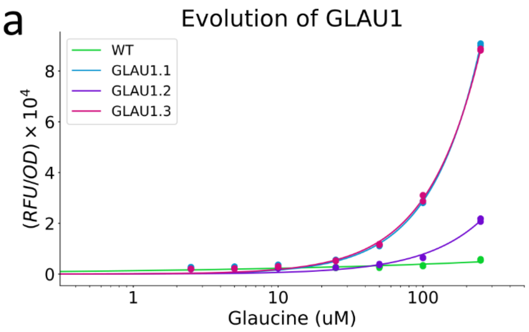


| | WT | GLAU1.1 | GLAU1.2 | GLAU1.3 |
|---|---|---|---|---|
| EC50 | N/A | N/A | N/A | N/A |
| Background | 2203.5 +/- 157.0 | 2490.7 +/- 364.4 | 1926.5 +/- 36.1 | 1765.2 +/- 5.3 |
| New mutations | N/A | T85L I88G Y92Y x2 | T85C I88M Y92F | **K63T L66W M70I** |

b

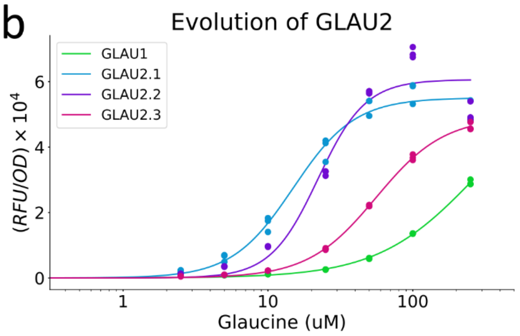


| | GLAU1 | GLAU2.1 | GLAU2.2 | GLAU2.3 |
|---|---|---|---|---|
| EC50 | 272.5 | 14.9 | 22.2 | 55.8 |
| Background | 570.6 +/- 15.2 | 327.0 +/- 36.8 | 305.0 +/- 7.6 | 246.7 +/- 20.1 |
| New mutations | N/A | T85C I88M Y92W | **T85T I88L Y92W** | C134I V138A |

c

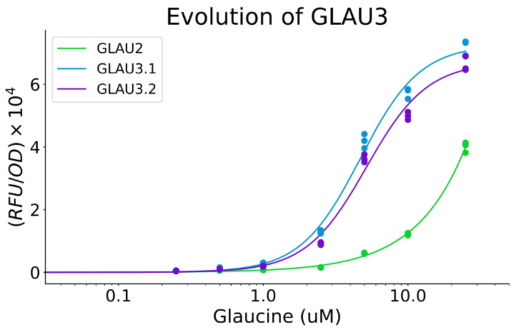


| | GLAU2 | GLAU3.1 | GLAU3.2 |
|---|---|---|---|
| EC50 | N/A | 4.7 | 5.2 |
| Background | 368.6 +/- 15.6 | 293.4 +/- 11.0 | 235.3 +/- 0.4 |
| New mutations | N/A | L133H C134L S137H x2 | **L133H C134L S137N** x4 |

d

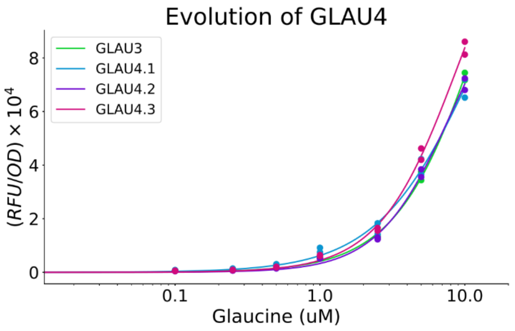


| | GLAU3 | GLAU4.1 | GLAU4.2 | GLAU4.3 |
|---|---|---|---|---|
| EC50 | 13.9 | 12.2 | 8.2 | 9.0 |
| Background | 547.9 +/- 16.0 | 559.6 +/- 10.9 | 346.0 +/- 14.9 | 431.1 +/- 49.4 |
| New mutations | N/A | H99N | **T189A** | T57A |

e

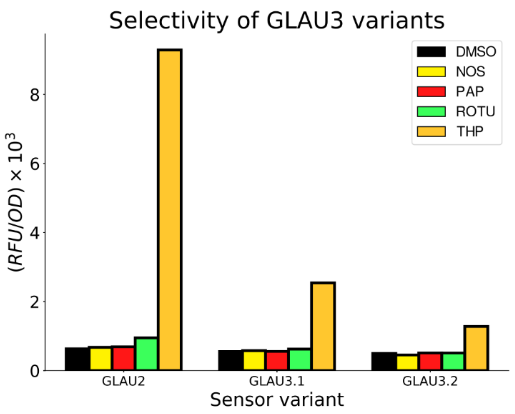


f

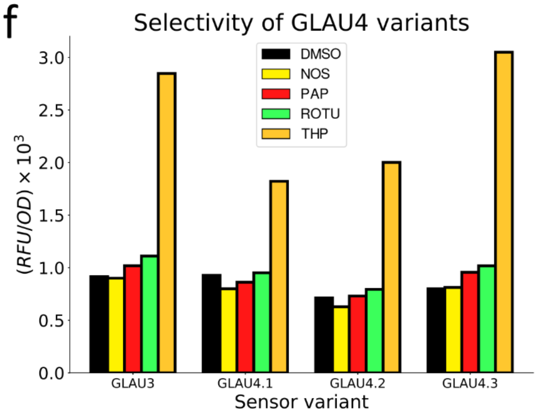


**Supplementary Figure 5.** Performance of all top GLAU variants recovered.
(**a-d**) Dose response functions of top unique GLAU variants. Variants were chosen based on their signal/noise ratio measured during evolution (See **Figure 2c**). All variants were subcloned into a new pReg backbone prior to characterization with the pGFP plasmid. The “x2” symbol denotes that this amino acid sequence was recovered twice following evolution. The variant genotype highlighted in green was chosen as the template for the following round of evolution. Dose response measurements were performed in biological triplicate. (**e,f**) Selectivity of generation three and four sensor variants. Cells were induced with 100 uM of all non-target BIAs, separately.

a

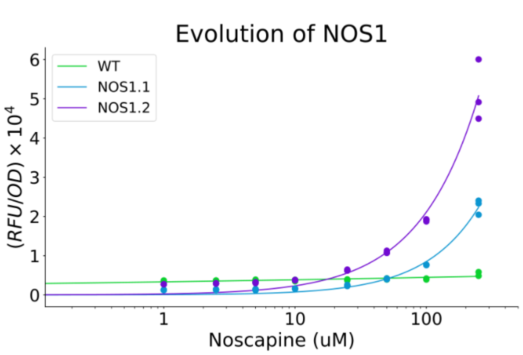


| | WT | NOS1.1 | NOS1.2 |
|---|---|---|---|
| EC50 (uM) | N/A | N/A | N/A |
| Background | 3490.7 +/- 226.3 | 1174.3 +/- 27.4 | 2639.4 +/- 79.1 |
| New mutations | N/A | **K63T L66M M70L** x4 | K63S L66M M70L |

b

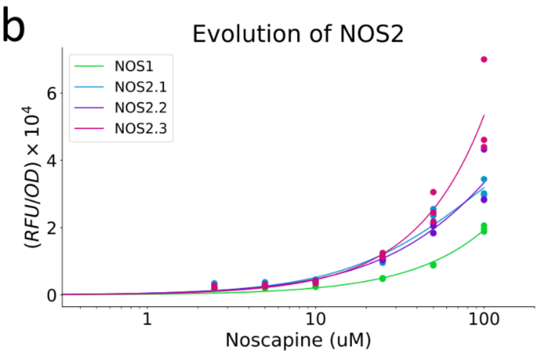


| | NOS1 | NOS2.1 | NOS2.2 | NOS2.3 |
|---|---|---|---|---|
| EC50 (uM) | N/A | 100.4 | 287.2 | N/A |
| Background | 1595.9 +/- 133.5 | 3108.0 +/- 47.2 | 1791.5 +/- 105.2 | 2086.9 +/- 441.9 |
| New mutations | N/A | L156M | L130V V138L | **V138L** |

c

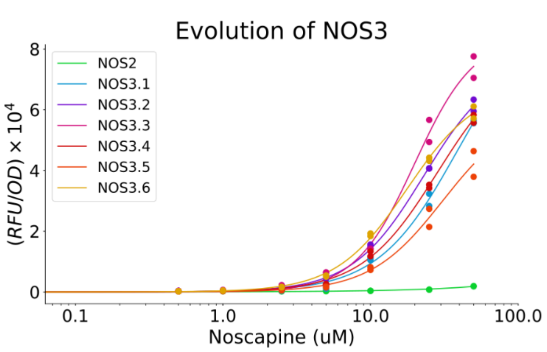


| | NOS2 | NOS3.1 | NOS3.2 | NOS3.3 | NOS3.4 | NOS3.5 | NOS3.6 |
|---|---|---|---|---|---|---|---|
| EC50 (uM) | N/A | 38.8 | 23.0 | 19.9 | 28.4 | 30.8 | 17.7 |
| Background | 271.5 +/- 23.5 | 214.7 +/- 17.9 | 239.3 +/- 27.4 | 252.8 +/- 1.3 | 195.3 +/- 29.5 | 189.3 +/- 6.5 | 169.9 +/- 15.5 |
| New mutations | N/A | E120T A123A D124R x2 | E120S A123A D124R | E120A A123A D124R x2 | E120M A123A D124R | E120R A123G D124R | **E120R A123A D124R** x3 |

d

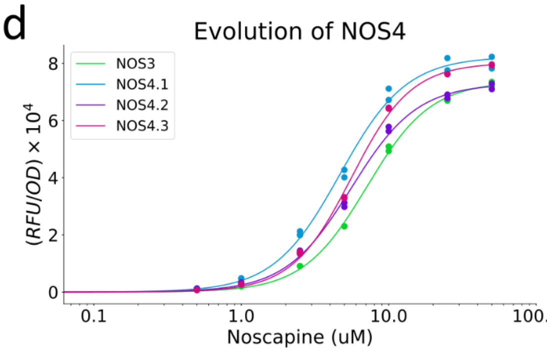


| | NOS3 | NOS4.1 | NOS4.2 | NOS4.3 |
|---|---|---|---|---|
| EC50 (uM) | 7.1 | 4.7 | 5.6 | 5.6 |
| Background | 194.1 +/- 14.2 | 123.0 +/- 14.6 | 173.8 +/- 26.5 | 149.6 +/- 2.4 |
| New mutations | N/A | **Q64H H135Y L156M** | D65N | L97F |

e

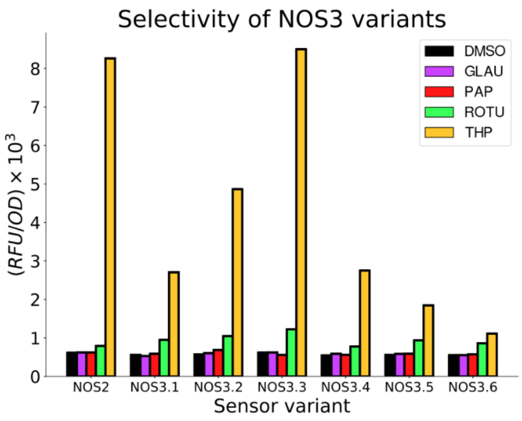


f

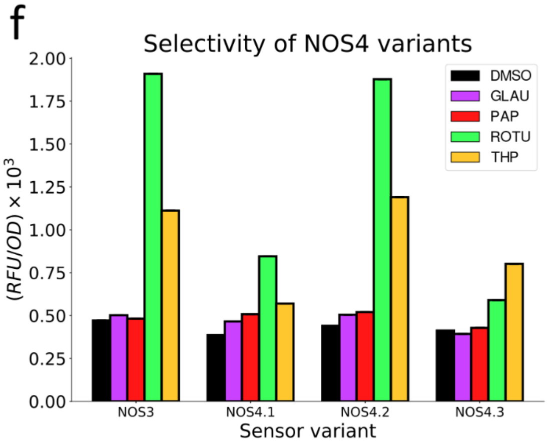


**Supplementary Figure 6.** Performance of all top NOS variants recovered. See Supplementary Figure 5 legend for details.

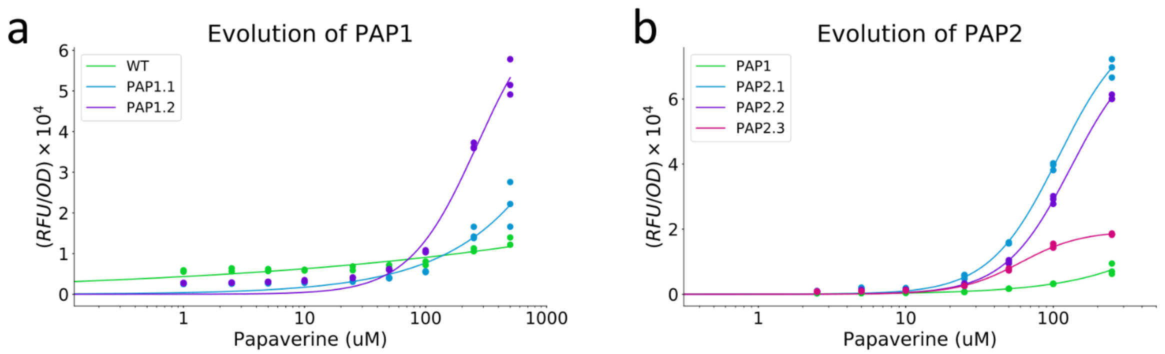


| | WT | PAP1.1 | PAP1.2 |
|---|---|---|---|
| EC50 (uM) | N/A | N/A | 265.6 |
| Background | 5728.3 +/- 309.4 | 2581.0 +/- 201.7 | 2626.8 +/- 31.9 |
| New mutations | N/A | T85L<br>I88F<br>Y92G | **T85M**<br>**I88I** x2<br>**Y92G** |

| | PAP1 | PAP2.1 | PAP2.2 | PAP2.3 |
|---|---|---|---|---|
| EC50 (uM) | N/A | 108.3 | 132.0 | 60.0 |
| Background | 237.1 +/- 13.8 | 697.5 +/- 98.9 | 673.4 +/- 10.4 | 621.5 +/- 16.4 |
| New mutations | N/A | **K63R**<br>**L66L**<br>**M70L** | K63H<br>L66L<br>M70L | C134I x2 |

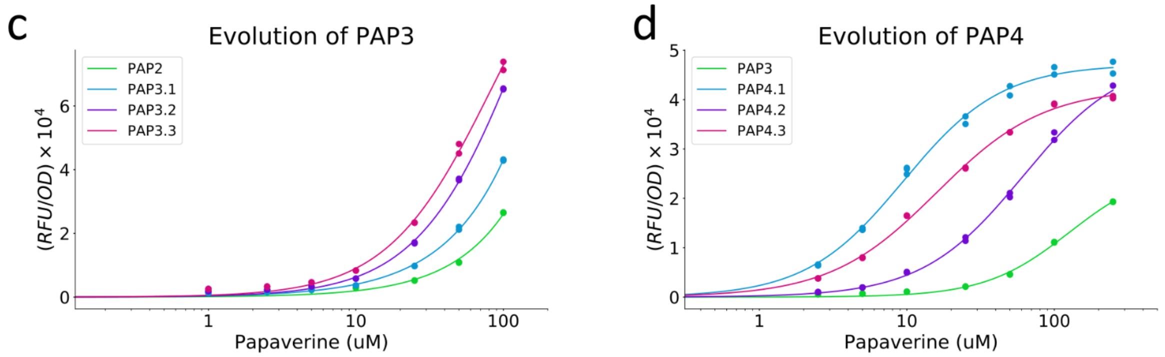


| | PAP2 | PAP3.1 | PAP3.2 | PAP3.3 |
|---|---|---|---|---|
| EC50 (uM) | N/A | N/A | 139.7 | 82.9 |
| Background | 1577.1 +/- 51.0 | 1108.0 +/- 38.2 | 1567.7 +/- 14.5 | 2037.1 +/- 63.4 |
| New mutations | N/A | E120H<br>A123G<br>D124E | L133I<br>C134L<br>S137R | **L133I**<br>**C134L**<br>**S137K** |

| | PAP3 | PAP4.1 | PAP4.2 | PAP4.3 |
|---|---|---|---|---|
| EC50 (uM) | 126.8 | 9.5 | 61.1 | 15.9 |
| Background | 417.1 +/- 19.6 | 1266.0 +/- 49.4 | 508.2 +/- 4.5 | 980.4 +/- 6.5 |
| New mutations | N/A | E120H<br>A123G x3<br>D124L | **E120H**<br>**A123D**<br>**D124L** | E120R<br>A123D<br>D124E |

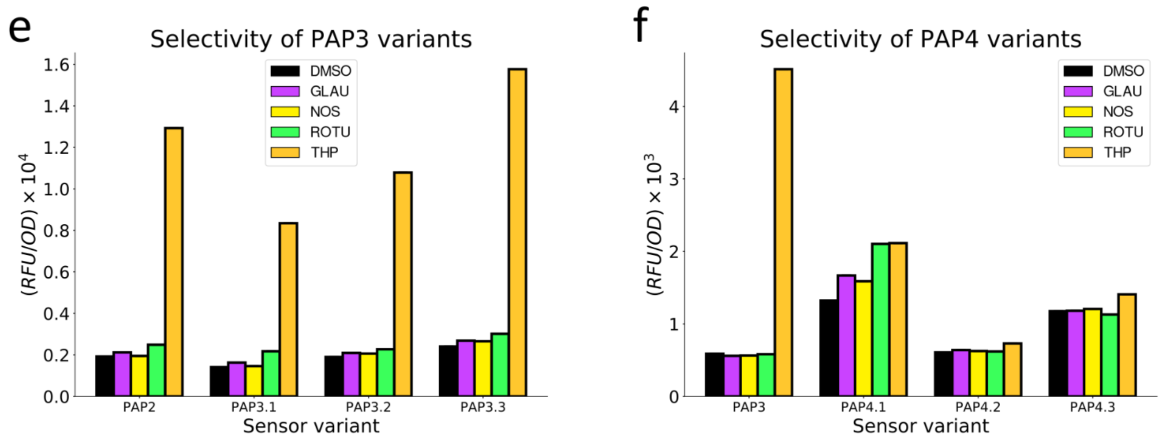


**Supplementary Figure 7**. Performance of all top PAP variants recovered. See Supplementary Figure 5 legend for details.

a

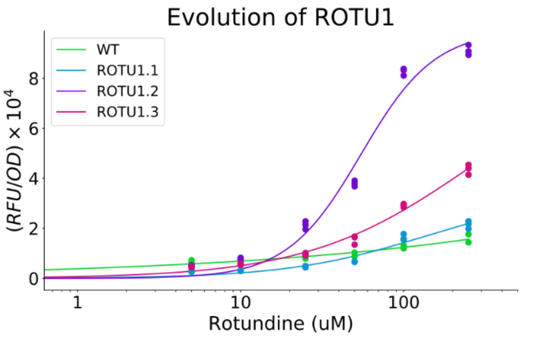


| | WT | ROTU1.1 | ROTU1.2 | ROTU1.3 |
|---|---|---|---|---|
| EC50 (uM) | N/A | 134.5 | 54.8 | 256.3 |
| Background | 5610.5 +/- 149.2 | 2974.2 +/- 90.4 | 4159.6 +/- 115.7 | 4191.9 +/- 195.6 |
| New mutations | N/A | K63K<br>L66A<br>M70L | **K63Y**<br>**L66H**<br>**M70L** | T85S<br>I88M x3<br>Y92I |

b

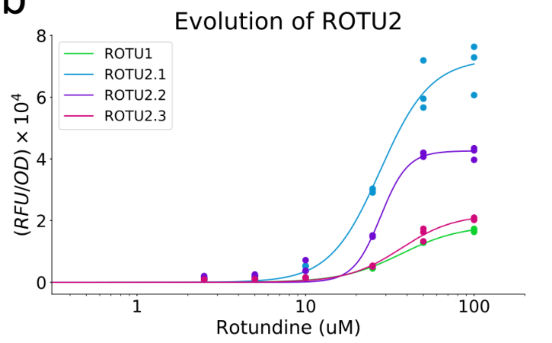


| | ROTU1 | ROTU2.1 | ROTU2.2 | ROTU2.3 |
|---|---|---|---|---|
| EC50 (uM) | 36.9 | 27.7 | 27.9 | 36.8 |
| Background | 720.0 +/- 23.2 | 498.2 +/- 28.1 | 1359.9 +/- 278.4 | 787.0 +/- 53.6 |
| New mutations | N/A | **R148S**<br>**D152K**<br>**L156Y** | C134M | R148K<br>D152N |

c

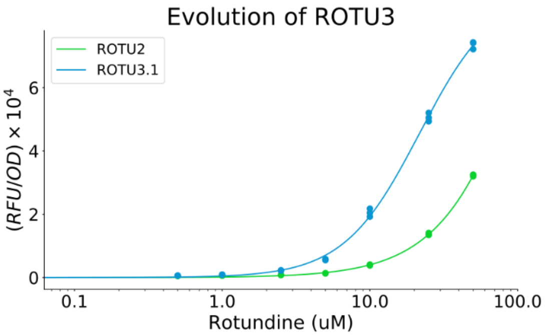


| | ROTU2 | ROTU3.1 |
|---|---|---|
| EC50 (uM) | 172.6 | 21.7 |
| Background | 495.0 +/- 30.8 | 466.9 +/- 40.0 |
| New mutations | N/A | **T85I**<br>**I88T** |

d

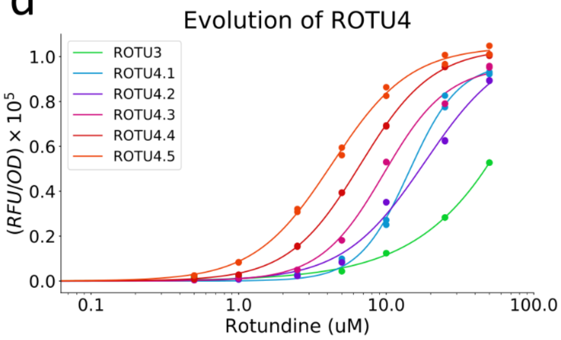


| | ROTU3 | ROTU4.1 | ROTU4.2 | ROTU4.3 | ROTU4.4 | ROTU4.5 |
|---|---|---|---|---|---|---|
| EC50 (uM) | 201.9 | 14.3 | 18.3 | 9.6 | 6.7 | 4.3 |
| Background | 2032.9 +/- 40.6 | 300.1 +/- 28.1 | 261.8 +/- 85.5 | 305.0 +/- 30.6 | 268.9 +/- 15.5 | 303.8 +/- 13.0 |
| New mutations | N/A | E120K<br>A123I<br>D124E | C134L | L133I<br>C134L | E120K<br>A123I<br>D124E C134L | **E120K**<br>**A123I D124E**<br>**L133I C134L** |

e

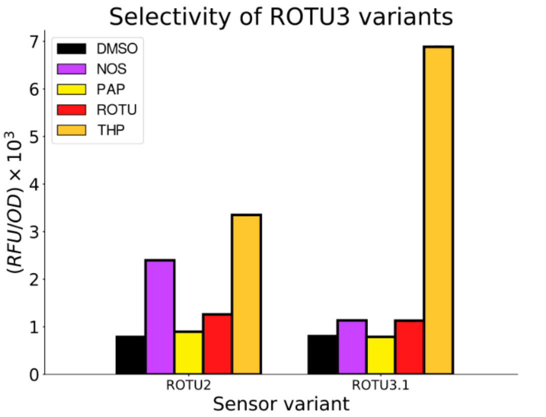


f

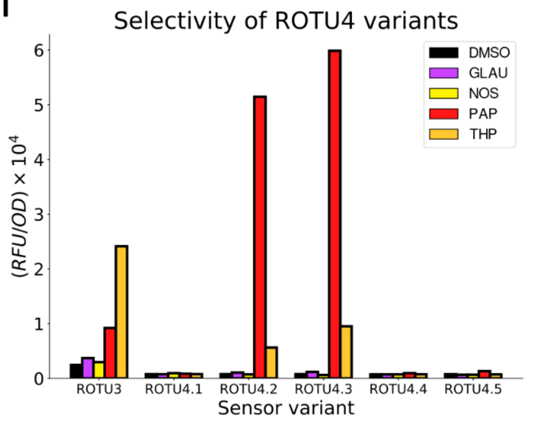


**Supplementary Figure 8.** Performance of all top ROTU variants recovered.
See Supplementary Figure 5 legend for details.

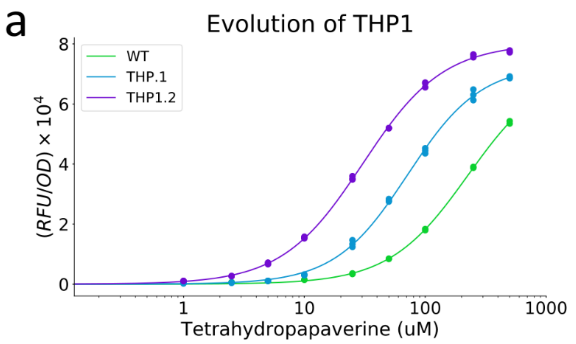


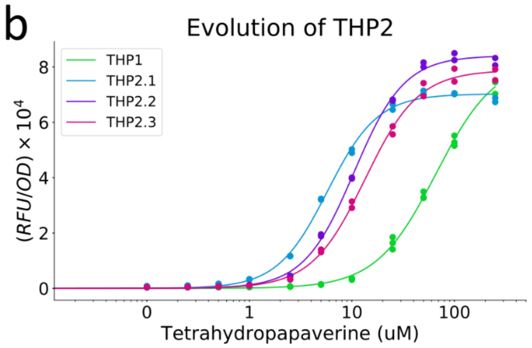


| | WT | THP.1 | THP1.2 |
|---|---|---|---|
| EC50 (uM) | 225.9 | 71.7 | 30.4 |
| Background | 560.1 +/- 44.6 | 200.3 +/- 34.4 | 594.5 +/- 11.4 |
| New mutations | N/A | **T85S I88L Y92I** | T85N I88L Y92I |

| | THP1 | THP2.1 | THP2.2 | THP2.3 |
|---|---|---|---|---|
| EC50 (uM) | 66.1 | 5.8 | 10.5 | 13.4 |
| Background | 353.7 +/- 34.5 | 702.1 +/- 18.7 | 370.2 +/- 12.3 | 385.0 +/- 27.2 |
| New mutations | N/A | K63Y L66W M70A | **K63F L66W M70S** x2 | K63W L66W M70A |

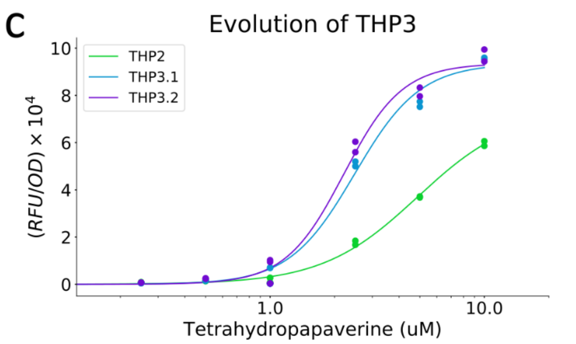


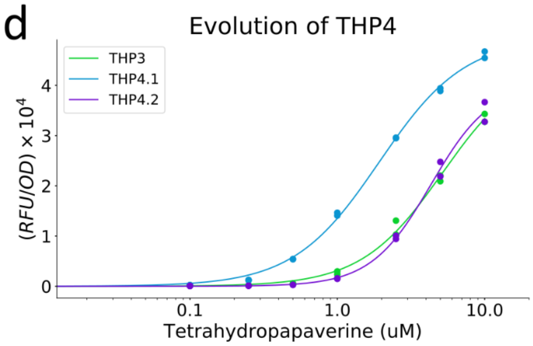


| | THP2 | THP3.1 | THP3.2 |
|---|---|---|---|
| EC50 (uM) | 4.9 | 2.5 | 2.2 |
| Background | 553.9 +/- 34.3 | 283.5 +/- 18.1 | 313.2 +/- 22.6 |
| New mutations | N/A | **L133I C134V S137E** | L133I C134V S137K x2 |

| | THP3 | THP4.1 | THP4.2 |
|---|---|---|---|
| EC50 (uM) | 5.2 | 1.9 | 4.3 |
| Background | 70.8 +/- 10.4 | 96.2 +/- 4.3 | 64.4 +/- 2.2 |
| New mutations | N/A | D169N | **I26T S85T T116S E120V** |

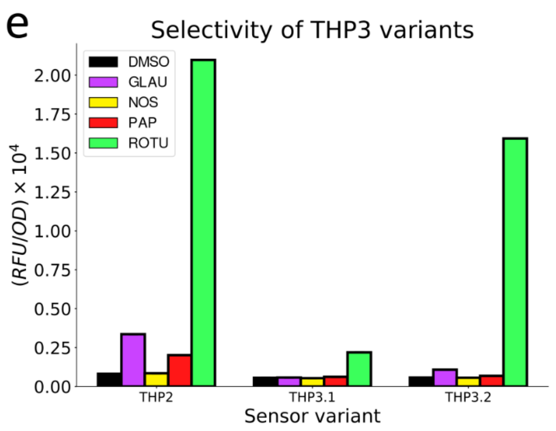


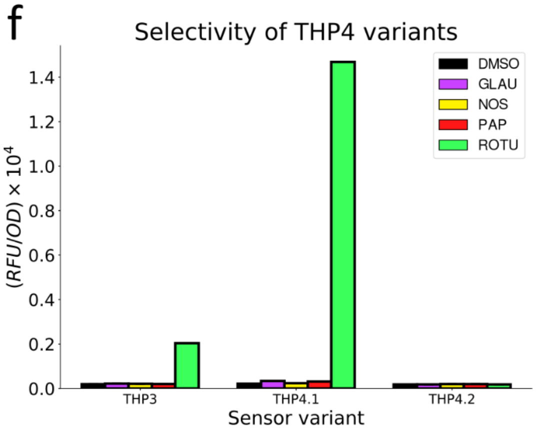


**Supplementary Figure 9.** Performance of all top THP variants recovered.
See Supplementary Figure 5 legend for details.

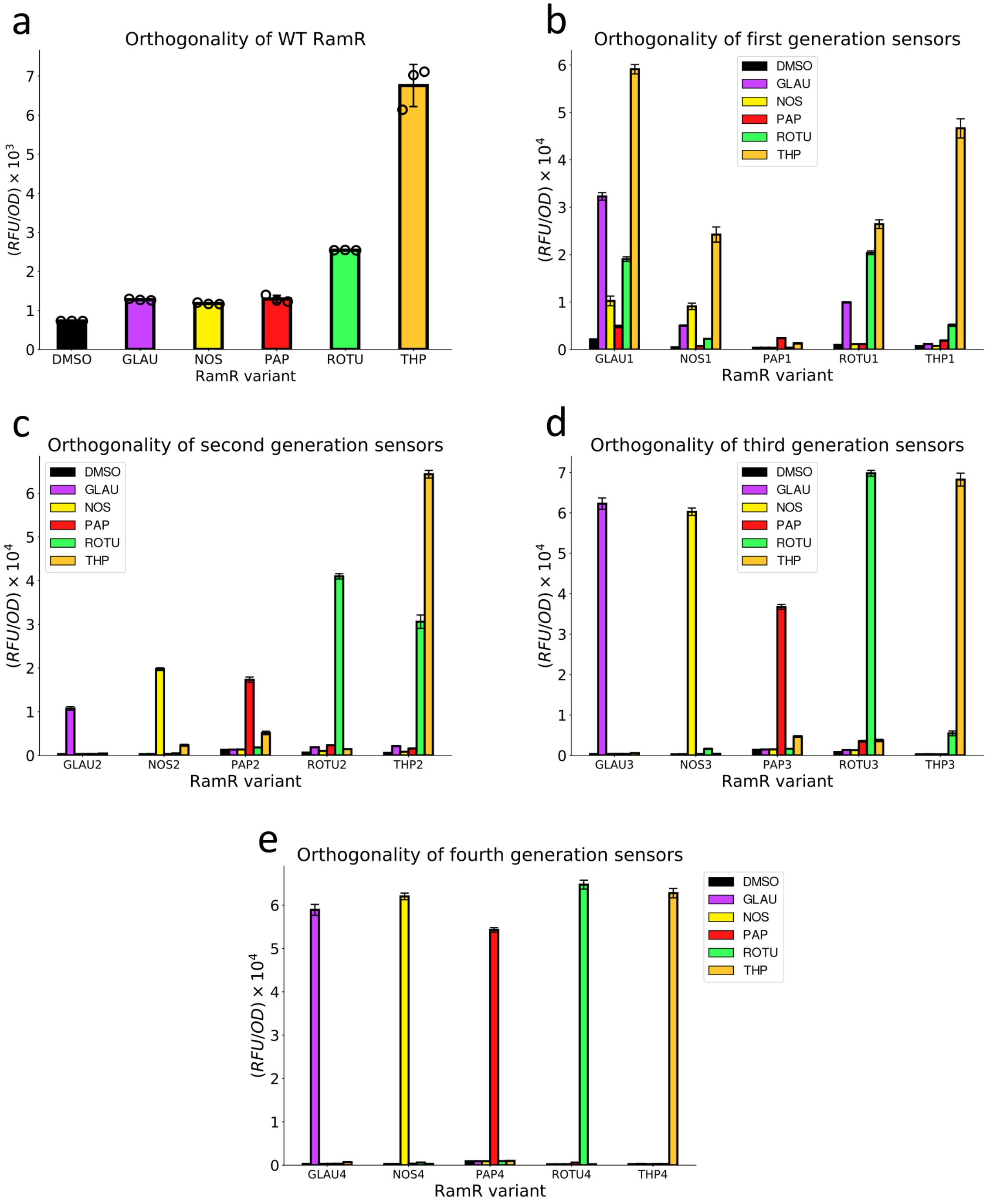


**Supplementary Figure 10.** Orthogonality of all final RamR variants.
Fluorescent response of cells expressing pGFP and pReg with WT RamR (**a**), Gen1 variants (**b**), Gen2 variants (**c**), Gen3 variants (**d**), and Gen4 variants (**e**) that were induced with 100 uM of each BIA, separately. Measurements were performed in biological triplicate. See "Methods - Orthogonality Assays" for the list of promoters used to express each variant.

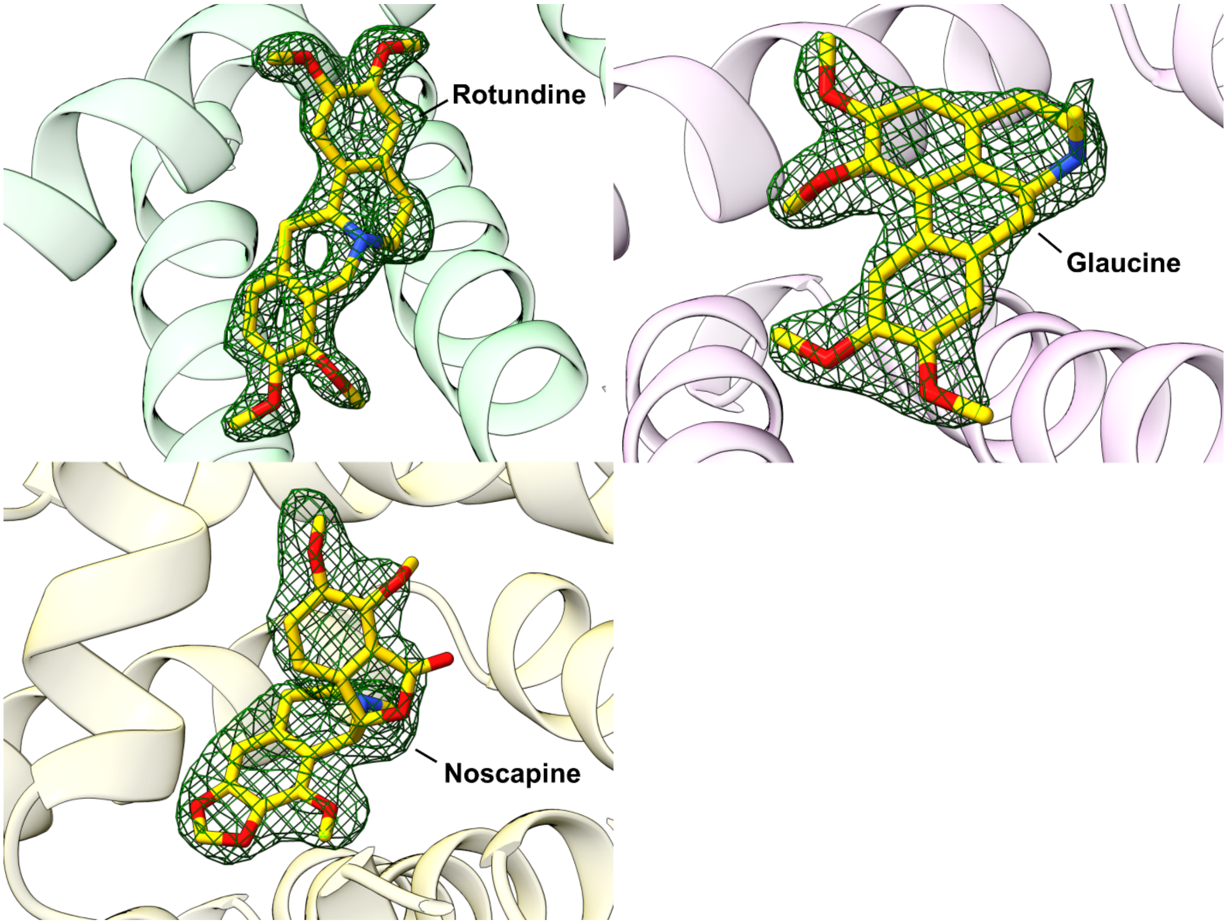


**Supplementary Figure 11.** Electron density omit maps shown for each BIAs in sticks occupying the binding site of each RamR variant (ROTU4 - green, GLAU4 - purple, NOS4 - pale yellow). Fo-Fc maps (contoured at 3.0σ for Rotundine / Glaucine, and 2.5σ for Noscapine) are shown as forest green mesh superimposed on the model of each BIA.

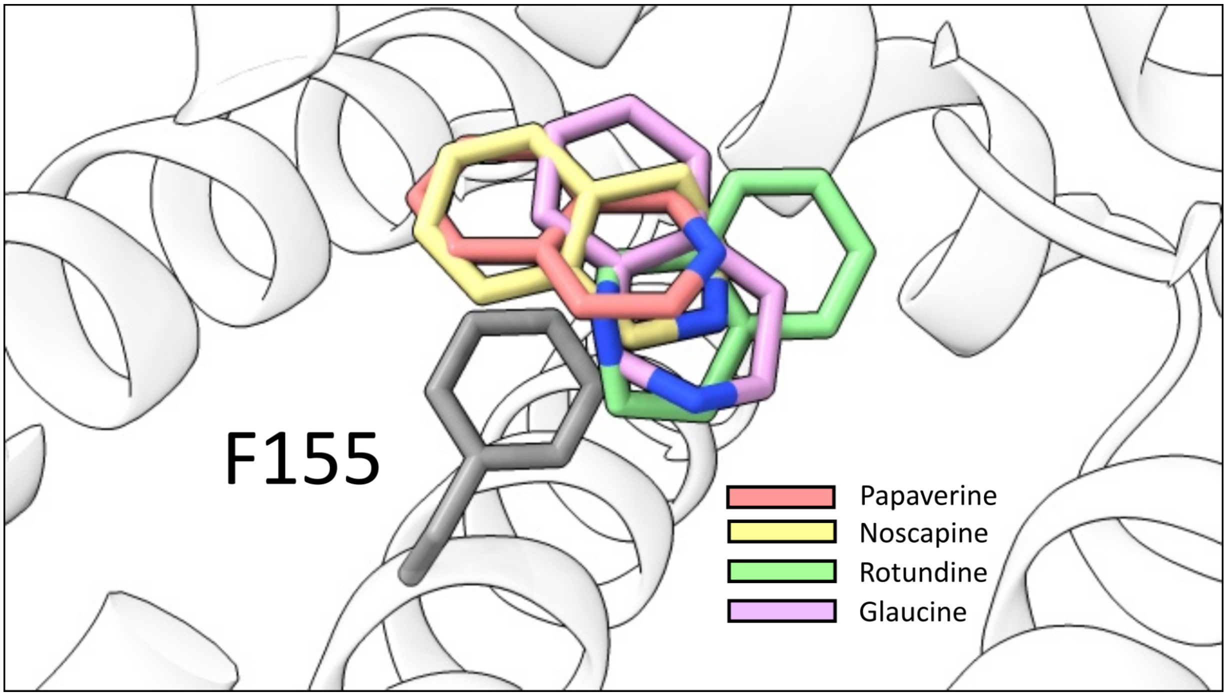


**Supplementary Figure 12**. Superimposed BIA structures from all BIA-specific RamR variants.
The isoquinoline moiety of glaucine, papaverine, noscapine, and rotundine are shown in the context of the RamR ligand binding cavity relative to the F155 residue.

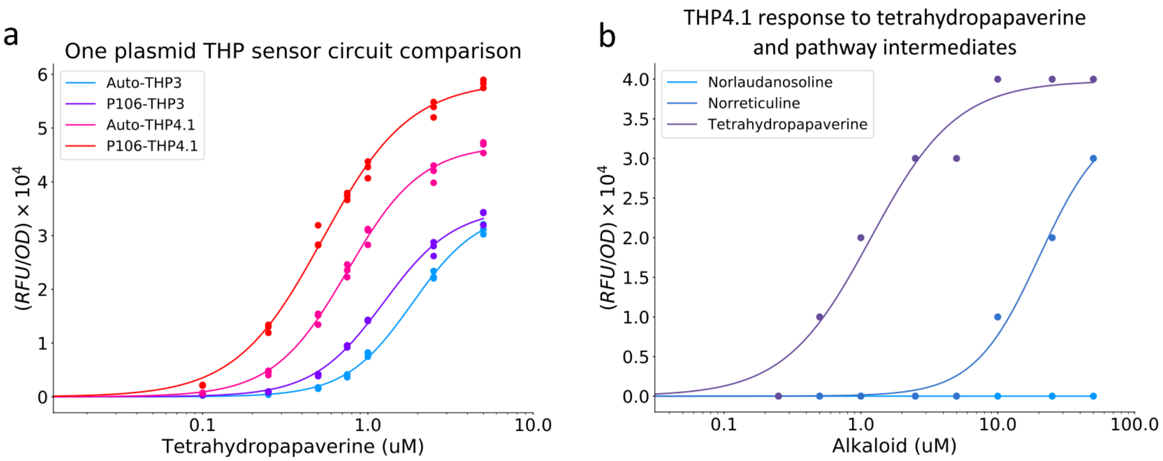


**Supplemental Figure 13.** Characterization of the THP reporter plasmid (pThpR).
(**a**) Dose response function of pThpR variants with different THP sensor variants and regulation types. “Auto” denotes that the THP sensor regulates it’s own expression whereas “P106” denotes that the THP sensor is constitutively expressed from the P106 Anderson promoter. Both THP3 and THP4.1 were compared. The P106-THP4.1 variant of pThpR was used for subsequent fluorescence-based THP measurement assays. (**b**) The dose response of pThpR (P106-THP4.1) to norlaudanosoline, norreticuline, and tetrahydropapaverine.

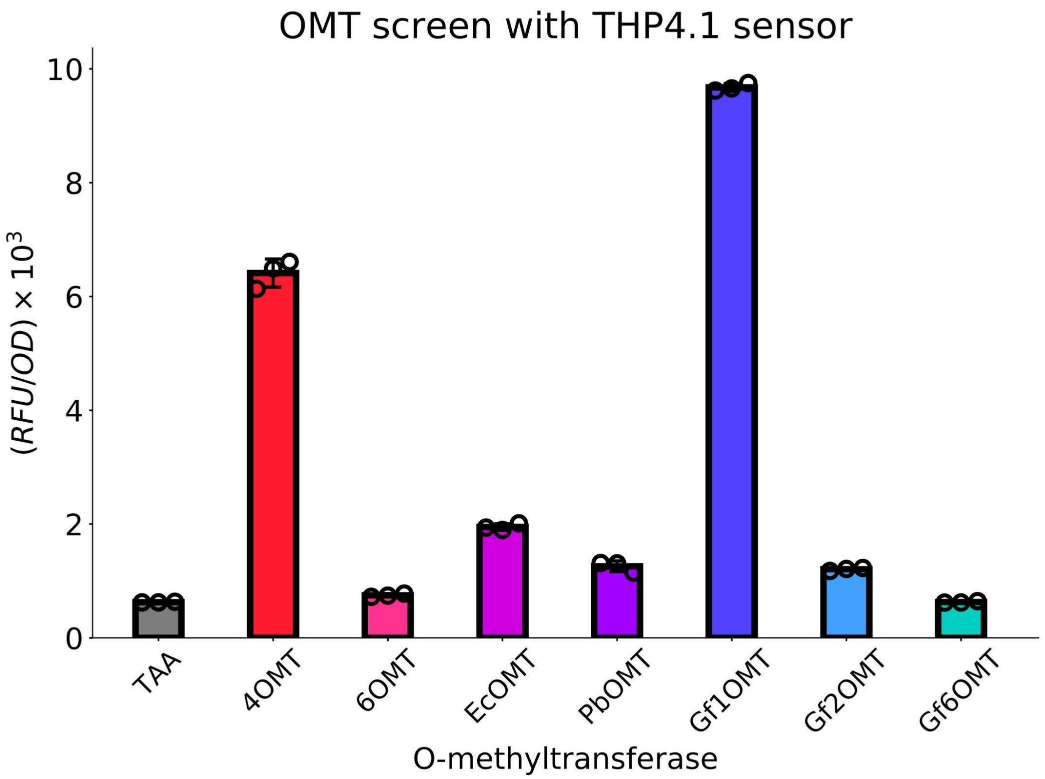


**Supplementary Figure 14**. Screening O-methyltransferases using pThpR.
Cells expressing either an empty plasmid (TAA) or a BIA methyltransferase were co-transformed with pThpR and grown in the presence of 100 uM of norlaudanosoline and 1 mg/mL ascorbic acid for 18 hours at 30C and culture fluorescence was subsequently measured. Measurements were performed in biological triplicate.

a

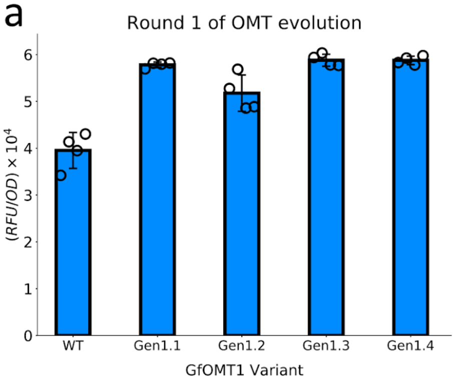


| Variant name | Gen1.1 | Gen1.2 | Gen1.3 | Gen1.4 |
|---|---|---|---|---|
| New mutations | S4P V102A T178S F189L E293V | K95R E232V E293V | **K146R T178A** | S54G I182N E350V |

b

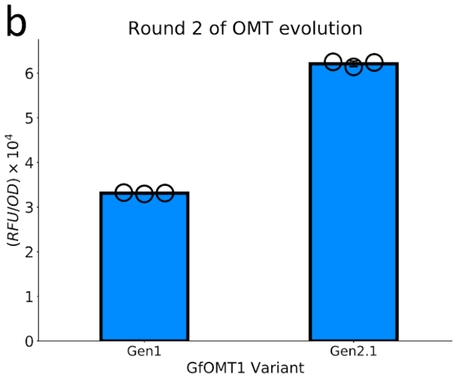


| Variant name | Gen2.1 |
|---|---|
| New mutations | **W22L** x4 |

c

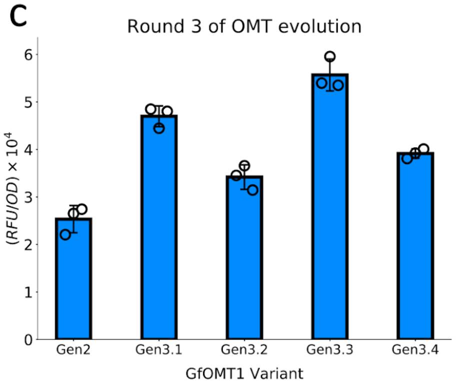


| Variant name | Gen3.1 | Gen3.2 | Gen3.3 | Gen3.4 |
|---|---|---|---|---|
| New mutations | A291V | L177M | **I258V** | V94D L177M D261N |

d

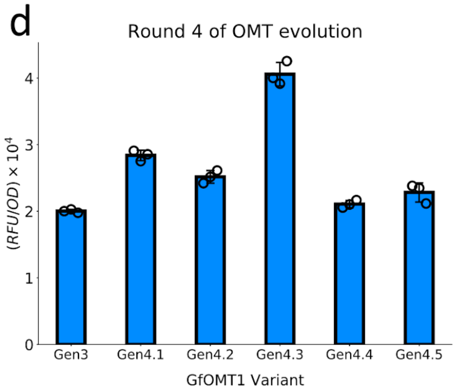


| Variant name | Gen4.1 | Gen4.1 | Gen4.1 | Gen4.1 | Gen4.1 |
|---|---|---|---|---|---|
| New mutations | A291V | L177M | **A291V L177M** | I118V D183N | S31P M121V |

e

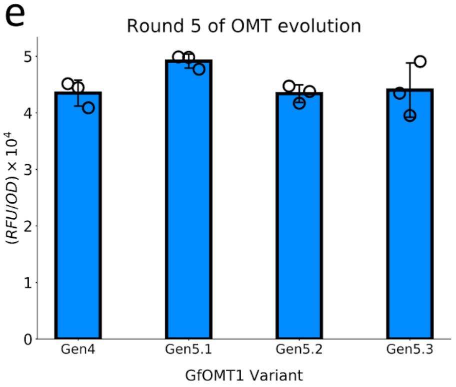


| Variant name | Gen5.1 | Gen5.2 | Gen5.3 |
|---|---|---|---|
| New mutations | **M121V** | D183N | M121V D183N |

f

| Round | Norlaudanosine used in assay |
|---|---|
| One | 100uM |
| Two | 100uM |
| Three | 100uM |
| Four | 20uM |
| Five | 10uM |

**Supplemental Figure 15.** Performance of all top OMT variants recovered from each round of evolution. (**a-e**) Fluorescent response of top unique GLAU variants using the pThpR reporter. All variants were subcloned into a fresh pReg backbone prior to characterization with the pThpR plasmid. The "x4" symbol denotes that this amino acid sequence was recovered four times following evolution. The variant genotype highlighted in green was chosen as the template for the following round of evolution. Measurements were performed in biological triplicate or quadruplicate. (**f**) Concentration of the norlaudanosoline substrate used to characterize the performance of evolved OMTs in (**a-e**).

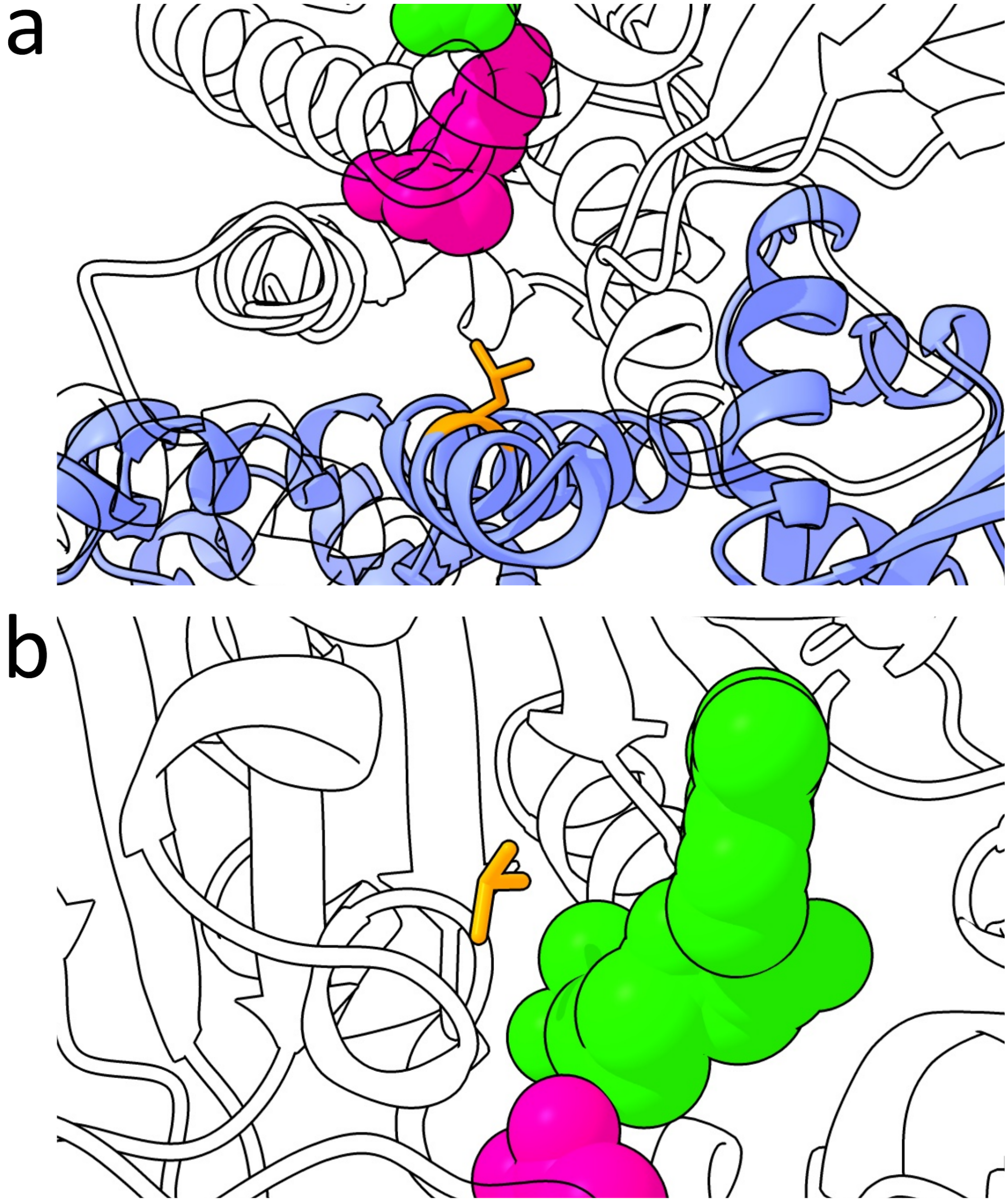


**Supplemental Figure 16**. Local environment of W22L and I258V OMT mutations.
A homology structure of the GEN5 OMT was constructed using SwissModel to infer the local environment of enzyme mutations. The substrate norlaudanosoline is shown in pink, the co-factor S-adenosyl-methionine is shown in green and mutations are shown in orange. One monomer is transparent while the other monomer is colored blue. (**a**) Environment of the W22L mutation that first appears in the GEN2 OMT variant. (**b**) Environment of the I258V mutations that first appears in the GEN3 OMT variant.

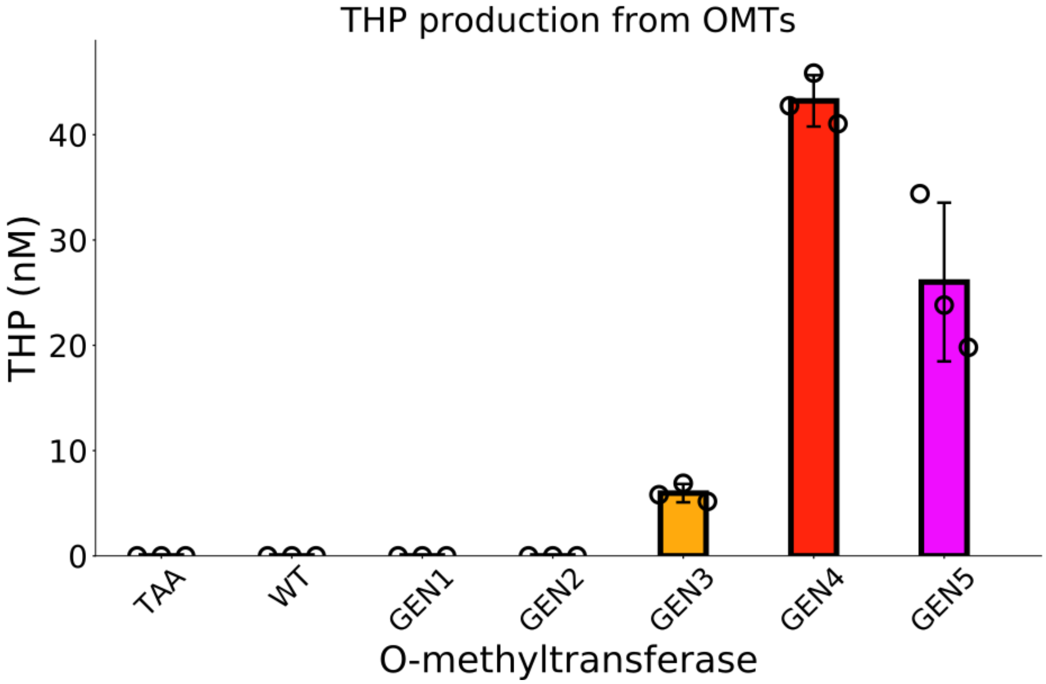


**Supplemental Figure 17.** Quantification of THP produced by each OMT variant.
Cells expressing an empty plasmid (TAA) or an OMT variant were cultured in the presence of 10 uM NOR and 1 mg/mL ascorbic acid for 18 hours at 30 C and THP was quantified, after a 1:100 fold dilution, with LC/MS by fitting to a standard curve ($R^2$ = 0.9999). Measurements were made with filtered cell supernatant and were performed in biological triplicate.

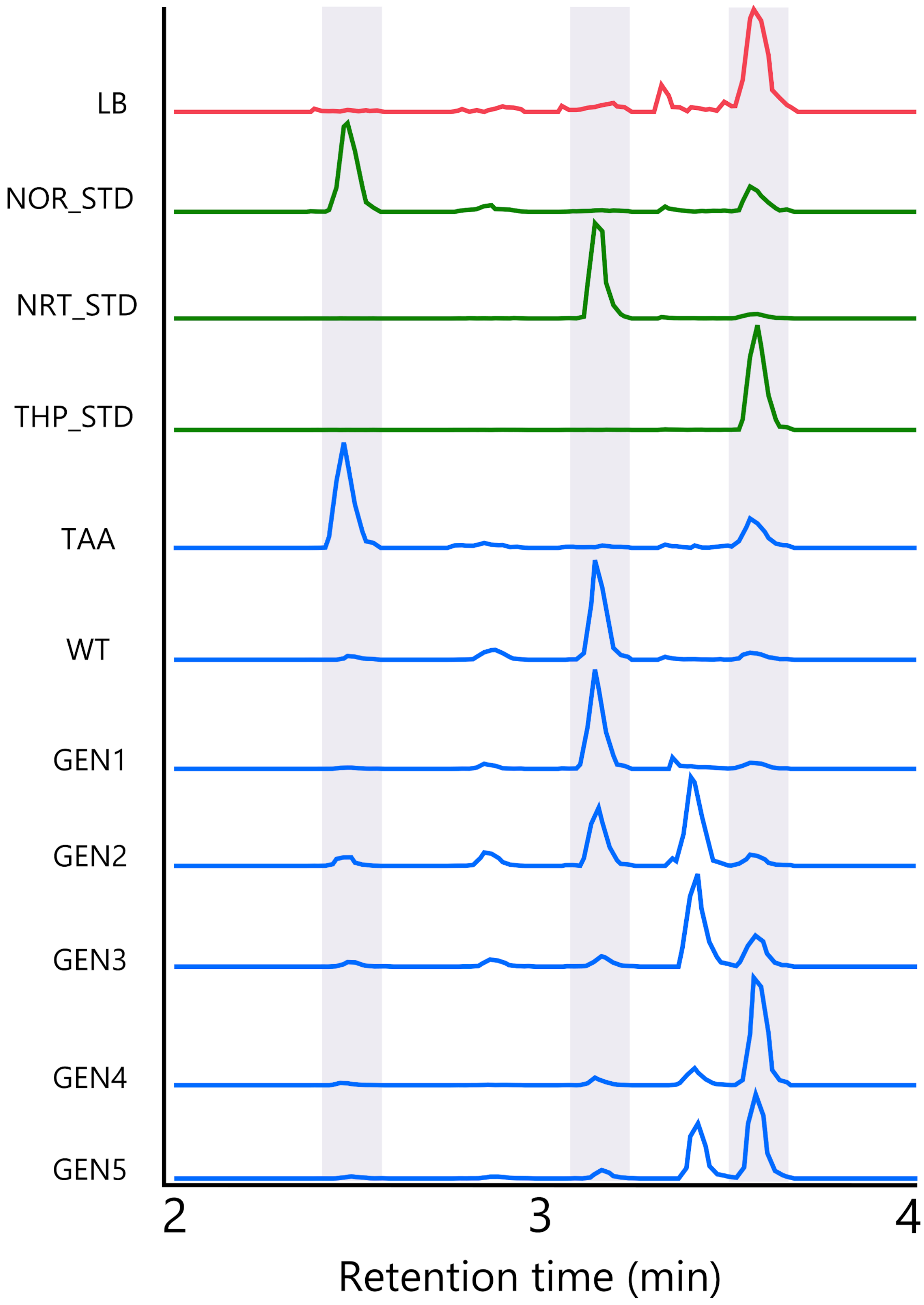


**Supplemental Figure 18**. Representative ion extracted chromatograms of samples and standards. Standards are labelled as “X_STD” and samples are from strains expressing either an empty plasmid (TAA) or enzyme variants (WT, GEN1, GEN2, GEN3, GEN4, GEN5). All LC-MS chromatograms were selected for the theoretical m/z ratios of norlaudanosoline (NOR), 6-O-Methylnorlaudanosoline, norreticuline (NRT), norlaudanine, and tetrahydropapaverine (THP) (See **Supplementary Table 3** below). Grey bars provide a reference for the expected elution times of NOR, NRT, and THP. The LB control sample displays a significant background peak at the expected retention time for THP. This peak may be caused by the tripeptides Pro-Glu-Val or Asp-Leu-Pro, which have a similar exact mass compared to THP

| Template → Sensor variant | Alkaloid used (uM) | Promoter expressing template | Libraries used |
|---|---|---|---|
| WT → GLAU1 | 100 | P103-RBS(elvj) | Y, C, G |
| GLAU1 → GLAU2 | 25 | P103-RBS(elvj) | Y, G, E |
| GLAU2 → GLAU3 | 5 | P103-RBS(elvj) | P, B, E |
| GLAU3 → GLAU4 | 1 | P114-RBS5 | P, G, E |
| WT → NOS1 | 100 | P103-RBS(elvj) | Y, C, G |
| NOS1 → NOS2 | 50 | P103-RBS(elvj) | Y, G, E |
| NOS2 → NOS3 | 25 | P106-RBS5 | P, Y, B, E |
| NOS3 → NOS4 | 2.5 | P114-RBS5 | Y, B, E |
| WT → PAP1 | 200 | P103-RBS(elvj) | Y, C, G |
| PAP1 → PAP2 | 100 | P103-RBS(elvj) | C, G, E |
| PAP2 → PAP3 | 25 | P103-RBS(elvj) | P, B, E |
| PAP3 → PAP4 | 2.5 | P103-RBS(elvj) | P, G, E |
| WT → ROTU1 | 100 | P103-RBS(elvj) | Y, C, G |
| ROTU1 → ROTU2 | 25 | P103-RBS(elvj) | Y, E, G |
| ROTU2 → ROTU3 | 10 | P106-RBS5 | P, Y, B, E |
| ROTU3 → ROTU4 | 2.5 | P114-RBS5 | P, B, E |
| WT → THP1 | 50 | P103-RBS(elvj) | Y, C, G |
| THP1 → THP2 | 5 | P103-RBS(elvj) | C, G, E |
| THP3 → THP3 | 2 | P106-RBS5 | P, B, E |
| THP3 → THP4 | 1 | P114-RBS5 | P, G, E |

**Supplementary Table 1**. Key parameters of each round of RamR evolution.
Each row indicates the round of evolution ("Template → Sensor variant"), the amount of the target BIA applied to the LB agar plate for screening ("Alkaloid used (uM)"), the promoter/RBS used to express the RamR variant template undergoing evolution ("promoter expressing template"), and the libraries used to introduce diversity ("Libraries used"). For column three, the colored box represents the relative expression level with red being strongest, orange being medium, and yellow being the weakest. For column four, letter codes represent the following (Y= yellow = T85, I88, Y92. C= cyan = K63, L66, M70. P= purple = E120, A123, D124. B = blue = L133, C134, S137. G = grey = R148, D152, L156. E = random mutagenesis)

| | PAP4 | ROTU4 | NOS4 | GLAU4 |
|---|---|---|---|---|
| **Data collection** | | | | |
| Space group | C2 | P1 | P1 | P1 |
| Cell dimensions | | | | |
| a, b, c (Å) | 106.76, 68.57, 69.57 | 46.14, 50.84, 50.83 | 41.63, 54.86, 92.64 | 43.18, 54.20, 91.52 |
| α, β, γ (°) | 90.00, 127.53, 90.00 | 120.05, 90.17, 89.96 | 74.14, 81.82, 89.96 | 104.90, 98.00, 89.99 |
| Resolution (Å) | 50.00-1.6 (1.63-1.60)* | 50.00-1.73 (1.76-1.73) | 50.00-2.21 (2.25-2.21) | 50.00 - 2.00 (2.03-2.00) |
| $R_{sym}$/ $R_{pim}$ | 0.055(0.474)/0.034(0.306) | 0.060(0.285)/0.056(0.263) | 0.072(0.354)/0.068(0.341) | 0.061(0.320)/0.056(0.300) |
| CC ½ $^{\Upsilon}$ | 0.947 (0.751) | 0.936 (0.830) | 0.921 (0.76) | 0.943 (0.813) |
| I / σ | 21.3 (1.7) | 16.6 (1.9) | 12.1 (1.6) | 14.7 (1.7) |
| Completeness (%) | 99.8 (99.5) | 96.0 (95.1) | 95.0 (96.2) | 94.3 (84.2) |
| Redundancy | 3.5 (3.2) | 1.9 (1.8) | 1.8 (1.8) | 1.8 (1.5) |
| **Refinement** | | | | |
| Resolution (Å) | 48.82-1.60 (1.66-1.60) | 46.14-1.74 (1.80-1.74) | 44.07 - 2.21 (2.29 - 2.21) | 43.76 - 2.00 (2.07 - 2.00) |
| No. reflections | 52242 (5094) | 39481 (3649) | 36872 (3591) | 50140 (4375) |
| $R_{work}$ | 0.1868 (0.2380) | 0.2162 (0.2855) | 0.2418 (0.2837) | 0.2022 (0.2552) |
| $R_{free}^{\pm}$ | 0.2108 (0.2635) | 0.2577 (0.2714) | 0.2849 (0.3413) | 0.2334 (0.2938) |
| **No. atoms** | 3239 | 3140 | 5950 | 6345 |
| Protein | 2935 | 2842 | 5657 | 5865 |
| Ligand/ion | 60 | 62 | 140 | 124 |
| Water | 244 | 236 | 153 | 356 |
| **B-factors (Å²)** | | | | |
| Protein | 25.4 | 23.3 | 51.7 | 42.9 |
| Ligand/ion | 16.8 | 21.2 | 53.4 | 31.7 |
| Water | 38.6 | 31.7 | 48.4 | 45.1 |
| **R.m.s. deviations** | | | | |

| | | | | |
|---|---|---|---|---|
| Bond lengths (Å) | 0.006 | 0.001 | 0.01 | 0.008 |
| Bond angles (°) | 0.72 | 0.48 | 0.84 | 0.75 |
| **Ramachandran plot** | | | | |
| Favored | 99.45% | 99.14% | 99.15% | 99.31% |
| Allowed | 0.55% | 0.86% | 0.85% | 0.69% |
| Outliers | 0.00% | 0.00% | 0.00% | 0.00% |
| **Molprobity score^** | 1.22 / 98th percentile | 1.50 / 92nd percentile | 1.81 / 93rd percentile | 1.53 / 96th percentile |

*Values for the corresponding parameters in the outermost shell in parenthesis.
ϒ$CC_{1/2}$ is the Pearson correlation coefficient for a random half of the data, the two numbers represent the lowest and highest resolution shell, respectively.
±$R_{free}$ is the $R_{work}$ calculated for about 10% of the reflections randomly selected and omitted from refinement.
^MolProbity score is calculated by combining clashscore with rotamer and Ramachandran percentage and scaled based on X-ray resolution. The percentage is calculated with 100th percentile as the best and 0th percentile as the worst among structures of comparable resolution.

**Supplementary Table 2**. X-ray Crystallography Data Collection and Refinement Statistics

| | Observed EIC Areas | | | | |
|---|---|---|---|---|---|
| **Sample** | **NOR-4OH (288.1230 m/z)** | **3OH (302.1387 m/z)** | **NRT-2OH (316.1543 m/z)** | **1OH (330.1700 m/z)** | **THP-0OH (344.1856 m/z)** |
| **Retention Time (min.)** | **2.46** | **2.85** | **3.14** | **3.41** | **3.58** |
| 1(TAA_1) | 1,900,714.32 | 0.00 | 0.00 | 0.00 | 732,062.09 |
| 2(TAA_2) | 1,915,746.57 | 0.00 | 0.00 | 0.00 | 739,719.41 |
| 3(TAA_3) | 2,429,528.49 | 0.00 | 0.00 | 0.00 | 683,951.95 |
| 4(WT_1) | 292,886.91 | 838,007.19 | 5,016,094.31 | 0.00 | 715,848.03 |
| 5(WT_2) | 183,019.07 | 678,233.01 | 4,455,448.19 | 0.00 | 726,021.76 |
| 6(WT_3) | 553,463.54 | 956,624.78 | 5,512,157.20 | 0.00 | 724,935.11 |
| 7(Gen1_1) | 127,968.53 | 262,284.74 | 5,028,466.49 | 0.00 | 662,206.48 |
| 8(Gen1_2) | 189,597.68 | 357,195.05 | 5,328,381.15 | 0.00 | 685,462.09 |
| 9(Gen1_3) | 69,624.59 | 194,528.60 | 4,578,048.19 | 0.00 | 627,833.84 |
| 10(Gen2_1) | 641,245.71 | 730,076.27 | 2,721,068.90 | 4,013,082.04 | 768,144.56 |
| 11(Gen2_2) | 251,793.95 | 485,465.06 | 1,499,029.94 | 4,748,195.68 | 825,357.07 |
| 12(Gen2_3) | 457,481.05 | 582,826.88 | 2,227,163.03 | 4,367,581.54 | 796,201.94 |
| 13(Gen3_1) | 304,453.82 | 474,632.34 | 603,324.03 | 6,336,545.51 | 2,220,655.54 |
| 14(Gen3_2) | 259,191.32 | 327,542.61 | 565,407.61 | 6,114,959.98 | 2,469,776.82 |
| 15(Gen3_3) | 388,680.89 | 444,919.64 | 622,087.57 | 6,641,956.94 | 2,075,163.88 |
| 16(Gen4_1) | 242,876.32 | 0.00 | 639,571.41 | 1,994,412.63 | 10,694,641.10 |
| 17(Gen4_2) | 139,427.97 | 0.00 | 488,824.56 | 1,353,523.79 | 11,406,507.42 |
| 18(Gen4_3) | 249,690.78 | 0.00 | 684,700.15 | 2,054,059.80 | 10,304,271.54 |
| 19(Gen5_1) | 33,523.35 | 55,140.14 | 377,855.13 | 3,057,004.77 | 8,779,382.47 |
| 20(Gen5_2) | 159,548.08 | 235,055.43 | 565,294.52 | 4,882,357.32 | 6,352,723.48 |
| 21(Gen5_3) | 385,426.52 | 238,451.26 | 608,492.27 | 5,854,903.52 | 5,431,738.76 |
| 22(NOR_50nM_STD) | 1,674,119.68 | 0.00 | 0.00 | 0.00 | 650,085.33 |
| 23(NRT_50nM_STD) | 0.00 | 0.00 | 6,704,519.34 | 0.00 | 659,703.80 |
| 24(THP_10nM_STD) | 0.00 | 0.00 | 0.00 | 0.00 | 3,178,252.46 |
| 25(THP_50nM_STD) | 0.00 | 0.00 | 0.00 | 0.00 | 11,969,036.43 |

*THP overlaps with matrix peaks, so area never actually goes to zero

**Supplemental Table 3**. Observed extracted ion chromatograms areas for all controls, standards, and enzymatic reactions. The top two rows indicate the m/z ratio and retention time for compounds of interest (norlaudanosoline (NOR-4OH), 6-O-Methylnorlaudanosoline (3OH), norreticuline (NRT-2OH), norlaudanine (1OH), and tetrahydropapaverine (THP-0OH). All sample measurements were performed in biological triplicate.

**> pReg**

GAAGCTAGAGTAAGTAGTTCGCCAGTTAATAGTTTGCGCAACGTTGTTGCCATTGCTGCAGGCATCG
TGGTGTCACGCTCGTCGTTTGGTATGGCTTCATTCAGCTCCGGTTCCCAACGATCAAGGCGAGTTAC
ATGATCCCCCATGTTGTGCAAAAAAGCGGTTAGCTCCTTCGGTCCTCCGATCGTTGTCAGAAGTAAG
TTGGCCGCAGTGTTATCACTCATGGTTATGGCAGCACTGCATAATTCTCTTACTGTCATGCCATCCGT
AAGATGCTTTTCTGTGACTGGTGAGTACTCAACCAAGTCATTCTGAGAATAGTGTATGCGGCGACCG
AGTTGCTCTTGCCCGGCGTCAACACGGGATAATACCGCGCCACATAGCAGAACTTTAAAAGTGCTCA
TCATTGGAAAACGTTCTTCGGGGCGAAAACTCTCAAGGATCTTACCGCTGTTGAGATCCAGTTCGAT
GTAACCCACTCGTGCACCCAACTGATCTTCAGCATCTTTTACTTTCACCAGCGTTTCTGGGTGAGCA
AAAACAGGAAGGCAAAATGCCGCAAAAAAGGGAATAAGGGCGACACGGAAATGTTGAATACTCAT
ACTCTTCCTTTTTCAATATTATTGAAGCATTTATCAGGGTTATTGTCTCATGAGCGGATACATATTTGA
ATGTATTTAGAAAAATATGCGCCTTGAGCGACACGAATTATGCAGTGATTTACGACCTGCACAGCCAT
ACCACAGCTTCCGATGGCTGCCTGACGCCAGAAGCATTGGTGCACCGTGCAGTCGATGATAAGCTG
TCAAACATGAGAATTGTGCCTAATGAGTGAGCTAACTTACATTAATTGCGTTGCGCTCACTGCCCGC
TTTCCAGTCGGGAAACCTGTCGTGCCAGCTGCATTAATGAATCGGCCAACGCGCGGGGAGAGGCGG
TTTGCGTATTGGGCGCCAGGGTGGTTTTTCTTTTCACCAGTGAGACGGGCAACAGCTGATTGCCCTT
CACCGCCTGGCCCTGAGAGAGTTGCAGCAAGCGGTCCACGCTGGTTTGCCCCAGCAGGCGAAAAT
CCTGTTTGATGGTGGTTAACGGCGGGATATAACATGAGCTATCTTCGGTATCGTCGTATCCCACTACC
GAGATATCCGCACCAACGCGCAGCCCGGACTCGGTAATGGCGCGCATTGCGCCCAGCGCCATCTGA
TCGTTGGCAACCAGCATCGCAGTGGGAACGATGCCCTCATTCAGCATTTGCATGGTTTGTTGAAAAC
CGGACATGGCACTCCAGTCGCCTTCCCGTTCCGCTATCGGCTGAATTTGATTGCGAGTGAGATATTTA
TGCCAGCCAGCCAGACGCAGACGCGCCGAGACAGAACTTAATGGGCCCGCTAACAGCGCGATTTGC
TGGTGACCCAATGCGACCAGATGCTCCACGCCCAGTCGCGTACCATCTTCATGGGAGAAAATAATAC
TGTTGATGGGTGTCTGGTCAGAGACATCAAGAAATAACGCCGGAACATTAGTGCAGGCAGCTTCCA
CAGCAATGGCATCCTGGTCATCCAGCGGATAGTTAATGATCAGCCCACTGACGCGTTGCGCGAGAAG
ATTGTGCACCGCCGCTTTACAGGCTTCGACGCCGCTTCGTTCTACCATCGACACCACCACGCTGGCA
CCCAGTTGATCGGCGCGAGATTTAATCGCCGCGACAATTTGCGACGGCGCGTGCAGGGCCAGACTG
GAGGTGGCAACGCCAATCAGCAACGACTGTTTGCCCGCCAGTTGTTGTGCCACGCGGTTGGGAATG
TAATTCAGCTCCGCCATCGCCGCTTCCACTTTTTCCCGCGTTTTCGCAGAAACGTGGCTGGCCTGGT
TCACCACGCGGGAAACGGTCTGATAAGAGACACCGGCATACTCTGCGACATCGTATAACGTTACTGG
TTTCACATTCACCACCCTGAATTGACTCTCTTCCGGGCGCTATCATGCCATACCGCGAAAGGTTTTGC
ACCATTCGATGGTGTCGGGACGTCAGGTGGCACTTTTCGGGGAAATGTGCGCGGAACCCCTATTTG
TTGCGGCCGCGAAGACAGCGTTATCAGAGATGAGACACCGTGCGATAATGTCGGGCAATCAGGTGC
GACTCGGTACCAAATTCCAGAAAAGAGGGGAGCGGGAAACCGCTCCCCTTTTTCGTTTTGGTCCC
AATCTATCGATTGTATGGACTTCATCTTCATTACCTCGTATCATTGTACACCTGCCGAACGCAAGGGC
ATGGGCTGTCGACCTTTGAAAAGTACCCTGATAGCTAGCTCAGTCCTAGGGATTATGCTAGCAATTAC
GAGCCCCATAGGGTGGTGTGTACCACCCCTGATGAGTCCAAAAGGACGAAATGGGGCCTCTACAAA
TAATTTTGTTTAACGGAACCACGTATCAGAAGGAGGTTAGTATATGGTTGCTCGCCCAAAGTCTGAG
GACAAAAAGCAGGCATTGCTTGAAGCGGCAACTCAAGCCATCGCGCAATCAGGCATTGCCGCTAGT
ACCGCTGTAATTGCACGCAATGCGGGAGTTGCGGAAGGGACGTTGTTCCGCTATTTCGCAACGAAA
GATGAGTTGATCAACACCCTTTACTTACATTTGAAACAGGACCTGTGCCAATCAATGATCATGGAATT
GGATCGTTCTATTACTGACGCTAAGATGATGACCCGTTTTATCTGGAACAGTTATATTAGCTGGGGATT
GAACCACCCAGCTCGCCATCGTGCCATTCGTCAGTTGGCGGTTTCTGAAAAGTTGACGAAGGAAAC
CGAACAACGCGCGGATGATATGTTCCCGGAGTTACGCGACTTGTGCCACCGTAGTGTTCTTATGGTG
TTTATGTCCGACGAGTACCGCGCCTTCGGCGACGGGTTGTTCTTGGCGCTTGCTGAGACGACTATGG
ATTTCGCTGCGCGCGACCCGGCTCGCGCTGGTGAGTACATTGCGTTGGGCTTCGAGGCTATGTGGC
GCGCACTTACGCGCGAAGAGCAGTAATAATCCTAAGAATTCGAGGAGTGCAGGCTCGGTAACATAC
GGTCTAGCTATCTGACTATCGCCGCTGTGAGCTCGGTACCAAATTCCAGAAAAGAGGCCGCGAAAG
CGGCCTTTTTTCGTTTTGGTCCAAGCCCGATGCGCCAGAGTTGTTTCTGAAACATGGCAAAGGTATC
ACTAGTCTTCGCGGCCGCCATGCTGTCCAGGCAGGTAGATGACGACCATCAGGGACAGCTTCAAGG
ATCGCTCGCGGCTCTTACCAGCCTAACTTCGATCATTGGACCGCTGATCGTCACGGCGATTTATGCCG
CCTCGGCGAGCACATGGAACGGGTTGGCATGGATTGTAGGCGCCGCCCTATACCTTGTCTGCCTCCC
CGCGTTGCGTCGCGGTGCATGGAGCCGGGCCACCTCGACCTGAATGGAAGCCGGCGGCACCTCGC
TAACGGATTCACCACTCCAAGAATTGGAGCCAATCAATTCTTGCGGAGAACTGTGAATGCGCAAAC
CAACCCTTGGCAGAACATATCCATCGCGTCCGCCATCTCCAGCAGCCGCACGCGGCGCATCTCGGG
CAGCGTTGGGTCCTGGCCACGGGTGCGCATGATCGTGCTCCTGTCGTTGAGGACCCGGCTAGGCTG
GCGGGGTTGCCTTACTGGTTAGCAGAATGAATCACCGATACGCGAGCGAACGTGAAGCGACTGCTG
CTGCAAAACGTCTGCGACCTGAGCAACAACATGAATGGTCATCGGTTTCCGTGTTTCGTAAAGTCT

GGAAACGCGGAAGTCAGCGCCCTGCACCATTATGTTCCGGATCTGCATCGCAGGATGCTGCTGGCT
ACCCTGTGGAACACCTACATCTGTATTAACGAAGCGCTGGCATTGACCCTGAGTGATTTTTCTCTGG
TCCCGCCGCATCCATACCGCCAGTTGTTTACCCTCACAACGTTCCAGTAACCGGGCATGTTCATCATC
AGTAACCCGTATCGTGAGCATCCTCTCTCGTTTCATCGGTATCATTACCCCCATGAACAGAAATCCCC
CTTACACGGAGGCATCAGTGACCAAACAGGAAAAAACCGCCCTTAACATGGCCCGCTTTATCAGAA
GCCAGACATTAACGCTTCTGGAGAAACTCAACGAGCTGGACGCGGATGAACAGGCAGACATCTGTG
AATCGCTTCACGACCACGCTGATGAGCTTTACCGCAGCTGCCTCGCGCGTTTCGGTGATGACGGTG
AAAACCTCTGACACATGCAGCTCCCGGAGACGGTCACAGCTTGTCTGTAAGCGGATGCCGGGAGCA
GACAAGCCCGTCAGGGCGCGTCAGCGGGTGTTGGCGGGTGTCGGGGCGCAGCCATGACCCAGTCA
CGTAGCGATAGCGGAGTGTATACTGGCTTAACTATGCGGCATCAGAGCAGATTGTACTGAGAGTGCA
CCGGTGTGAAATACCGCACAGATGCGTAAGGAGAAAATACCGCATCAGGCGCTCTTCCGCTTCCTC
GCTCACTGACTCGCTGCGCTCGGTCGTTCGGCTGCGGCGAGCGGTATCAGCTCACTCAAAGGCGGT
AATACGGTTATCCACAGAATCAGGGGATAACGCAGGAAAGAACATGTGAGCAAAAGGCCAGCAAA
AGGCCAGGAACCGTAAAAAGGCCGCGTTGCTGGCGTTTTTCCATAGGCTCCGCCCCCCTGACGAGC
ATCACAAAAATCGACGCTCAAGTCAGAGGTGGCGAAACCCGACAGGACTATAAAGATACCAGGCGT
TTCCCCCTGGAAGCTCCCTCGTGCGCTCTCCTGTTCCGACCCTGCCGCTTACCGGATACCTGTCCGC
CTTTCTCCCTTCGGGAAGCGTGGCGCTTTCTCATAGCTCACGCTGTAGGTATCTCAGTTCGGTGTAG
GTCGTTCGCTCCAAGCTGGGCTGTGTGCACGAACCCCCCGTTCAGCCCGACCGCTGCGCCTTATCC
GGTAACTATCGTCTTGAGTCCAACCCGGTAAGACACGACTTATCGCCACTGGCAGCAGCCACTGGTA
ACAGGATTAGCAGAGCGAGGTATGTAGGCGGTGCTACAGAGTTCTTGAAGTGGTGGCCTAACTACG
GCTACACTAGAAGGACAGTATTTGGTATCTGCGCTCTGCTGAAGCCAGTTACCTTCGGAAAAAGAGT
TGGTAGCTCTTGATCCGGCAAACAAACCACCGCTGGTAGCGGTGGTTTTTTTGTTTGCAAGCAGCA
GATTACGCGCAGAAAAAAAGGATCTCAAGAAGATCCTTTGATCTTTTCTACGGGGTCTGACGCTCAG
TGGAACGAAAACTCACGTTAAGGGATTTTGGTCATGAGATTATCAAAAAGGATCTTCACCTAGATCC
TTTTAAATTAAAAATGAAGTTTTAAATCAATCTAAAGTATATATGAGTAAACTTGGTCTGACAGTTAC
CAATGCTTAATCAGTGAGGCACCTATCTCAGCGATCTGTCTATTTCGTTCATCCATAGTTGCCTGACT
CCCCGTCGTGTAGATAACTACGATACGGGAGGGCTTACCATCTGGCCCCAGTGCTGCAATGATACCG
CGCGATCCACGCTCACCGGCTCCAGATTTATCAGCAATAAACCAGCCAGCCGGAAGGGCCGAGCGC
AGAAGTGGTCCTGCAACTTTATCCGCCTCCATCCAGTCTATTAATTGTTGCCGG

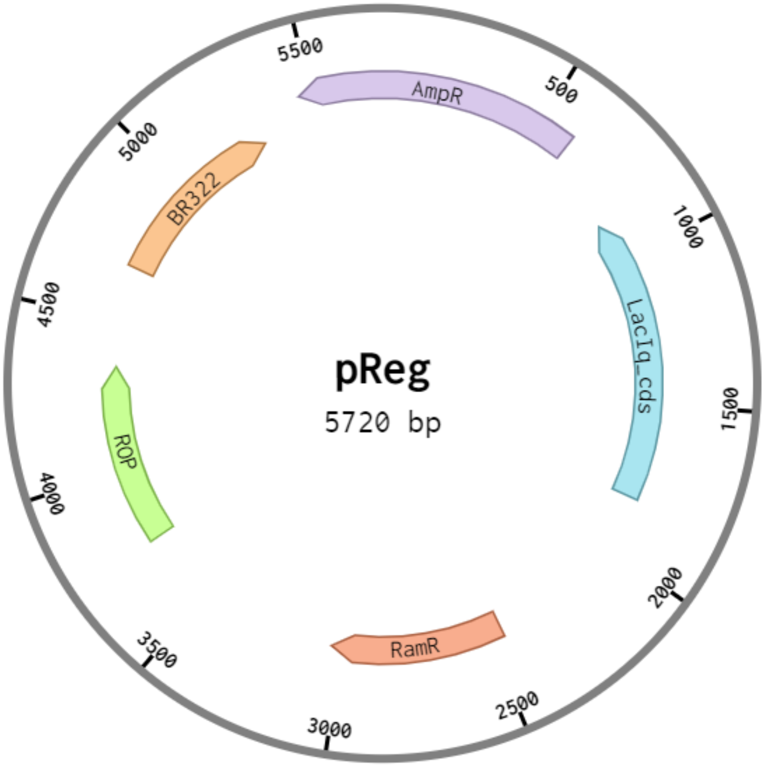


**> pGFP**

CGAGCATACTATCACGTCGGCGACCACTAGTCAGTTAACGCAAGGGCATGGGCTGTCGACCTTTGAA

AAGTACCTTGACGGCGTATCTTTGCTTTCTATAATGAGTGCTTACTCACTCATACAATAGTCAGTCATA
AGTCTGGGCTAAGCCCACTGATGAGTCGCTGAAATGCGACGAAACTTATGACCTCTACAAATAATTT
TGTTTAACGTAAACCTCCGGGTTAATAAGGAGTAATTATGGCATCCAAGGGCGAGGAGCTCTTTACT
GGCGTAGTACCAATTCTCGTAGAGCTCGATGGCGATGTAAATGGCCATAAGTTTTCCGTACGCGGCG
AGGGCGAGGGCGATGCAACTAACGGCAAGCTCACTCTCAAGTTTATTTGTACTACTGGCAAGCTCCC
AGTACCATGGCCAACTCTCGTAACTACTCTGACCTATGGCGTACAATGTTTTTCCCGCTATCCAGATC
ACATGAAGCAACATGATTTTTTTAAGTCCGCAATGCCAGAGGGCTATGTACAAGAGCGCACTATTAG
CTTTAAGGATGATGGCACCTATAAGACTCGCGCAGAGGTAAAGTTTGAGGGCGATACTCTCGTAAAT
CGCATTGAGCTCAAGGGCATTGATTTTAAGGAGGATGGCAATATTCTCGGCCATAAGCTGGAGTATA
ATTTCAATTCCCATAATGTATATATTACCGCAGATAAGCAAAAGAATGGCATTAAGGCGAATTTTAAG
ATTCGCCATAATGTGGAGGATGGCTCCGTACAACTCGCAGATCATTATCAACAAAATACTCCAATTGG
CGATGGCCCAGTACTCCTCCCAGATAATCATTATCTCTCCACTCAATCCGTGCTCTCCAAAGATCCAA
ATGAGAAGCGCGATCACATGGTACTCCTGGAGTTTGTAACTGCAGCAGGCATTACTCATGGCATGGA
TGAGCTCTATAAGCTCGAGCACCACCACCACCACCACTGATAATCCAAACCTGTTATATGTTAGCTGA
GACTAGTTGGAAGTGTGGCTGTCCTCAAGCGTTTTAGTTCGTCGGTCAGTTTCACCTGATTTACGTA
AAAACCCGCTTCGGCGGGTTTTTGCTTTTGGAGGGGCAGAAAGATGAATGACTGTCGGCCATTCGA
TGGTGTCGGGTAGCATAACCCCTTGTGATAGTCTTCGCGGCCGCTCACACTGCTTCCGGTAGTCAAT
AAACCGGTAAACCAGCAATAGACATAAGCGGCTATTTAACGACCCTGCCCTGAACCGACGACCGGG
TCGAATTTGCTTTCGAATTTCTGCCATTCATCCGCTTATTATCACTTATTCAGGCGTAGCAACCAGGC
GTTTAAGGGCACCAATAACTGCCTTAAAAAAATTACGCCCCGCCCTGCCACTCATCGCAGTACTGTT
GTAATTCATTAAGCATTCTGCCGACATGGAAGCCATCACAAACGGCATGATGAACCTGAATCGCCAG
CGGCATCAGCACCTTGTCGCCTTGCGTATAATATTTGCCCATAGTGAAAACGGGGGCGAAGAAGTTG
TCCATATTGGCCACGTTTAAATCAAAACTGGTGAAACTCACCCAGGGATTGGCTGAGACGAAAAAC
ATATTCTCAATAAACCCTTTAGGGAAATAGGCCAGGTTTTCACCGTAACACGCCACATCTTGCGAATA
TATGTGTAGAAACTGCCGGAAATCGTCGTGGTATTCACTCCAGAGCGATGAAAACGTTTCAGTTTGC
TCATGGAAAACGGTGTAACAAGGGTGAACACTATCCCATATCACCAGCTCACCGTCTTTCATTGCCA
TACGGAACTCCGGATGAGCATTCATCAGGCGGGCAAGAATGTGAATAAAGGCCGGATAAAACTTGT
GCTTATTTTTCTTTACGGTCTTTAAAAAGGCCGTAATATCCAGCTGAACGGTCTGGTTATAGGTACATT
GAGCAACTGACTGAAATGCCTCAAAATGTTCTTTACGATGCCATTGGGATATATCAACGGTGGTATAT
CCAGTGATTTTTTTCTCCATTTTAGCTTCCTTAGCTCCTGAAAATCTCGATAACTCAAAAAATACGCC
CGGTAGTGATCTTATTTCATTATGGTGAAAGTTGGAACCTCTTACGTGCCGATCACGTCTCATTTTCG
CCAAAGTTGGCCAGGGCTTCCCGGTATCAACAGGGACACCAGGATTTATTTATNNTGCGAAGTGATC
TTCCGTCACAGGTATTTATTCGGCGCAAAGTGCGTCGGGTGATGCTGCCAACTTACTGATTTAGTGTA
TGATGGTGTTTTTGAGGTGCTCCAGTGGCTTCTGTTTCTATCAGCTGTCCCTCCTGTTCAGCTACTGA
CGGGGTGGTGCGTAACGGCAAAAGCACCGCCGGACATCAGCGCTAGCGGAGTGTATACTGGCTTAC
TATGTTGGCACTGATGAGGGTGTCAGTGAAGTGCTTCATGTGGCAGGAGAAAAAAGGCTGCACCGG
TGCGTCAGCAGAATATGTGATACAGGATATATTCCGCTTCCTCGCTCACTGACTCGCTACGCTCGGTC
GTTCGACTGCGGCGAGCGGAAATGGCTTACGAACGGGGCGGAGATTTCCTGGAAGATGCCAGGAA
GATACTTAACAGGGAAGTGAGAGGGCCGCGGCAAAGCCGTTTTTCCATAGGCTCCGCCCCCCTGAC
AAGCATCACGAAATCTGACGCTCAAATCAGTGGTGGCGAAACCCGACAGGACTATAAAGATACCAG
GCGTTTCCCCCTGGCGGCTCCCTCGTGCGCTCTCCTGTTCCTGCCTTTCGGTTTACCGGTGTCATTC
CGCTGTTATGGCCGCGTTTGTCTCATTCCACGCCTGACACTCAGTTCCGGGTAGGCAGTTCGCTCCA
AGCTGGACTGTATGCACGAACCCCCCGTTCAGTCCGACCGCTGCGCCTTATCCGGTAACTATCGTCT
TGAGTCCAACCCGGAAAGACATGCAAAAGCACCACTGGCAGCAGCCACTGGTAATTGATTTAGAGG
AGTTAGTCTTGAAGTCATGCGCCGGTTAAGGCTAAACTGAAAGGACAAGTTTTGGTGACTGCGCTC
CTCCAAGCCAGTTACCTCGGTTCAAAGAGTTGGTAGCTCAGAGAACCTTCGAAAAACCGCCCTGCA
AGGCGGTTTTTTCGTTTTCAGAGCAAGAGATTACGCGCAGACCAAAACGATCTCAAGAAGATCATCT
TATTAATCAGATAAAATATTTCTAGATTTCAGTGCAATTTATCTCTTCAAATGTAGCACCTGAAGTCAG
CCCCATACGATATAAGTTGTAATTCTCATGTTAGTCATGCCCCGCGCCCACCGGAAGGAGCTGACTG
GGTTGAAGGCTCTCAAGGGCATCGGTCGAGATCCCGGTGCCTAATGAGTGAGCTAACTTACATTAAT
TGCGTTGCGCTCACTGCCCGCTTTCCAGTCGGGAAACCTGTCGTGCCAGCTGCATTAATGAATCGGC
CAACGCGCGGGGAGAGGCGGTTTGCGTATTGGGCGCCAGGGTGGTTTTTCTTTTCACCAGTGAGAC
GGGCAACAGCTGATTGCCCTTCACCGCCTGGCCCTGAGAGAGTTGCAGCAAGCGGTCCACGCTGGT
TTGCCCCAGCAGGCGAAAATCCTGTTTGATGGTGGTTAACGGCGGGATATAACATGAGCTATCTTCG
GTATCGTCGTATCCCACTACCGAGATATCCGCACCAACGCGCAGCCCGGACTCGGTAATAGCGCGCA
TTGCGCCCAGCGCCATCTGATCGTTGGCAACCAGCATCGCAGTGGGAACGATGCCCTCATTCAGCAT
TTGCATGGTTTGTTGAAAACCGGACATGGCACTCCAGTCGCCTTCCCGTTCCGCTATCGGCTGAATT
TGATTGCGAGTGAGATATTTATGCCAGCCAGCCAGACGCAGACGCGCCGAGACAGAACTTAATGGG
CCCGCTAACAGCGCGATTTGCTGGTGACCCAATGCGACCAGATGCTCCACGCCCAGTCGCGTACCAT

CTTCATGGGAGAAAATAATACTGTTGATGGGTGTCTGGTCAGAGACATCAAGAAATAACGCCGGAA
CATTAGTGCAGGCAGCTTCCACAGCAATGGCATCCTGGTCATCCAGCGGATAGTTAATGATCAGCCC
ACTGACGCGTTGCGCGAGAAGATTGTGCACCGCCGCTTTACAGGCTTCGACGCCGCTTCGTTCTACC
ATCGACACCACCACGCTGGCACCCAGTTGATCGGCGCGAGATTTAATCGCCGCGACAATTTGCGAC
GGCGCGTGCAGGGCCAGACTGGAGGTGGCAACGCCAATCAGCAACGACTGTTTGCCCGCCAGTTG
TTGTGCCACGCGGTTGGGAATGTAATTCAGCTCCGCCATCGCCGCTTCCACTTTTTCCCGCGTTTTC
GCAGAAACGTGGCTGGCCTGGTTCACCACGCGGGAAACGGTCTGATAAGAGACACCGGCATACTCT
GCGACATCGTATAACGTTACTGGTTTCACATTCACCACCCTGAATTGACTCTCTTCCGGGCGCTATCA
TGCCATACCGCGAAAGGTTTTGCGCCATTCGATGGTGTCCGGGATCTCGACGCTCTCCCTTATGCGA
CGCGGCCGCGAAGACACTGTCCTCAATGGTTCGTTGTGATGGCGGTAGGAATGTAATCGTTAATCCG
CAAATAACGTAAAAACCCGCTTCGGCGGGTTTTTTTATGGGGGGAGTTTAGGGAAAGAGCATTTGTC
ATCCCGTTGAATATGGCTCGCATCTTAT

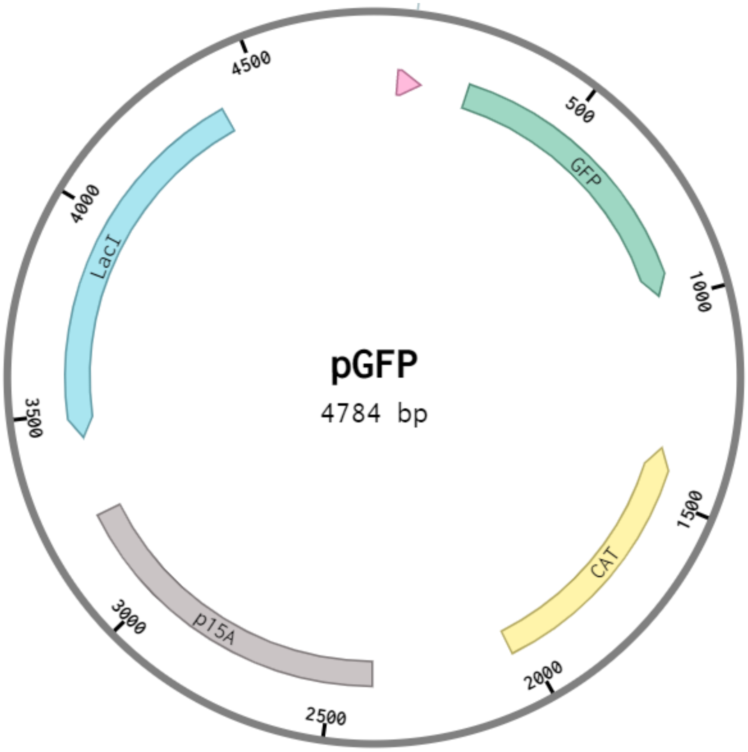


**> pGfOMT1**

TGTAGATAACTACGATACGGGAGGGCTTACCATCTGGCCCCAGTGCTGCAATGATACCGCGCGATCC
ACGCTCACCGGCTCCAGATTTATCAGCAATAAACCAGCCAGCCGGAAGGGCCGAGCGCAGAAGTGG
TCCTGCAACTTTATCCGCCTCCATCCAGTCTATTAATTGTTGCCGGGAAGCTAGAGTAAGTAGTTCGC
CAGTTAATAGTTTGCGCAACGTTGTTGCCATTGCTGCAGGCATCGTGGTGTCACGCTCGTCGTTTGG
TATGGCTTCATTCAGCTCCGGTTCCCAACGATCAAGGCGAGTTACATGATCCCCCATGTTGTGCAAA
AAAGCGGTTAGCTCCTTCGGTCCTCCGATCGTTGTCAGAAGTAAGTTGGCCGCAGTGTTATCACTCA
TGGTTATGGCAGCACTGCATAATTCTCTTACTGTCATGCCATCCGTAAGATGCTTTTCTGTGACTGGT
GAGTACTCAACCAAGTCATTCTGAGAATAGTGTATGCGGCGACCGAGTTGCTCTTGCCCGGCGTCAA
CACGGGATAATACCGCGCCACATAGCAGAACTTTAAAAGTGCTCATCATTGGAAAACGTTCTTCGGG
GCGAAAACTCTCAAGGATCTTACCGCTGTTGAGATCCAGTTCGATGTAACCCACTCGTGCACCCAAC
TGATCTTCAGCATCTTTTACTTTCACCAGCGTTTCTGGGTGAGCAAAAACAGGAAGGCAAAATGCC
GCAAAAAAGGGAATAAGGGCGACACGGAAATGTTGAATACTCATACTCTTCCTTTTTCAATATTATT
GAAGCATTTATCAGGGTTATTGTCTCATGAGCGGATACATATTTGAATGTATTTAGAAAAATATGCGCC
TTGAGCGACACGAATTATGCAGTGATTTACGACCTGCACAGCCATACCACAGCTTCCGATGGCTGCC
TGACGCCAGAAGCATTGGTGCACCGTGCAGTCGATGATAAGCTGTCAAACATGAGAATTGTGCCTA
ATGAGTGAGCTAACTTACATTAATTGCGTTGCGCTCACTGCCCGCTTTCCAGTCGGGAAACCTGTCG
TGCCAGCTGCATTAATGAATCGGCCAACGCGCGGGGAGAGGCGGTTTGCGTATTGGGCGCCAGGGT
GGTTTTTCTTTTCACCAGTGAGACGGGCAACAGCTGATTGCCCTTCACCGCCTGGCCCTGAGAGAG
TTGCAGCAAGCGGTCCACGCTGGTTTGCCCCAGCAGGCGAAAATCCTGTTTGATGGTGGTTAACGG
CGGGATATAACATGAGCTATCTTCGGTATCGTCGTATCCCACTACCGAGATATCCGCACCAACGCGCA

GCCCGGACTCGGTAATGGCGCGCATTGCGCCCAGCGCCATCTGATCGTTGGCAACCAGCATCGCAG
TGGGAACGATGCCCTCATTCAGCATTTGCATGGTTTGTTGAAAACCGGACATGGCACTCCAGTCGCC
TTCCCGTTCCGCTATCGGCTGAATTTGATTGCGAGTGAGATATTTATGCCAGCCAGCCAGACGCAGA
CGCGCCGAGACAGAACTTAATGGGCCCGCTAACAGCGCGATTTGCTGGTGACCCAATGCGACCAGA
TGCTCCACGCCCAGTCGCGTACCATCTTCATGGGAGAAAATAATACTGTTGATGGGTGTCTGGTCAG
AGACATCAAGAAATAACGCCGGAACATTAGTGCAGGCAGCTTCCACAGCAATGGCATCCTGGTCAT
CCAGCGGATAGTTAATGATCAGCCCACTGACGCGTTGCGCGAGAAGATTGTGCACCGCCGCTTTACA
GGCTTCGACGCCGCTTCGTTCTACCATCGACACCACCACGCTGGCACCCAGTTGATCGGCGCGAGAT
TTAATCGCCGCGACAATTTGCGACGGCGCGTGCAGGGCCAGACTGGAGGTGGCAACGCCAATCAGC
AACGACTGTTTGCCCGCCAGTTGTTGTGCCACGCGGTTGGGAATGTAATTCAGCTCCGCCATCGCCG
CTTCCACTTTTTCCCGCGTTTTCGCAGAAACGTGGCTGGCCTGGTTCACCACGCGGGAAACGGTCT
GATAAGAGACACCGGCATACTCTGCGACATCGTATAACGTTACTGGTTTCACATTCACCACCCTGAAT
TGACTCTCTTCCGGGCGCTATCATGCCATACCGCGAAAGGTTTTGCACCATTCGATGGTGTCGGGAC
GTCAGGTGGCACTTTTCGGGGAAATGTGCGCGGAACCCCTATTTGTTGCGGCCGCGAAGACAGCGT
TATCAGAGATGAGACACCGTGCGATAATGTCGGGCAATCAGGTGCGACTCGGTACCAAATTCCAGA
AAAGAGGGGAGCGGGAAACCGCTCCCCTTTTTCGTTTTGGTCCCAATCTATCGATTGTATGGACTT
CATCTTCATTACCTCGTATCATTGTACACCTGCCGAACGCAAGGGCATGGGCTGTCGACCTTTGAAA
AGTACCTTTATGGCTAGCTCAGTCCTAGGTACAATGCTAGCAATCAAGATACTGAGCACAGCTGTCAC
CGGATGTGCTTTCCGGTCTGATGAGTCCGTGAGGACGAAACAGCCTCTACAAATAATTTTGTTTAAA
CTAGTGAACCACGAGGCCTACATATGGGAGTTTCAGATAATAAACCAGAAAGCCAAGAGGTTGACAT
CAAAGCGCAGGCCCATCTGTGGAACATCATTTACGGGTTTGCAGATTCATTAGTATTACGCTGTGCTG
TTGAGATTGGCATTGCAGATATCATTAAGAGTAATAATGGGAGTATTAGCGTAACGGAACTTGCTTCT
AAGCTGCCCATCACTAACGTAAATAGCGACAACCTTTACCGCGTACTTCGCTACCTTGTGCATATGGG
AATTTTGAAGGAAGTCAGTGATTCAAATGAAGTCAAACTGTATTCTCTTCAGCCAGTCGCAACTCTG
CTGCTTCGTGATGCTGAGCGTTCCATGGTCCCAATTATCCTGGGGATGACCCAGAAGGACTTTATGA
TTCCCTGGCATTTCATGAAGGAAGGTTTGGGGAATGACACAACGGCCTTCGAAAAGGGGATGGGCA
TGACAATCTGGCAATACTTGGAAGGACATCCCGAGCAAAGCAATTTGTTTAACGAAGGAATGGCTG
GTGAGACTCGCCTTTTGACTAAGTCTTTAATCGATGGCTGTCGTGATACCTTCGAGGGCCTGACAAG
CCTTTGCGATGTCGGCGGGGGTAATGGGACAACAATTAAGGGCATCTACGACGCATTCCCTCAGATC
AAGTGTTCCGTCTATGATTTGCCACATGTAATCGCTAGTTCCCCCGAGCACCCGAACATCGAACGCA
TTCCAGGTGACATGTTCAAGTCAGTTCCAAGTGCCCAGGCCATCTTATTGAAACTGATTTTGCATGA
CTGGACGGACGAAGAATGCGTCAACATCTTGATCAAATGCCGCGAAGCTGTACCCAAGGATACCGG
TAAAGTCATTATTGTGGACGTGGCTCTTGAGGAAGAGAGTCAACATGAGTTGACCAAGACTCGTCTT
ATCCTTGATATTGATATGCTGGTTAATACTGGAGGTCGCGAACGTTCCGAAGATGACTGGGAGAAGC
TGTTAAAGCGTGCCGGTTTTCGTGGGCATAAGATTCGTCACATCGCGGCCATCCAAAGTGTAATCGA
AGCCTTTCCGTAATAATCCTAAGAATTCGAGGAGTGCAGGCTCGGTAACATACGGTCTAGCTATCTG
ACTATCGCCGCTGTGAGCTCGGTACCAAATTCCAGAAAAGAGGCCGCGAAAGCGGCCTTTTTTCGT
TTTGGTCCAAGCCCGATGCGCCAGAGTTGTTTCTGAAACATGGCAAAGGTATCACTAGTCTTCGCGG
CCGCCATGCTGTCCAGGCAGGTAGATGACGACCATCAGGGACAGCTTCAAGGATCGCTCGCGGCTC
TTACCAGCCTAACTTCGATCATTGGACCGCTGATCGTCACGGCGATTTATGCCGCCTCGGCGAGCAC
ATGGAACGGGTTGGCATGGATTGTAGGCGCCGCCCTATACCTTGTCTGCCTCCCCGCGTTGCGTCGC
GGTGCATGGAGCCGGGCCACCTCGACCTGAATGGAAGCCGGCGGCACCTCGCTAACGGATTCACCA
CTCCAAGAATTGGAGCCAATCAATTCTTGCGGAGAACTGTGAATGCGCAAACCAACCCTTGGCAGA
ACATATCCATCGCGTCCGCCATCTCCAGCAGCCGCACGCGGCGCATCTCGGGCAGCGTTGGGTCCTG
GCCACGGGTGCGCATGATCGTGCTCCTGTCGTTGAGGACCCGGCTAGGCTGGCGGGGTTGCCTTAC
TGGTTAGCAGAATGAATCACCGATACGCGAGCGAACGTGAAGCGACTGCTGCTGCAAAACGTCTGC
GACCTGAGCAACAACATGAATGGTCATCGGTTTCCGTGTTTCGTAAAGTCTGGAAACGCGGAAGTC
AGCGCCCTGCACCATTATGTTCCGGATCTGCATCGCAGGATGCTGCTGGCTACCCTGTGGAACACCT
ACATCTGTATTAACGAAGCGCTGGCATTGACCCTGAGTGATTTTTCTCTGGTCCCGCCGCATCCATAC
CGCCAGTTGTTTACCCTCACAACGTTCCAGTAACCGGGCATGTTCATCATCAGTAACCCGTATCGTG
AGCATCCTCTCTCGTTTCATCGGTATCATTACCCCCATGAACAGAAATCCCCCTTACACGGAGGCATC
AGTGACCAAACAGGAAAAAACCGCCCTTAACATGGCCCGCTTTATCAGAAGCCAGACATTAACGCT
TCTGGAGAAACTCAACGAGCTGGACGCGGATGAACAGGCAGACATCTGTGAATCGCTTCACGACCA
CGCTGATGAGCTTTACCGCAGCTGCCTCGCGCGTTTCGGTGATGACGGTGAAAACCTCTGACACAT
GCAGCTCCCGGAGACGGTCACAGCTTGTCTGTAAGCGGATGCCGGGAGCAGACAAGCCCGTCAGG
GCGCGTCAGCGGGTGTTGGCGGGTGTCGGGGCGCAGCCATGACCCAGTCACGTAGCGATAGCGGA
GTGTATACTGGCTTAACTATGCGGCATCAGAGCAGATTGTACTGAGAGTGCACCGGTGTGAAATACC
GCACAGATGCGTAAGGAGAAAATACCGCATCAGGCGCTCTTCCGCTTCCTCGCTCACTGACTCGCTG
CGCTCGGTCGTTCGGCTGCGGCGAGCGGTATCAGCTCACTCAAAGGCGGTAATACGGTTATCCACA

GAATCAGGGGATAACGCAGGAAAGAACATGTGAGCAAAAGGCCAGCAAAAGGCCAGGAACCGTAA
AAAGGCCGCGTTGCTGGCGTTTTTCCATAGGCTCCGCCCCCCTGACGAGCATCACAAAAATCGACG
CTCAAGTCAGAGGTGGCGAAACCCGACAGGACTATAAAGATACCAGGCGTTTCCCCCTGGAAGCTC
CCTCGTGCGCTCTCCTGTTCCGACCCTGCCGCTTACCGGATACCTGTCCGCCTTTCTCCCTTCGGGA
AGCGTGGCGCTTTCTCATAGCTCACGCTGTAGGTATCTCAGTTCGGTGTAGGTCGTTCGCTCCAAGC
TGGGCTGTGTGCACGAACCCCCCGTTCAGCCCGACCGCTGCGCCTTATCCGGTAACTATCGTCTTGA
GTCCAACCCGGTAAGACACGACTTATCGCCACTGGCAGCAGCCACTGGTAACAGGATTAGCAGAGC
GAGGTATGTAGGCGGTGCTACAGAGTTCTTGAAGTGGTGGCCTAACTACGGCTACACTAGAAGGACA
GTATTTGGTATCTGCGCTCTGCTGAAGCCAGTTACCTTCGGAAAAAGAGTTGGTAGCTCTTGATCCG
GCAAACAAACCACCGCTGGTAGCGGTGGTTTTTTTGTTTGCAAGCAGCAGATTACGCGCAGAAAAA
AAGGATCTCAAGAAGATCCTTTGATCTTTTCTACGGGGTCTGACGCTCAGTGGAACGAAAACTCAC
GTTAAGGGATTTTGGTCATGAGATTATCAAAAAGGATCTTCACCTAGATCCTTTTAAATTAAAAATGA
AGTTTTAAATCAATCTAAAGTATATATGAGTAAACTTGGTCTGACAGTTACCAATGCTTAATCAGTGA
GGCACCTATCTCAGCGATCTGTCTATTTCGTTCATCCATAGTTGCCTGACTCCCCGTCG

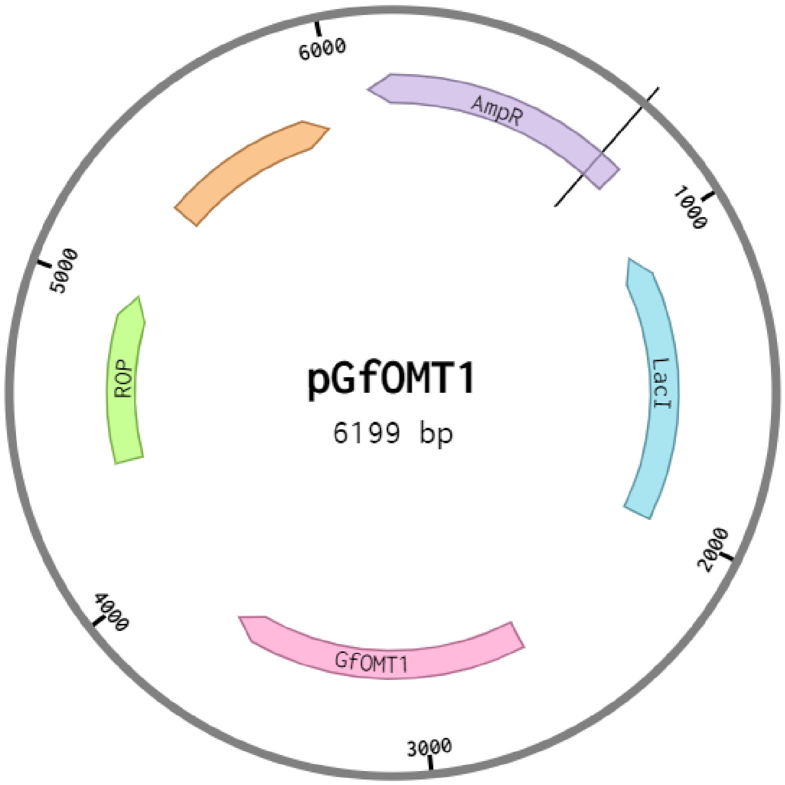


**> pThpR**

CCTGACACTCAGTTCCGGGTAGGCAGTTCGCTCCAAGCTGGACTGTATGCACGAACCCCCCGTTCA
GTCCGACCGCTGCGCCTTATCCGGTAACTATCGTCTTGAGTCCAACCCGGAAAGACATGCAAAAGC
ACCACTGGCAGCAGCCACTGGTAATTGATTTAGAGGAGTTAGTCTTGAAGTCATGCGCCGGTTAAGG
CTAAACTGAAAGGACAAGTTTTGGTGACTGCGCTCCTCCAAGCCAGTTACCTCGGTTCAAAGAGTT
GGTAGCTCAGAGAACCTTCGAAAAACCGCCCTGCAAGGCGGTTTTTTCGTTTTCAGAGCAAGAGAT
TACGCGCAGACCAAAACGATCTCAAGAAGATCATCTTATTAATCAGATAAAATATTTCTAGATTTCAG
TGCAATTTATCTCTTCAAATGTAGCACCTGAAGTCAGCCCCATACGATATAAGTTGTAATTCTCATGTT
AGTCATGCCCCGCGCCCACCGGAAGGAGCTGACTGGGTTGAAGGCTCTCAAGGGCATCGGTCGAG
ATCCCGGTGCCTAATGAGTGAGCTAACTTACATTAATTGCGTTGCGCTCACTGCCCGCTTTCCAGTCG
GGAAACCTGTCGTGCCAGCTGCATTAATGAATCGGCCAACGCGCGGGGAGAGGCGGTTTGCGTATT
GGGCGCCAGGGTGGTTTTTCTTTTCACCAGTGAGACGGGCAACAGCTGATTGCCCTTCACCGCCTG
GCCCTGAGAGAGTTGCAGCAAGCGGTCCACGCTGGTTTGCCCCAGCAGGCGAAAATCCTGTTTGAT
GGTGGTTAACGGCGGGATATAACATGAGCTATCTTCGGTATCGTCGTATCCCACTACCGAGATGTCCG
CACCAACGCGCAGCCCGGACTCGGTAATGGCGCGCATTGCGCCCAGCGCCATCTGATCGTTGGCAA
CCAGCATCGCAGTGGGAACGATGCCCTCATTCAGCATTTGCATGGTTTGTTGAAAACCGGACATGGC

ACTCCAGTCGCCTTCCCGTTCCGCTATCGGCTGAATTTGATTGCGAGTGAGATATTTATGCCAGCCAG
CCAGACGCAGACGCGCCGAGACAGAACTTAATGGGCCCGCTAACAGCGCGATTTGCTGGTGACCCA
ATGCGACCAGATGCTCCACGCCCAGTCGCGTACCATCTTCATGGGAGAAAATAATACTGTTGATGGG
TGTCTGGTCAGAGACATCAAGAAATAACGCCGGAACATTAGTGCAGGCAGCTTCCACAGCAATGGC
ATCCTGGTCATCCAGCGGATAGTTAATGATCAGCCCACTGACGCGTTGCGCGAGAAGATTGTGCACC
GCCGCTTTACAGGCTTCGACGCCGCTTCGTTCTACCATCGACACCACCACGCTGGCACCCAGTTGAT
CGGCGCGAGATTTAATCGCCGCGACAATTTGCGACGGCGCGTGCAGGGCCAGACTGGAGGTGGCA
ACGCCAATCAGCAACGACTGTTTGCCCGCCAGTTGTTGTGCCACGCGGTTGGGAATGTAATTCAGC
TCCGCCATCGCCGCTTCCACTTTTTCCCGCGTTTTCGCAGAAACGTGGCTGGCCTGGTTCACCACGC
GGGAAACGGTCTGATAAGAGACACCGGCATACTCTGCGACATCGTATAACGTTACTGGTTTCACATT
CACCACCCTGAATTGACTCTCTTCCGGGCGCTATCATGCCATACCGCGAAAGGTTTTGCGCCATTCG
ATGGTGTCCGGGATCTCGACGCTCTCCCTTATGCGACGCGGCCGCGGCATCAGAGCAGATTGTACTG
TGTCCTCAATGGTTCGTTGTGATGGCGGTAGGAATGTAATCGTTAATCCGCAAATAACGTAAAAACC
CGCTTCGGCGGGTTTTTTTATGGGGGGAGTTTAGGGAAAGAGCATTTGTCATCCCGTTGAATATGGC
TCGCATCTTATCGAGCATACTATCACGTCGGCGACCACTAGTCAGTTAACGCAAGGGCATGGGCTGT
CGACCTTTGAAAAGTACCTTGACGGCGTATCTTTGCTTTCTATAATGAGTGCTTACTCACTCATACAA
TAGTCAGTCATAAGTCTGGGCTAAGCCCACTGATGAGTCGCTGAAATGCGACGAAACTTATGACCTC
TACAAATAATTTTGTTTAACGTAAACCTCCGGGTTAATAAGGAGTAATTATGGCATCCAAGGGCGAG
GAGCTCTTTACTGGCGTAGTACCAATTCTCGTAGAGCTCGATGGCGATGTAAATGGCCATAAGTTTTC
CGTACGCGGCGAGGGCGAGGGCGATGCAACTAACGGCAAGCTCACTCTCAAGTTTATTTGTACTACT
GGCAAGCTCCCAGTACCATGGCCAACTCTCGTAACTACTCTGACCTATGGCGTACAATGTTTTTCCCG
CTATCCAGATCACATGAAGCAACATGATTTTTTTAAGTCCGCAATGCCAGAGGGCTATGTACAAGAG
CGCACTATTAGCTTTAAGGATGATGGCACCTATAAGACTCGCGCAGAGGTAAAGTTTGAGGGCGATA
CTCTCGTAAATCGCATTGAGCTCAAGGGCATTGATTTTAAGGAGGATGGCAATATTCTCGGCCATAA
GCTGGAGTATAATTTCAATTCCCATAATGTATATATTACCGCAGATAAGCAAAAGAATGGCATTAAGG
CGAATTTTAAGATTCGCCATAATGTGGAGGATGGCTCCGTACAACTCGCAGATCATTATCAACAAAAT
ACTCCAATTGGCGATGGCCCAGTACTCCTCCCAGATAATCATTATCTCTCCACTCAATCCGTGCTCTC
CAAAGATCCAAATGAGAAGCGCGATCACATGGTACTCCTGGAGTTTGTAACTGCAGCAGGCATTACT
CATGGCATGGATGAGCTCTATAAGCTCGAGCACCACCACCACCACCACTGATAATCCAAACCTGTTA
TATGTTAGCTGAGACTAGTTGGAAGTGTGGCTGTCCTCAAGCGTTTTAGTTCGTCGGTCAGTTTCAC
CTGATTTACGTAAAAACCCGCTTCGGCGGGTTTTTGCTTTTGGAGGGGCAGAAAGATGAATGACTG
TCGGCCATTCGATGGTGTCGGGTAGCATAACCCCTTGTGATACCTTTGCCATGTTTCAGAAACAACT
CTGGCGCATCGGGCTTGGACCAAAACGAAAAAAGGCCGCTTTCGCGGCCTCTTTTCTGGAATTTGG
TACCGAGCTCACAGCGGCGATAGTCAGATAGCTAGACCGTATGTTACCGAGCCTGCACTCCTCGAAT
TCTTAGGATTATTACTGCTCTTCGCGCGTAAGTGCGCGCCACATAGCCTCGAAGCCCAACGCAATGT
ACTCACCAGCGCGAGCCGGGTTGCGCGCAGCGAAATCCATAGTCGTCTCAGCAAGCGCCAAGAAC
AACCCGTCGCCGAAGGCGCGGTACTCGTCGGACATAAACACCATAAGAACCTCACGGTGGACGATG
TCGCGTAACTCCGGGAACATATCATCCGCGCGTTGTTCGGTCTCCTTCGTCAACTTTTCAGAAACCG
CCAACTGACGTATGGCACGATGGCGAGCTGGGTGGTTCAATCCCCAGCTAATGATACTGTTCCAGAG
AAAACGCGACATCATCTTAGCGTCAGTAATAGAACGATCCAATTCCATGATGGATGATTGGCACCAG
TCCTGGAACAAATGTAAGTAAAGGGTGTTGATCAACTCATCTTTCGTTGCGAAATAGCGGAACAAC
GTCCCTTCCGCAACTCCCGCATTGCGTGCAATTACAGCGGTACTAGCGGCAATGCCTGATTGCGCGA
TGGCTTGAGTTGCCGCTTCAAGCAATGCCTGCTTTTTGTCCTCAGACTTTGGGCGAGCAACCATATA
CTAACCTCCTTCTGATACGTGGTTCCGTTAAACAAAATTATTTGTAGAGGCCCCATTTCGTCCTTTTG
GACTCATCAGGGGTGGTACACACCACCCCTATGGGGCTCGTAATTGCTAGCATAATCCCTAGGACTGA
GCTAGCTATCAGGGTACTTTTCAAAGGTCGACAGCCCATGCCCTTGCGTTCGGCAGGTGTACAATGA
TACGAGGTAATGAAGATGAAGTCCATACAATCGATAGATTGGGACCAAAACGAAAAAAGGGGAGCG
GTTTCCCGCTCCCCTCTTTTCTGGAATTTGGTACCGAGTCGCACCTGATTGCCCGACATTATCGCACG
GTGTCTCATCTCTGATAACGCATATTGTCGTTAGAACTCGGCGCGGCCGCTCACACTGCTTCCGGTA
GTCAATAAACCGGTAAACCAGCAATAGACATAAGCGGCTATTTAACGACCCTGCCCTGAACCGACGA
CCGGGTCGAATTTGCTTTCGAATTTCTGCCATTCATCCGCTTATTATCACTTATTCAGGCGTAGCAAC
CAGGCGTTTAAGGGCACCAATAACTGCCTTAAAAAAATTAGAAAAACTCATCGAGCATCAAATGAA
ACTGCAATTTATTCATATCAGGATTATCAATACCATATTTTTGAAAAAGCCGTTTCTGTAATGAAGGAG
AAAACTCACCGAGGCAGTTCCATAGGATGGCAAGATCCTGGTATCGGTCTGCGATTCCGACTCGTCC
AACATCAATACAACCTATTAATTTCCCCTCGTCAAAAATAAGGTTATCAAGTGAGAAATCACCATGAG
TGACGACTGAATCCGGTGAGAATGGCAAAAGTTTATGCATTTCTTTCCAGACTTGTTCAACAGGCCA
GCCATTACGCTCGTCATCAAAATCACTCGCATCAACCAAACCGTTATTCATTCGTGATTGCGCCTGAG
CGAGACGAAATACGCGGTCGCTGTTAAAAGGACAATTACAAACAGGAATCGAATGCAACCGGCGCA
GGAACACTGCCAGCGCATCAACAATATTTTCACCTGAATCAGGATATTCTTCTAATACCTGGAATGCT

GTTTTCCCGGGGATCGCAGTGGTGAGTAACCATGCATCATCAGGAGTACGGATAAAATGCTTGATGG
TCGGAAGAGGCATAAATTCCGTCAGCCAGTTTAGTCTGACCATCTCATCTGTAACATCATTGGCAAC
GCTACCTTTGCCATGTTTCAGAAACAACTCTGGCGCATCGGGCTTCCCATACAATCGATAGATTGTCG
CACCTGATTGCCCGACATTATCGCGAGCCCATTTATACCCATATAAATCAGCATCCATGTTGGAATTTA
ATCGCGGCCTAGAGCAAGACGTTTCCCGTTGAATATGGCTCATTTTAGCTTCCTTAGCTCCTGAAAAT
CTCGATAACTCAAAAAATACGCCCGGTAGTGATCTTATTTCATTATGGTGAAAGTTGGAACCTCTTAC
GTGCCGATCACGTCTCATTTTCGCCAAAGTTGGCCAGGGCTTCCCGGTATCAACAGGGACACCAGG
ATTTATTTATNNTGCGAAGTGATCTTCCGTCACAGGTATTTATTCGGCGCAAAGTGCGTCGGGTGATG
CTGCCAACTTACTGATTTAGTGTATGATGGTGTTTTTGAGGTGCTCCAGTGGCTTCTGTTTCTATCAG
CTGTCCCTCCTGTTCAGCTACTGACGGGGTGGTGCGTAACGGCAAAAGCACCGCCGGACATCAGCG
CTAGCGGAGTGTATACTGGCTTACTATGTTGGCACTGATGAGGGTGTCAGTGAAGTGCTTCATGTGG
CAGGAGAAAAAAGGCTGCACCGGTGCGTCAGCAGAATATGTGATACAGGATATATTCCGCTTCCTCG
CTCACTGACTCGCTACGCTCGGTCGTTCGACTGCGGCGAGCGGAAATGGCTTACGAACGGGGCGGA
GATTTCCTGGAAGATGCCAGGAAGATACTTAACAGGGAAGTGAGAGGGCCGCGGCAAAGCCGTTTT
TCCATAGGCTCCGCCCCCCTGACAAGCATCACGAAATCTGACGCTCAAATCAGTGGTGGCGAAACC
CGACAGGACTATAAAGATACCAGGCGTTTCCCCCTGGCGGCTCCCTCGTGCGCTCTCCTGTTCCTGC
CTTTCGGTTTACCGGTGTCATTCCGCTGTTATGGCCGCGTTTGTCTCATTCCACG

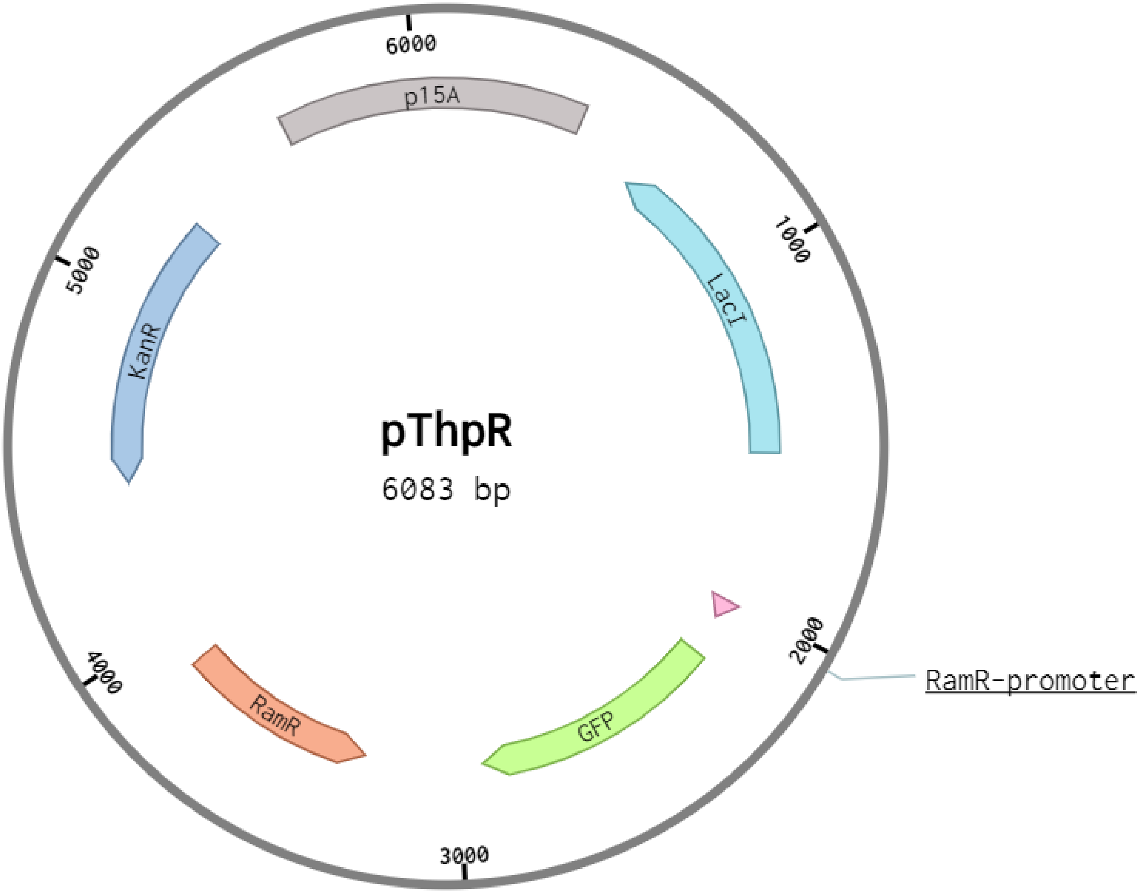


**Supplementary Table 4**. Sequences and graphical maps of representative plasmids.
